# Pan-cancer study on transition of signaling systems from primary to metastatic tumors

**DOI:** 10.1101/2024.11.11.619269

**Authors:** Wenjia Zhou, Junhua Zhang

## Abstract

Metastatic progression is responsible for the majority of cancer-related deaths. A better understanding of the underlying molecular mechanisms driving metastasis therefore remains of utmost importance. Here we take efforts to excavate mutant driver factors of metastases in the level of signaling pathways. We introduce two models, EntCDP and ModSDP, to detect the similarity and specificity of pathways in a pan-cancer context. The simulation studies confirm their feasibility and identification accuracy. Using mutation profiles of 17 cancer types with primary and metastatic cohorts from MSK-IMPACT, we apply the two models to investigate the transition of signaling systems from primary to metastatic tumors mainly from five perspectives. 1) We initially perform a comprehensive comparison of 15 primary-metastatic cancer pairs. Nearly all shared common driver genes or pathways, while specific driver gene sets of primary or/and metastatic tumors can be seen for almost each pair, except for breast cancer. 2) We use ModSDP to identify the specificity of metastases with the same primary site and highlight calcium signaling in breast cancer with bone metastasis, FoxO signaling for melanoma/lung lymph node metastasis, etc. 3) Oppositely, we use EntCDP to identify common factors yielding three frequently metastatic sites (liver, lung and brain), and recognized cushing syndrome, microRNAs in cancer and MAPK signaling pathway, separately. 4) Regarding high-tropism metastatic patterns, we investigated the relationship between the metastatic tumor and primary tumors in both locations. In the typical pattern of colorectal cancer metastasis to the liver, we detected hepatic signals in primary colorectal cancer. 5) Finally, we focused on the common and specific characteristics of relevant cancer types, such as gender-related cancers and gastrointestinal cancers. Study on gynecologic tumors suggest endocrine hormone change and virus infection as their common risk factors, while for male we highlight hedgehog signaling and EPH-related receptors as invasive potential of prostate cancer. Multiple interesting findings revealed by this study may be helpful for the understanding of the extent of signal changes during tumor metastasis. We expect that this study will provide a valuable resource for transforming our knowledge about signals in cancer metastasis into alternative clinical practice for advanced patients.

## Introduction

Cancer metastasis is a very complicated process, in which cancer cells detach from a primary tumor to enter circulation, endure pressure in blood vessels, at last colonize in a secondary tissue and grow in a hostile environment [1, 2]. Although most cancer deaths are due to metastatic spread, the molecular mechanisms of cancer metastasis remain largely unknown.

Many investigations of cancer progression and metastasis have been made on genomic characteristics in several aspects. For example, there were studies to characterize the overall genomic landscapes of metastases [3–5] or to investigate metastatic genomics complexity by reconstructing tumor evolution [6–8]. Furthermore, some studies tried to identify associations between genomic features and certain patterns of metastatic cancers [9].

We noticed that a lot of previous efforts on cancer metastasis focused on tumor specific cohorts, such as breast carcinoma [4,10], lung cancer [8], hepatocellular carcinoma [5], gastric cancer [11], ovarian high-grade serous carcinoma [12], et al. In recent years, several large-scale pan-cancer studies appeared [13–16], in which the authors tried to characterize genomic differences between primary and metastatic cancers, or to investigate the contribution of genomic changes to extraordinarily metastatic capacities of cancer cells. Although these efforts have yielded valuable insights into the genomic characteristics for metastatic cancers, comprehensive investigations on roles of signaling pathways for complex metastatic processes based on pan-cancer cohorts are seldom.

It is known that although many cancer cells are invasive, the majority will not survive during the metastasis process, and only a few will successfully colonize distant organs [1,2]. In fact, cancer metastasis is regulated by signaling pathways which govern cytoskeletal dynamics, turnover of cell–matrix, cell-to-cell junctions, and migration into adjacent tissues [17]. Coorperative signal systems constantly maintain cancer cell growth, proliferation, survival and reduce apoptosis. Therefore, it is very critical to deeply investigate the crucial roles of driver signaling pathways in cancer metastasis.

Among the classical methods known for identifying driver signaling pathways, de novo discovery of signal transduction pathways sheds light on the particular pattern of mutations and implies promising targets for clinical treatments [18, 19]. Instead of using information of gene-gene interaction, a driver pathway can be searched from a cloud of mutational genes through its biologically special characteristics: high coverage and exclusivity [18]. Based on these properties, the ComMDP and SpeMDP models were developed, at the pan-cancer level, to discover common gene sets across distinct tumor types and specific gene sets of certain one or multiple tumor types compared to other types, respectively. Comparative analysis naturally paves the way for cancer research and personalized clinical medication: commonality can inform the adoption of a consistent drug regimen across different cancers, while specificity can facilitate targeted drug delivery methods for individual cancer types. However, ComMDP aims to maximize the sum of weights of different cancers, but in practice, it is not always feasible to ensure that the identified gene sets will possess both properties in each cancer. One the other hand, SpeMDP overly emphasizes the contribution of coverage to the weight function.

To compensate for the shortcomings of ComMDP and SpeMDP and identify signaling pathways that are highly likely to be associated with cancer, we proposed two improved models called EntCDP (Entropy-based Common Driver Pathway) and ModSDP (Modified Specific Driver Pathway). We tested these two models on simulation datasets to show their advantage over ComMDP and SpeMDP. The EntCDP utilizes information entropy to measure the coverage of the submatrix across different cancers, ensuring that the weight of each cancer is as large as possible. The ModSDP partially addresses the inadequate consideration of exclusivity in the SpeMDP. Existing models have rarely been applied to the study of metastatic cancers, therefore, we apply our models to excavate mutant driver factors of metastases, and we expect such application will be very promising.

The biological data we collected are from a large-scale prospective clinical sequencing using a thorough assay called MSK-IMPCT, which was established to reveal massive tumor genomic spectrum of alternations of paired primary and metastatic cancers [13]. For mutation profiles of 17 cancer types with primary and metastatic cohorts of ∼ 10,000 patients, we investigated transition of signaling systems from primary to metastatic tumors by EntCDP and ModSDP mainly from five perspectives. 1) For an overall comparison of the 15 paired primary and metastatic cancers, almost each pair of them (14/15) has common driver gene sets or driver pathways (except non-melanoma skin cancer), and meanwhile, specific driver gene sets (driver pathways) of primary or/and metastatic tumors can be seen for almost each pair (14/15, except breast carcinoma). 2) For metastases with the same origin (such as brain, lung, liver metastases from melanoma), we focused on investigating which specific features may result in their different destinations. Among them, we highlight several signaling pathways: (a) the PI3K-Akt signaling pathway – it may contribute to many metastases, such as breast liver and lung metastases, as well as melanoma liver, lung, and lymph node metastases; (b) the FoxO signaling – it may contribute to melanoma lymph node metastasis and lung lymph node metastasis; (c) proteoglycans in cancer, the homologous recombination and the calcium signaling – they may be related to breast liver metastasis, breast lung metastasis and breast bone metastasis, respectively; (d) the JAK-STAT signaling and the Ras signaling – they may contribute to melanoma lung metastasis; (e) and the AMPK signaling – it may be related to melanoma liver metastasis. 3) For the same metastasis originating from different primary cancers (such as non-small-cell lung cancer, breast carcinoma, melanoma, colorectal cancer, pancreatic cancer, and prostate cancer metastasizing to liver), we paid more attention to exploring which commonalities to be involved in the metastatic processes with the same destination. We found cushing syndrome signaling may be a common factor making liver the frequent site for cancer metastases. MicroRNAs in cancer and MAPK signaling may be common driving factors for lung metastases and brain metastases, respectively. 4) For some typical patterns with high metastatic tropisms (such as liver metastasis from colorectal cancer), triple relations were investigated (such as commonalities or specificities among primary colorectal cancer, colorectal liver metastasis, and primary hepatocellular carcinoma). We suppose virus-associated pathways might be precursors of liver metastasis tendency in colorectal cancer cells. 5) For certain typical types of cancers (such as three gynecological cancers: breast carcinoma, ovarian, and endometrial cancer), we deeply investigated the complex common and specific characteristics among the primary and metastatic tumors, respectively. Our results showed that gynecologic tumors retain more of the characteristics of primary tumors after becoming metastases than male tumors. In summary, our research enhances the understanding of cancer metastasis mechanisms and offers valuable insights that may inform effective treatments, particularly for advanced-stage cancers.

## Results

In this study we introduced two optimization models, EntCDP and ModSDP, to detect the similarity and specificity of pathways in a pan-cancer context. A detailed description of the two models can be seen in the **Materials and Methods** section. Then we used simulation data to test the identification ability of the models, and used mutation profiles of 17 cancer types with primary and metastatic cohorts from MSK-IMPACT to investigate the transition of signaling systems from primary to metastatic tumors mainly from five perspectives based on the two models (Fig. 1).

**Figure 1.**
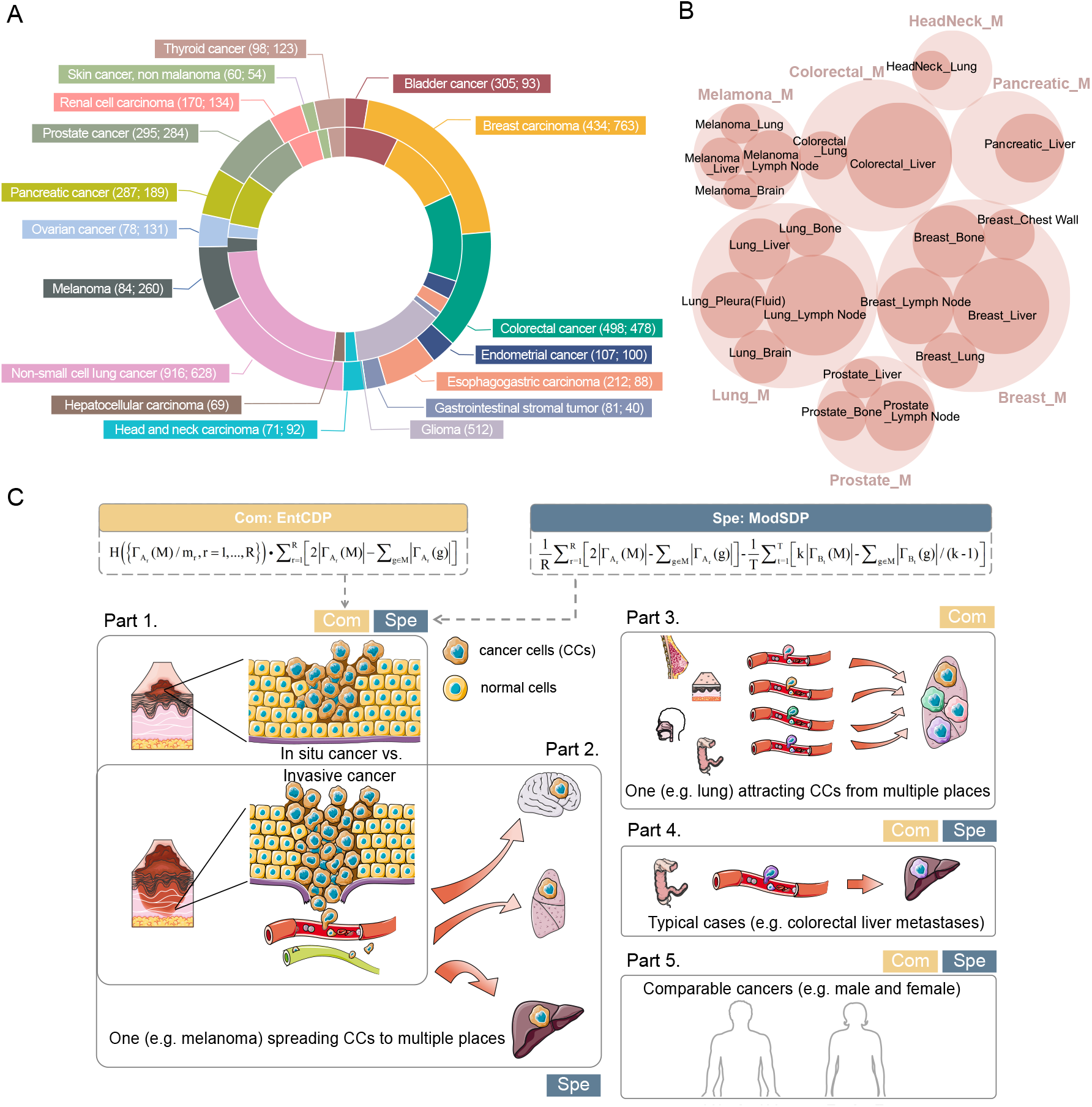
Overview of datasets and workflow. (A) Distribution of cancer types classified as either primary or metastatic tumors in the MSK-IMPACT cohorts. The first number in the parenthesis behind each cancer name is the number of its primary cases, and the latter is for metastases, corresponding to the inner and the outer ring, respectively. (B) Proportion of metastatic cancer sub-cohorts with the same origin. See **Materials and Methods** for naming rules. (C) Framework for our pathway analysis. The labels Com and Spe in the hatched area boxes indicate the EntCDP or/and ModSDP models used in this part. Part 1: Overall comparison of the paired primary and metastatic tumors; Part 2: for metastases with the same origin (such as melanoma), we investigate which specific features may result in their different destinations; Part 3: for the same metastasis originating from different primary cancers, we explore which commonalities to be involved in the metastatic processes with the same destination (such as lung); Part 4: for some typical patterns with high metastatic tropisms (such as Colorectal Liver), triple relations were investigated (such as commonalities or specificities among Colorectal P, Colorectal Liver and Liver P); Part 5: for certain typical types of cancer (such as three gynecological cancers), we deeply investigated the complex common and specific characteristics among the primary and metastatic tumors, respectively. CCs: cancer cells.

For the description of the simulation data and the biological data (Fig. 1A, B) used in this study please also refer to the **Materials and Methods** section.

### EntCDP can efficiently and accurately identify gene sets that are uniformly covered across various cancers

We first construct three intuitive simulation examples to illustrate the advantages of EntCDP over Com-MDP (Supplementary Text 1.3: Sim1 ∼ 3). In all three cases, two modules are embedded in each mutation matrix of two or three cancers. Additionally, module 2 has more uniform coverage than module in different cancers, therefore has larger biological significance.

In Sim1, we apply ComMDP and EntCDP to identify gene sets with three genes for two cancers (Fig. 2A). Using ComMDP, the objective function value of module 1 (17) is greater than the value of module 2 (16), so that ComMDP prefers module 1 than 2. Using EntCDP, since 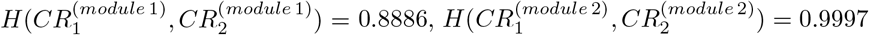, so the objective function values of module 1 and module 2 are calculated as 17 × 0.8886 = 15.1062 and 16 × 0.9997 = 15.9952, respectively. Therefore, EntCDP identifies module 2 from the two modules.

**Figure 2.**
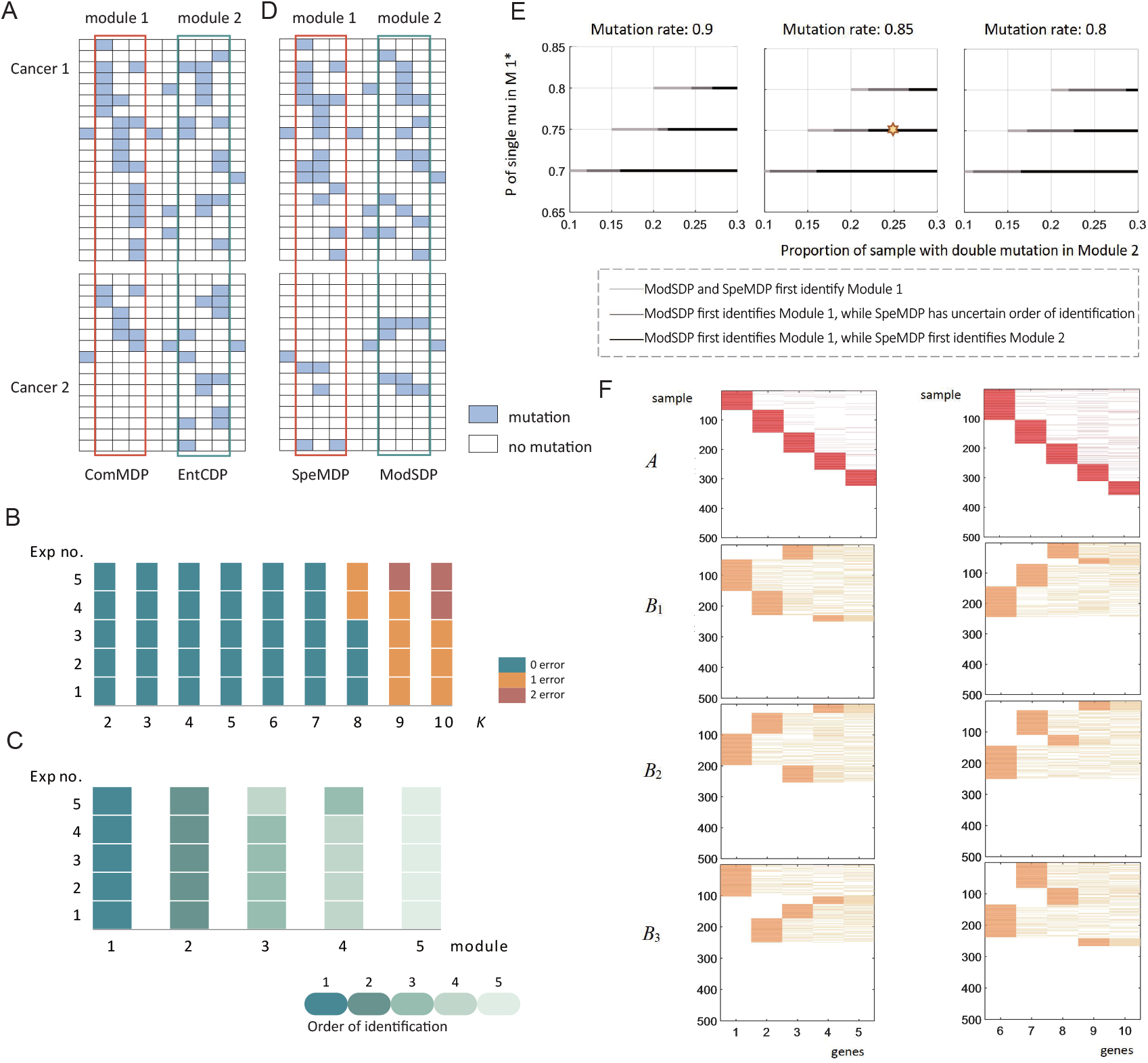
Results of the simulation results. (A) Schematic diagram of Sim1 to display the identification advantage of EntCDP over ComMDP, which simulates two cancers with uniform sample size. Each grid corresponds to a gene in a sample. Module 2 is more likely to be a common signaling pathway among all cancers than module 1. (B) Results of Sim4 for *k* = 2 to 10 with a constant mutation rate in case of uniform sample size. (C) Results of Sim7 and Sim8 for identifying five modules with *k* = 5 and decreasing proportion of mutated samples in case of uniform sample size. The green series indicates the full identification of the five genes in each module, and the color from dark to light implies the recognition order. (D) Schematic diagram of Sim9 to display the identification advantage of ModSDP over SpeMDP, which simulates two cancers. Module 2 is more likely to be a specific signaling pathway of cancer A relative to cancer B than module 1. (E) Results of Sim10. Comparison of order of identification with mutation rate at 0.9 (left), 0.85 (middle) and 0.8 (right). Black indicates that the SpeMDP identifies module 2 before module 1 and ModSDP identifies module 1 before module 2. Dark gray indicates that ModSDP identifies module 1 first, but not module 2, while SpeMDP sometimes first identifies module 1, sometimes identifies module 2. Light gray indicates that both models first identify module 1 in three simulation trials. *Proportion of samples with single mutation in module 1. (F) Heatmap of a case in Sim10 (yellow hexagon in E). The left and right side of mutation profiles illustrate module 1 and module 2 respectively in four simulated cancers.

Based on Sim1, we further explore scenarios where there are significant differences between two cancer samples (Supplementary Text 1.3: Sim2 and Supplementary Fig. S1A) and the identification of gene sets for three different cancers (Supplementary Text 1.3: Sim3 and Supplementary Fig. S1B). In all three instances, ComMDP prefers module 1 and EntCDP prefers module 2. It can be inferred that the ComMDP tends to identify gene sets with large coverage differences, whereas EntCDP mitigates the coverage rate differences of the gene set among various cancers by assigning the entropy-based weight to the gene set, which allows for the identification of gene sets that are more consistent with signaling pathways, even when the sample sizes differ significantly.

We conduct another five large-scale simulations (Supplementary Text 1.3: Sim4 ∼ 8) to evaluate the identification ability of EntCDP and each simulation is repeated five times in succession. Sim4 ∼ 6 test the accuracy of identifying modules with different number of genes (*k* = 2 ∼ 10) for three cancers. The sample sizes of the three cancers in Sim4 and Sim6 are relatively close, at 500, 550 and 600, whereas in Sim5, the sample sizes vary significantly, at 200, 500 and 800. Additionally, we added 2% and 4% ‘double mutation’ as disruptive factors of identification in Sim6. EntCDP can stably identify gene sets with the number of genes within 7, but occasionally misrecognize one or two genes outside the embedded modules and obtain non-significant cases when *k* = 8 ∼ 10 (Fig. 2B and Supplementary Fig. S2A). This is due to the very little non-exclusivity that the simulation itself adds to the ‘single mutation’ pattern (Supplementary Text 1.2). After the addition of ‘double mutation’, there are 3 wrong genes outside the embedded modules when *k* = 10 (Supplementary Fig. S2B), indicating that ‘double mutation’ will increase the instability of identification but this impact is considered acceptable.

Sim7 and Sim8 are designed to test the detection sequence of the EntCDP under different proportions of sample mutation rates for five cancers. The datasets in Sim8 have larger range of sample size, spanning from 200 to 1000, while those in Sim7 are from 300 to 500. For each embedded modules 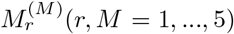 containing 6 genes in *A*_*r*_, the proportion of the sample selected to mutated is 0.75−*ϕ*(*r*)\**M, ϕ*(*r*) = 0.01*r* for both simulations. In the five repeated experiments, the model almost always identified the gene sets with the less obvious properties of the signaling pathways in order (Fig. 2C and Supplementary Fig. S2C), which indicates that the accuracy of the identification is not affected by the large-span sample size.

### ModSDP can efficiently and accurately identify gene sets with better exclusivity across various cancers

For ModSDP, we construct Sim9 and Sim10 to demonstrate the advantages of ModSDP over SpeMDP, and Sim11 ∼ 13 to evaluate the identification accuracy of ModSDP (Supplementary Text 1.4).

Like Sim1 ∼ 3, we give a schematic diagram of Sim 9, which simulates mutation matrices of two compared cancers with small samples. Obviously, module 2 has worse mutual exclusivity than module 1 in cancer 1, therefore it is less likely to be a signaling pathway. We apply SpeMDP and ModSDP to identify gene sets with three genes for two cancers (Fig. 2D). Using SpeMDP, the objective function value of module 1 (3 · 15 − 20 −(3 · 3 − 5) = 21) is greater than the value of module 2 (3 · 13 − 15 −(3 · 4 − 8) = 20), so that SpeMDP prefers module 1 than 2. Using ModSDP, the objective function values of module 1 and module 2 are calculated as 2 · 15 − 20 − (3 · 3 − 5)*/*2) = 8 and 2 · 13 − 15 − (3 · 4 − 8)*/*2 = 9, respectively. Therefore, ModSDP identifies module 2 from the two modules.

Further, we set up different proportions of mutation patterns for large-scale simulation data and study how the recognition results of the two models varies with these proportions (Sim10). Here, two zero matrices *A* and *B*_1_ ∼ *B*_3_ represent four types of cancer with column number of 900 and row number of 500. We embed *M* ^(*I*)^ and 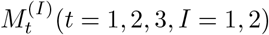 containing 5 genes in groups *A* and *B* respectively and implement mutation simulation with different combination of mutation rates. For group *A, M* ^(1)^ is carried out ‘single mutation’ and the proportion of samples to be mutated is 70%, 75% and 80% respectively, while *M* ^(2)^ is carried out combination of ‘single mutation’ and ‘double mutation’ and the proportion of samples harboring the first pattern is fixed at 60% but proportion in second pattern varying from 10% to 30%, with each increment of 2%. Additionally, we perform ‘double mutation’ for each module of group *B* on 60% of the samples. Each simulation is repeated three times under different parameters. In Sim10, *M* ^(1)^ are actually the specific signaling pathway for group *A* but not for group *B*. Differences in identification order between SpeMDP and ModSDP are plotted in Fig. 2E. Comparative to SpeMDP, we found that ModSDP may effectively judge the better features of gene sets and features better flexibility in identifying pathways. When we fix the mutation rate at 0.85, the proportion of samples with ‘single mutation’ and ‘double mutation’ at 0.75 and 0.25, respectively (yellow hexagon in Fig. 2E), heatmap drawn in this situation (Fig. 2F) reflects that SpeMDP tends to identify gene sets with slightly higher coverage but poor exclusivity, whereas ModSDP tends to identify gene sets with better exclusivity and slightly smaller coverage.

To confirm the recognition accuracy of the ModSDP, Sim11 tests it under different *k*, and Sim12 and Sim13 investigate the effects of different proportional mutation patterns. Both Sim11 and Sim12 simulate specific gene sets for one cancer (group *A*) relative to the other three cancers (group *B*), while Sim13 investigated two cancers versus five cancers. In Sim11, ModSDP can completely identify the first 8 modules, but occasionally has wrong identification when *k* = 9 and 10 (Supplementary Fig. S3). Results for Sim12 and Sim13 show its excellent identification ability in terms of different combinations of mutation rates, even for multiple cancer types (Supplementary Fig. S4).

In summary, we verify the effectiveness of the improvement in EntCDP and ModSDP on the simulation data. Next, we demonstrate the applicability of the two models on real cancer data and decipher the molecular mechanisms of cancer metastasis based on large-scale primary and metastatic cohorts from five perspectives (Fig. 1C).

### Overall comparison of primary and metastatic tumors

We first compared primary and metastatic cancers using EntCDP and ModSDP to understand the signal changes in an overall view for each of the 15 types of cancer with the paired data (Fig. 1A, Part1 and Data S1). For the compared tumors (for example, Breast P and Breast M), we used Breast P & Breast M to represent the common driver gene sets of Breast P and Breast M, and used Breast P / Breast M to represent the specific driver gene sets of Breast P versus Breast M, and vice versa. The enriched KEGG (Kyoto Encyclopedia of Genes and Genomes) pathways were displayed for the gene sets identified from the above two models (see **Materials and Methods**).

Our study shows that most metastatic tumors except non-melanoma skin cancer (14/15) have common driver gene sets or driver pathways with primary tumors (Fig. 3A, ‘Com P&M’ column). However, it might also be different between the two compared groups since new genetic mutations accumulate as cancerous cells grow clonally. Specific driver gene sets or driver pathways of primary versus metastatic tumors or vice versa can be seen in almost all cancer types mentioned here (14/15) (Fig. 3A, ‘Spe P/M’ or ‘Spe M/P’ columns). The specificity is sometimes more obvious on only one side. For example, the primary bladder cancer (Bladder P) has specific driver gene sets versus the metastatic bladder cancer (Bladder M), and Colorectal M has specific driver gene sets versus Colorectal P, but in each such case the reverse is not significant. Moreover, we observed that the primary and metastatic tumors for the breast carcinoma have more commonalities than specificities, because we identified common driver gene sets for Breast P and Breast M, but neither specific driver genes for Breast P versus Breast M nor for the reverse. However, the non-melanoma skin cancer has opposite properties, that is, both the primary and metastatic tumors have their own specific driver gene sets, but no common driver genes were identified (Fig. 3A).

**Figure 3.**
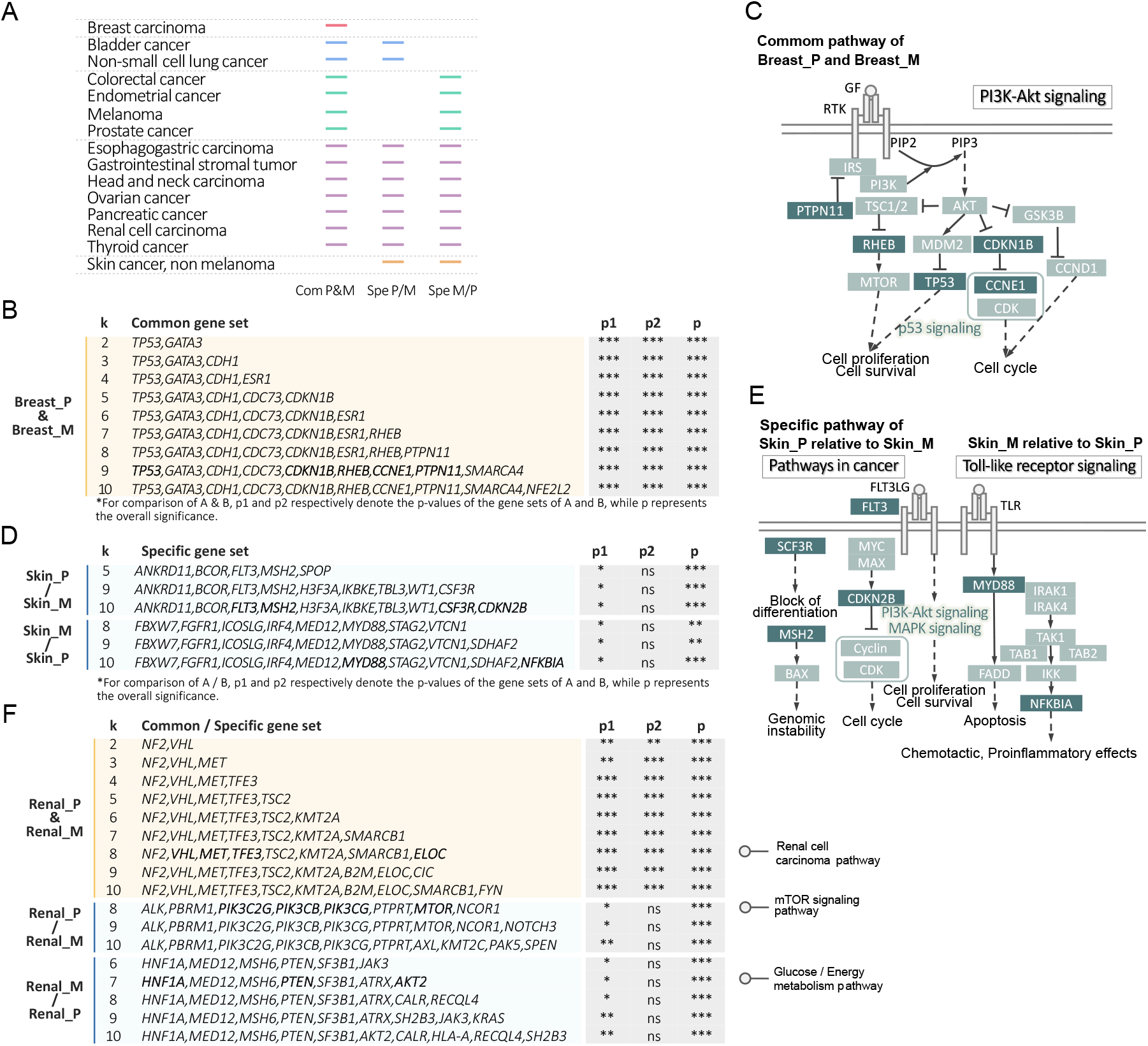
Comparison of driver genes and their enriched pathways between primary and metastatic tumors. (A) Label of common or specific results with computationally significant gene sets. (B, D, F) Results of EntCDP and ModSDP models for breast carcinoma (B), non-melanoma skin cancer (D), and renal cell carcinoma (F) as examples. The yellow area shows the common gene sets with different gene numbers (*k*) while blue denotes specific gene sets relative to each other. *: 0.01≤ *p<*0.05; **:0.001 ≤ *p<*0.01; ***: *p* ≤ 0.001. ns: not significant (for the meaning of *p*_1_, *p*_2_ and *p*, see **Materials and Methods**). (C, E) Common signaling pathways mapped for Breast_P and Breast_M, and specific pathways for Skin_P and Skin M at *k* = 9 and 10, respectively. The genes in the dark green background are those bold in (B) and (D), in other words, the identified driver genes by EntCDP or ModSDP.

In details, we first reported common driver gene sets between primary and metastatic tumors in patients with breast carcinoma (Fig. 3B). It is seen that two typical cancer genes *TP53* and *GATA3* [4,20] *are always identified with k* = 2 ∼ 10. As an example we looked at the case *k* = 9, where *TP53, RHEB, PTPN11, CDKN1B* and *CCNE1* were involved in the PI3K-AKT signaling (Fig. 3B, C and Data S3.3). It is known that nonreceptor protein tyrosine phosphatase SHP2, encoded by *PTPN11*, regulates breast cancer proliferation by mediating degradation of Cyclin D1 (*CCND1*) via the PI3K/AKT/GSK3*β* signaling pathway [21, 22]. No specific genes discovered for either group may indicate that breast cells are prone to behave accordantly within the mammary microenvironment, which benefits the use of medical drugs.

In contrast, non-melanoma skin cancer exhibits conspicuous distinctions with no detective commonality between the corresponding primary and metastatic cancers (Fig. 3A and Data S3.31-3.32). Compared with Skin M, Skin P is modulated by the pathways in cancer signaling since *CDKN2B, FLT3, CSF3R* and *MSH2* are identified as four key driver factors (Fig. 3D, E). The last two genes are also predicted to affect the activity of skin cell via the hematopoietic cell lineage, providing a new guideline for molecular targeted therapy of skin cancer. On the other hand, the Toll-like receptor (TLR) signaling (Fig. 3E), represented by *MYD88* and *NFKBIA* (identified as specific driver genes of Skin M), has been implicated in several forms of tumorigenesis via its inflammatory signaling downstream [23]. Our findings may promote TLR-based therapies for skin diseases like non-melanoma skin cancer [24].

There are seven types of cancer which have not only common driver gene sets but also specific sets for both primary and metastatic patients (such as head and neck carcinoma, ovarian cancer, renal cell carcinoma, pancreatic cancer, etc.) (Fig. 3A). For example, we surprisingly found renal cell carcinoma pathway (including *TFE3, VHL, ELOC* and *MET*) as the common signaling between samples with Renal P and Renal M (Fig. 3F, Renal P & Renal M, *k* = 8). We also discovered specific pathways, the mTOR pathway (including *PIK3C2G, PIK3CB, PIK3CG* and *MTOR*) (Fig. 3F, Renal P / Renal M, *k* = 8), and the glucose/energy metabolism pathway (including *HNF1A, AKT2* and *PTEN*) (Fig. 3F, Renal M / Renal P, *k* = 7) for the two groups, respectively (Data S3.28-3.30). For the latter case, *AKT2* and *PTEN* are also known to regulate the PD-L1 expression and PD-1 checkpoint pathway in cancer. It has been suggested that glucose deficiency could upregulate PD-L1 expression in renal cancer cell lines [25], so the two genes may pave the way between glucose metabolism and elevated PD-L1 expression, and direct the immune evasion and spread of renal carcinoma cells.

One-side specific enriched pathways are identified for six types of cancer (in addition to bladder cancer and colorectal cancer mentioned above, others include non-small cell lung cancer (NSCLC), melanoma, prostate cancer, etc.) (Fig. 3A). For example, we found the signaling by GPCR, typical of *CASP8, TERT, PTEN* and *TRAF2*, in metastatic melanoma, which was reported to modulate multiple crucial cellular processes during melanomagenesis, including proliferation and migration [26]. For NSCLC, Lung P turned out to be affected by *PDGFRA, RASA1, BRAF* and *EGFR* via the MAPK signaling pathway (Lung P / Lung M; Data S2, Table S1.8.2 and Data S3.16-3.17), in which *BRAF* and *EGFR* represent prevailing therapeutic targets for NSCLC [27].

Overall, the identified genes in common signaling pathways vary in different cancers and are known for their high frequency of mutations in corresponding cancers. Generally, the *PIK3CA*, Ras and KMT2 families are more likely to play an important role in signaling pathways related to migration and invasion.

### Abnormal signals may predict the seeding site of metastasis

As the cancer progresses to an advanced stage, it can usually metastasize to other different tissues. We wonder what drives tumor cells living in their lesions towards to different residences (Fig. 1C, Part 2). In this section, three types of cancers, breast carcinoma, melanoma and NSCLC, were chosen to investigate this question, mainly because their tumor cells demonstrate tenacious vitality during migration. We performed ModSDP to contrast different metastases from the same origin regarding their special signal manifestations relative to the primary cancer as well as pairwise comparisons within themselves.

### Different signals in tumor cells originating from breast

We start with breast carcinoma since advanced breast carcinoma is hard to treat and almost always fatal due to its high metastatic potential [28]. Among its secondary sites of metastasis, the metastasis lymph node predominates because it serves as the sign and prognosis of breast cancer [29]. Other breast cancer cells also preferentially metastasize to the lung, liver, chest wall and bone (Fig. 1B). Here, we mainly explored the five types of metastases from the breast (Fig. 4A).

**Figure 4.**
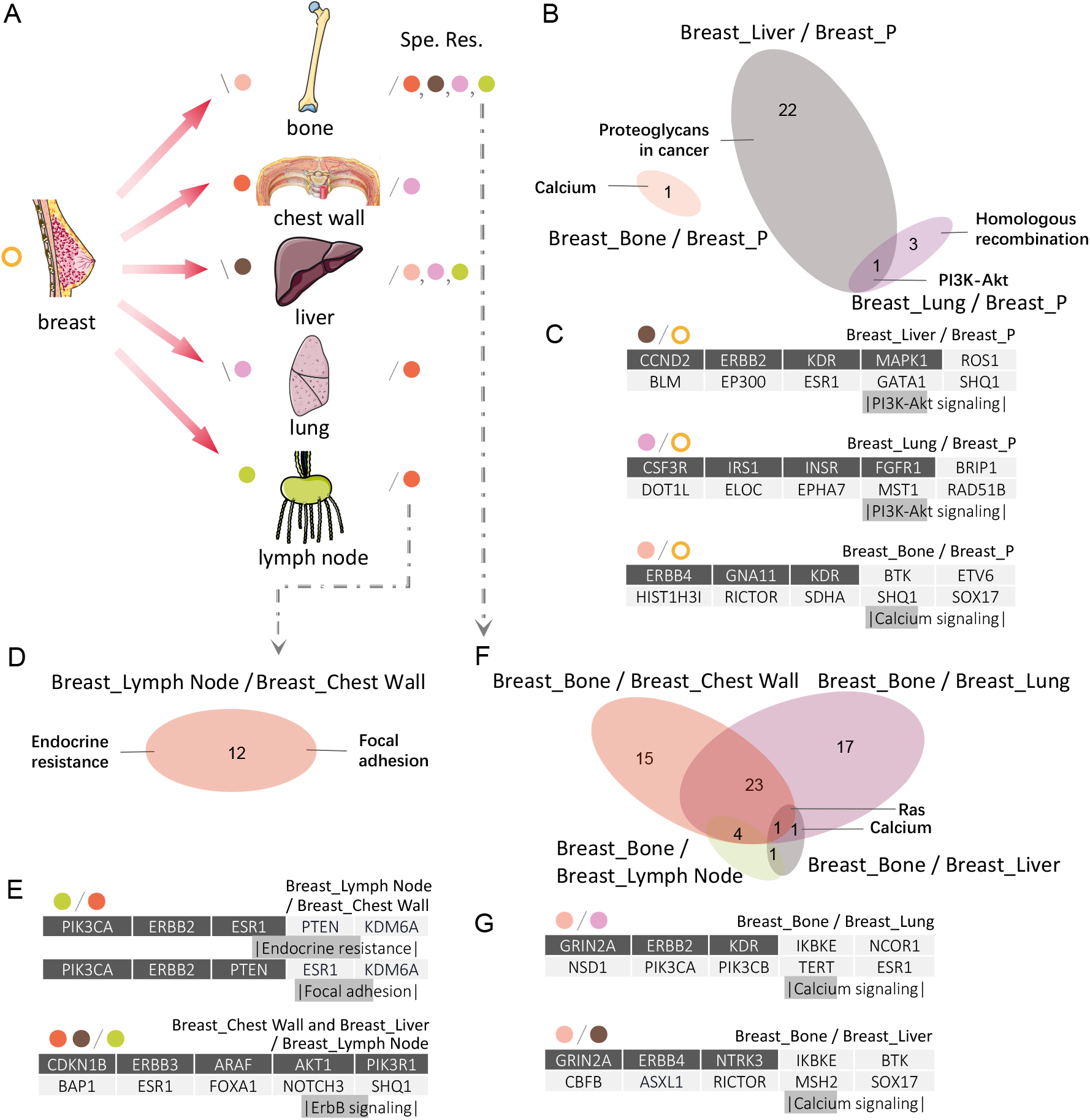
Specificity of different metastases from breast. (A) Five common sites of metastatic breast carcinoma and the results of ModSDP. Different colored dots in the left and middle columns denote primary breast carcinoma and five metastases from the breast. In the middle column, right slashes (\) on the left of dots indicate whether there are significant specific signaling pathways (SSSP) for each subtype of the metastases relative to the primary breast carcinoma. Left slashes (/) and colored dots on the right column exhibit that there are SSSP of the subtype before the left slash relative to each dot of subtypes separated by commas. For example, the four dots on the right side of the bone metastasis indicate that Breast Bone has SSSP compared to any other four subtypes. (B, D, F) Venn diagram displays the number of SSSP of some metastasis subtypes compared with primary tumors (B) or other subtypes (D, F). (C, E, G) Specific gene set of the subtype before left slash relative to the subtype behind it at some certain *k* and its enriched pathway corresponding to (B, D, F). The length of gray filling of each pathway is in the scale of the number of dark gray genes. Spe. Res. : Specific results.

New discoveries compensate for the lack of signal information about the specificity of Breast M mentioned in Part 1. The results reveal that Breast Liver has the most intensive variety of signals relative to Breast P. Its specific genes are enriched in 23 pathways, followed by Breast Lung and Breast Bone, with 4 and 1 enriched pathways, respectively (Fig. 4B and Data S4.1-4.3). Both Breast Liver and Breast Lung are specifically regulated by not only the same signaling pathways such as PI3K-Akt through different gene combinations, *CCND2, ERBB2, KDR, MAPK1* and *CSF3R, IRS1, INSR, FGFR1* (Fig. 4B, C), but also distinct pathways such as proteoglycans (PGs) in cancer and homologous recombination, respectively (Fig. 4B). Previous reports have investigated that activating the PI3K-Akt signaling pathway would promote breast cancer metastasizing toward certain organs including lung and liver [30, 31]. While PGs perform multiple functions in some stages of the metastatic cascade [32], homologous recombination is essential for preserving genome integrity, whose member, *RAD51B*, plays a critical role in breast cancer growth and metastasis [33]. Although specific genes of Breast_Bone relative to Breast_P are significantly enriched in only one pathway called calcium signaling (Fig. 4B) that induces metastatic cells to settle in bones during advanced-stage breast cancer [34], we emphasize the potential role of its three genes, *ERBB4, GNA11* and *KDR* (Fig. 4C and Table 1), in prompting cell migration from the breast.

**Table 1.**
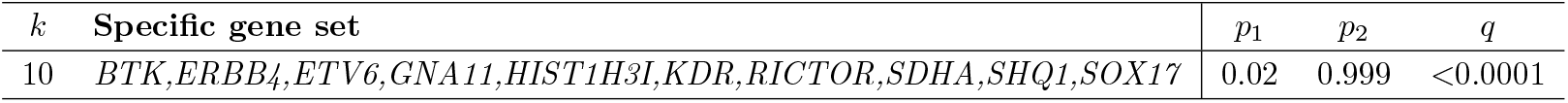
Breast_Bone specific mutated driver gene sets relative to Breast_P.

| *k* | Specific gene set | *p*<sub>1</sub> | *p*<sub>2</sub> | *q* |
| --- | --- | --- | --- | --- |
| 10 | *BTK,ERBB4,ETV6,GNA11,HIST1H3I,KDR,RICTOR,SDHA,SHQ1,SOX17* | 0.02 | 0.999 | <0.0001 |

ModSDP was also used to seek for unique features of metastases of breast carcinoma in pairs. The extraordinary relationship between lymph nodes and the breast stimulates us to primarily investigate the specificity of lymph nodes.

The results show that Breast_Lymph Node has specific genes relative to Brease_Chest Wall, while the inverse is not the case. However, Brease Chest Wall and Brease_Liver together exhibit specificity in relation to Breast_Lymph Node (Fig. 4E and Data S4.9, 4.14). For the former, specificity is embodied in breast cancer-related pathways such as endocrine resistance as well as metastasis-related pathways such as focal adhesion (Fig. 4D), the evidences of which are supported by outcomes in [35, 36]. For the latter, the ErbB signaling pathway was detected and represented by *CDKN1B, ERBB3, ARAF, AKT1* and *PIK3R1* (Fig. 4E). As one of the ErbB family members, *ERBB3* has been implicated in breast cancer [37]. Additionally, knockdown of *AKT1* promotes epithelial-mesenchymal transition (EMT) in breast cancer cells and increases the risk of metastasis [38].

The mutation profile of bone metastasis seems more complicated since it has significant specific pathways relative to any other four metastases from breast carcinoma, and the number of enriched pathways is relatively much larger among the five subtypes (Fig. 4A, F). Compared with Breast_Lung or Breast_Liver, the calcium signaling pathway was detected again as an essential pathway for Breast_Bone. Either genes *GRIN2A, ERBB2, KDR* or *GRIN2A, ERBB4, NTRK3*, may be used for combined targeted therapy (Fig. 4G). We highlight *GRIN2A* due to its frequent appearance in detected signaling pathways as a specific driver, such as the calcium pathway just mentioned and the Ras signaling pathway shared by three comparative cases: the specificities of Breast_Bone relative to Breast_Chest Wall, Breast_Lung, and Breast_Liver. (Fig. 4F, G and Data S4.5-4.6). Previous studies demonstrated that *GRIN2A* was associated with some other cancers. For example, *GRIN2A* mutations correlated with survivals in melanoma [39]. And recently, *GRIN2A* was screened as a candidate target against ovarian cancer by network pharmacology analysis [40]. Therefore, *GRIN2A* is likely to be a potential target gene for bone metastasis therapy in breast cancer.

### Different signals in tumor cells originating from melanoma

Melanoma is the malignant tumor with high rates of recurrence and metastasis [41]. Lymphoma occupies the largest number of samples with metastatic melanoma in our study, followed by the liver, lung and breast metastases (Fig. 1B). We investigated the heterogeneity of the four metastases in accordance with the analysis for the breast carcinoma (Supplementary Fig. S9A).

Despite metastatic features displayed in macroscopic comparison, more details lie in the specific genes of each subtype of the metastases relative to the primary tumor. After the completion of metastasis, accumulation of multiple gene mutations activates new tumorigenesis pathways. Melanoma Lymph Node catches our attentions because its specific genes relative to Melanoma_P are enriched in the largest number of pathways (Supplementary Fig. S9B) among the four types of metastases, which demonstrates greater risk of life. Typical genes in the PI3K-Akt signaling pathway, *AKT3, PTEN, PIK3CD, FLT1* and *FLT3*, were detected as specific risk factors for Melanoma_Lymph Node relative to Melanoma_P (Supplementary Fig. S9C and Data S2, Table S2.3.4). This pathway was also detected as being specific to both Melanoma_Lung and Melanoma_Liver relative to Melanoma_P (Supplementary Fig. S9B), which is consistent with the evidence that the regulation of the PI3K-Akt pathway would affect the proliferative and invasive capabilities of melanoma cells [42]. In addition, what impresses us is the respective identification of the JAK-STAT (including *PIM1, EP300* and *JAK3*) and AMPK (including *RPS6KB2, TSC2* and *IGF1R*) signaling pathways as novel hints of targeted treatments for Melanoma Lung and Melanoma Liver relative to Melanoma P (Supplementary Fig. S9B and Data S4.32-4.33). Aberrant JAK-STAT signaling has been identified to contribute to cancer progression and metastatic development [43]. The AMPK signaling negatively regulates the cellular energy status and is also involved in the process of tumor invasion and migration [44, 45]. These findings suggest a therapeutic strategy targeting these two pathways in the corresponding malignant tumors.

In regard to the comparison of the four metastases from melanoma, we discovered that each has specific gene sets relative to all or some of the others (Supplementary Fig. S9A). This characteristics is quite different from that of the metastases originating from breast carcinoma (Fig. 4A). We noticed that enriched pathways were detected for Melanoma_Lung relative to Melanoma_Liver or Melanoma_Lymph Node and one overlap pathway was the Ras signaling (Supplementary Fig. S9D). Given the successful treatment of *KRAS*^*G*12*C*^-mutated NSCLC [46], we suggest that *PAK1, IGF2* and *ABL1* or *NRAS, FLT4* and *RAC1* are essential for Ras-mediated metastatic potential of cancer cells migrating from melanoma to lung (Supplementary Fig. S9E). Additionally, lymph nodes showed greater heterogeneity since 14 common signaling pathways were identified in the context of comparison with the other three metastatic tumors (Supplementary Fig. S9F). Actually, *ERBB3, ATR, MAP2K1* and *PTEN* are frequent causative factors in all three cases and play an important role in cancer-related pathways such as cellular senescence, PI3K-Akt and FoxO signalings (Supplementary Fig. S9F). One recent research has examined the correlation between lymph node metastases and the PI3K-Akt signaling as well as its downstream target *FOXO4* [46], which supports our outcome and confirms the target potential of mutated genes in the PI3K-Akt or FoxO signaling.

The third cancer in this section, NSCLC, which frequently metastasizes in advanced cancer patients, we studied its five metastases, i.e., Lung_Bone, Lung_Brain, Lung_Liver, Lung_Lymph Node and Lung Pleura (Fluid) (Supplementary Fig. S6). We were also concerned about lymph node metastasis, which accounts for the largest number of samples among the five metastases. We found that the FoxO signaling seems to exert a powerful driving force for tumor cells to settle in the lymph nodes. Like Breast_Lymph Node, Lung_Lymph Node also diverges less from primary lung cancer (Fig. 4A and Supplementary Fig. S6A), implying that targeting primary alterations may also have lethal effect on metastatic tumor cells. See Supplementary Text 2.1 for more details about metastases originating from NSCLC.

### Three main metastatic lesions attract cancer cells to adapt to their tissue microenvironment

The human body is supposed to be a cell-based city in which some organs could turn out to be ‘tourist resorts’ of metastatic cells. An opposing perspective naturally turns to organs seemingly attracting circulating tumor cells for residence. Here, we chose three colonies - brain, liver and lung - due to their prevalence and previous reports [47]. These organs cover 3, 6, and 4 datasets, respectively, each associated with different primary lesions that exhibit a relatively high tendency to metastasize. To determine what makes the three organs frequent sites of cancer cell dissemination, we implemented EntCDP to detect whether there are some similarities between sub-cohorts with the same cancer metastases (Fig. 1C, Part 3).

### Liver as the preferred site for cancer metastases

Covering the largest number of metastatic origins in the three cancer types, hepatocellular carcinoma (HCC) presents its diversity and complexity. Its microenvironment facilitates cells of multiple types of cancer to establish secondary metastases. Here, we selected six types, colorectal cancer, breast carcinoma, pancreatic cancer, NSCLC, prostate cancer and melanoma, all with a relatively high propensity for the liver cancer (Fig. 1B and Fig. 5A).

**Figure 5.**
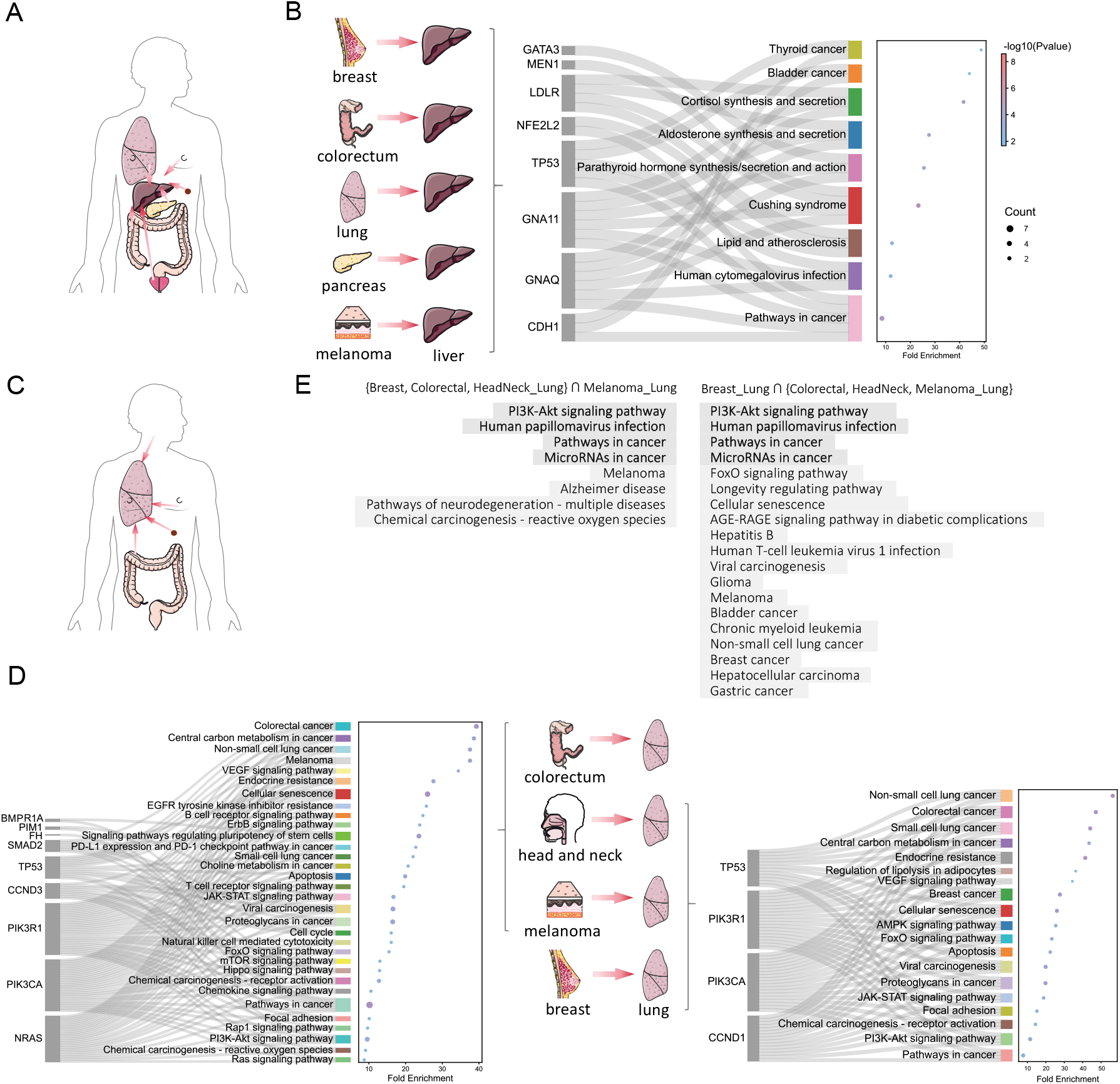
Commonality of cancer metastases of the same destination. (A, C) Schematic of the direction of HCC (A) or NSCLC (C) metastases. (B, D) Sankey plot of common genes of HCC (B) or NSCLC (D) metastases with some primary focis and the enriched pathways they are involved in. The dot plot shows the fold enrichment and the total number of genes in each enriched pathway. (E) Nominally common signaling pathways of the four NSCLC metastases obtained by intersecting common pathways of the three metastases with pathways of the remaining one.

Surprisingly, we identified *GNAQ, GNA11, LDLR* and *MEN1*, along with their highly correlated pathway, named cushing syndrome, which liver metastases from breast carcinoma, colorectal cancer, lung cancer, melanoma and pancreatic cancer have in common (Fig. 5B and Data S2, Table S3.1.1). Cushing syndrome is caused by excessive glucocorticoid secretion, and is often accompanied by liver carcinogenesis [48]. Mutations in these genes may serve as useful markers for predicting the metastatic potential of liver cancer.

Prostate Liver stands out due to its minimal commonalities with the other five subtypes, or even with any combination of three among them. After narrowing the comparisons down to two, we found a common signaling pathway called signaling by GPCR (including *SMAD2, TP53, STK11, MEN1, k* = 7) for the Pancreatic Liver, Colorectal Liver and Prostate Liver datasets (Table 2 and Data S5.2). Therapies for prostate cancer by manipulating GPCR-mediated pathways have recently been reported [49], which may be helpful in the prevention and cure of prostate cancer liver metastases. Furthermore, the PI3K-Akt signaling pathway (including *BRCA1, FGFR4, TP53, k* = 8) was identified for Colorectal Liver and Prostate Liver, and it was the only significant pathway by enrichment analysis (Data S2, Table S3.1.3 and Data S5.3).

**Table 2.**
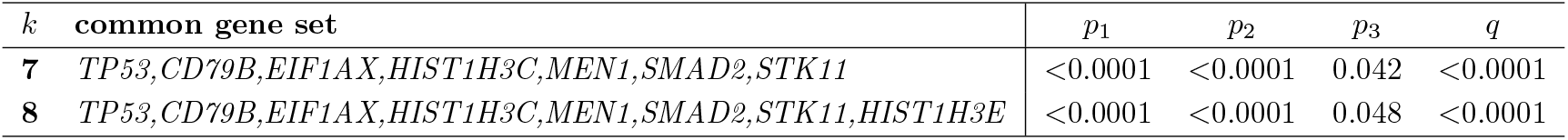
Significant common driver gene sets between Colorectal_Liver, Pancreatic_Liver and Prostate Liver.

For Breast Liver, NSCLC Liver and Melanoma Liver, we also detected some cancer-related pathways including not only some well-known signaling pathways, the pathways in cancer (including *MSH6, CDH1, GNAQ, GNA11, IGF1, ESR1* and *TP53*) and transcriptional misregulation in cancer (including *IGF1, TP53* and *MEN1*), but also growth hormone synthesis, secretion and action (including *GNAQ, GNA11* and *IGF1*) when *k* = 10 (Data S2, Table S3.1.4 and Data S5.4). The effects of growth hormone on hepatic diseases are not well understood, but it has been reported to influence insulin action in the liver [50]; therefore, dysregulation of this signaling may disturb the normal activities of liver cells and strengthen their communication with circulating tumor cells.

### Lung as the preferred site for cancer metastases

Lung metastases are also the lethal determinant in many cancers [51], and we chose those originated from breast carcinoma, colorectal cancer, head and neck cancer as well as melanoma. (Fig. 1B and Fig. 5C).

In the analysis of common signaling pathways, microRNAs in cancer, which has been widely reported to regulate the progression, invasion, metastasis and drug resistance of lung cancer [52–54], appears in every enrichment record (Data S5.5-5.9). Common gene sets can be deeply detected in two groups, Colorectal_Lung, HeadNeck_Lung and Melanoma_Lung as well as Breast_Lung, Colorectal_Lung and HeadNeck_Lung (Fig. 5D), with Breast_Lung and Melanoma_Lung excluded, respectively. Moreover, enrichment analysis reveals the four-subtype commonality since each group and the one not part of it share the same pathways in both cases (Fig. 5E; Data S5.5-5.8). For example, Breast Lung, Colorectal Lung and HeadNeck Lung seem to be affected by the FoxO signaling pathway, while lung tumors from melanoma (i.e., Melanoma Lung) are also predicted to suffer from it. Even though *NRAS* is the only identified gene they share, its great potential as a key clinical target should be emphasized. However, there is a close connection between Breast_Lung and Melanoma_Lung because we identified common gene set {*TP53, ASXL2, AURKB, CCND1, GATA3, NRAS, MAP2K4, GNA11, SMAD2*} that involve in dozens of signaling pathways (Data S2, Table S3.2.3 and Data S5.9).

Once again, shared pathways of the two groups of four subtypes are intersected to obtain convincing common signaling pathways of lung metastases. Actually, four significant signaling pathways (Fig. 5E) are credibly selected to drive distant migration to the lung, among which the PI3K-Akt signaling pathway and human papillomavirus (HPV) infection are frequently detected as common pathways of some lung metastases by enrichment analysis. Recent reports have shown that infection with HPV is associated with lung cancer, one of which provided evidence of O-linked-nacetylglucosamine transferase (OGT) acting in the spread of HPV-infected cervical cancer cells to the lung [55]. We also suggest the proliferation-associated genes cyclin D (*CCND1, CCND3*) and their associated cyclin-dependent kinases (*CDK4*) [56] as therapeutic targets for lung metastasis patients (Data S2, Table S3.2.1-3.2.3), especially those with HPV-positive.

Additional exploration of brain metastases also dawns on us what common factors make them bear the brunt of horrible invasion (see Supplementary Text 2.2 for more details). In general, cushing syndrome for liver metastases, microRNAs in cancer for lung metastases and MAPK signaling for brain metastases suggest adaptive changes of tumor cells in a new environment. Those common mutated genes in compared metastatic tumors need to be the top priority for treatments because they are assumed to accommodate tumor cells to new circumstances.

### Typical patterns with high metastatic tropisms

After the basic exploration of ‘fertile soils’ and ‘diligent sowers’ that urge the attraction and spread of cancer cells in the last two parts, we selected four well-known patterns [57] and showed them in greater details (Fig. 1C, Part 4). The four types of primary tumors and their most common metastatic tropisms are (1) colorectal liver metastasis, (2) NSCLC cancer with brain metastasis and (3) breast carcinoma and prostate cancer with bone metastases. We attempted to explain the causes of the typical patterns of cancer metastases in signaling pathways and recommend some crucial driver pathways and genes for therapeutic targets.

### Liver microenvironment facilitates colorectal cancer cells to further metastasize

The liver is the most common site of metastatic spread in colorectal cancer, and such metastasis constitutes a severe life-limiting factor [58, 59]. Here, we attempted to supplement some latent pathways and discuss the evolution of the metastatic cascade by comparing Colorectal_P (Fig. 6A, left) and Liver_P (Fig. 6A, right) to Colorectal_Liver.

**Figure 6.**
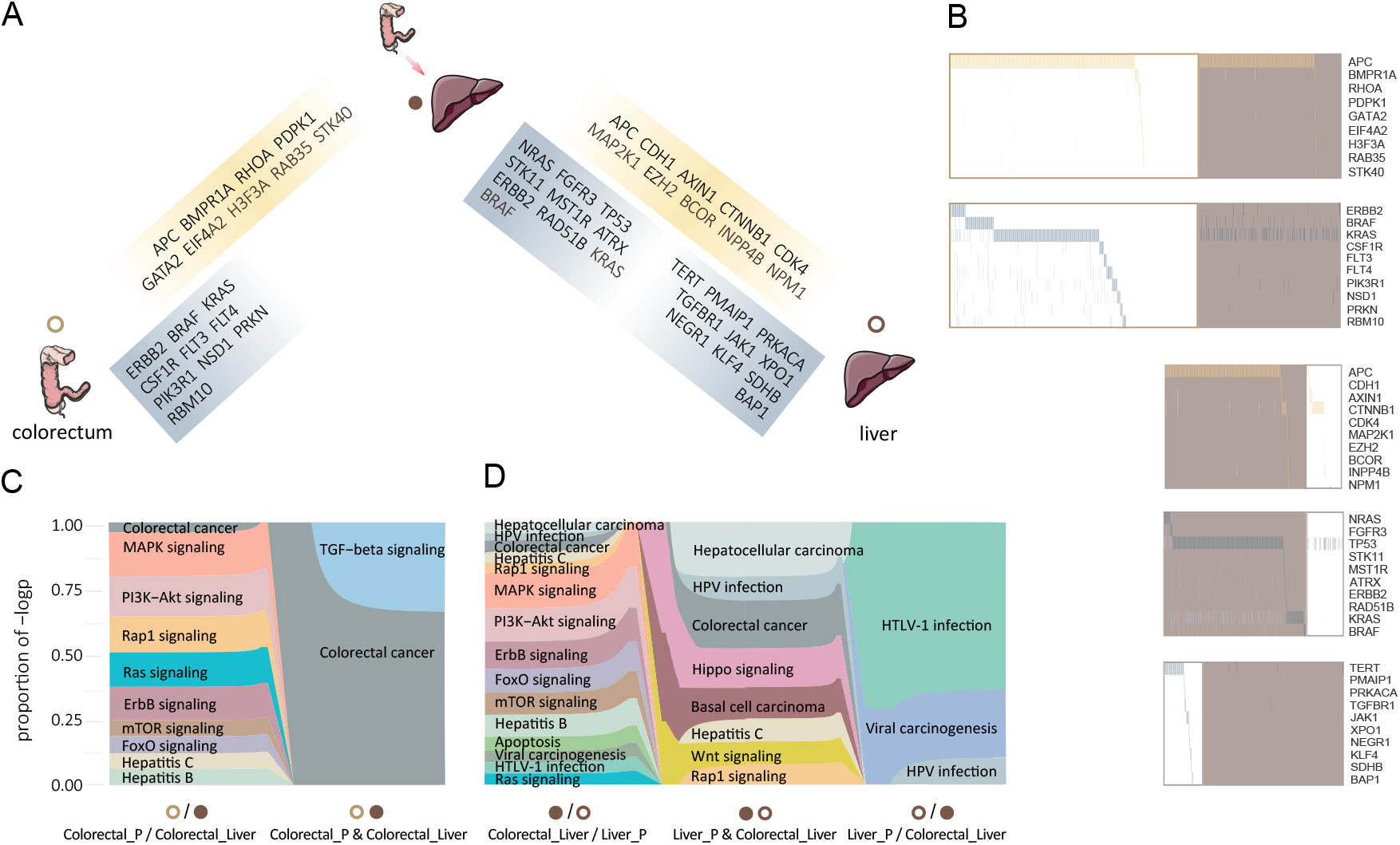
Signals in colorectal liver metastasis. (A) Significant common (yellow) or specific (blue) gene sets in the comparison between Colorectal Liver and Colorectal P (left) or Colorectal Liver and Liver P (right). Solid and hollow circles represent metastatic and primary tumors, respectively. (B) Heatmaps of the gene sets of two comparisons in (A). Solid and hollow squares represent metastatic and primary tumors, and yellow and blue represent common and specific results, respectively. (C, D) Significant signaling pathways obtained by enrichment analysis using gene sets in (A).

To detect the crumbling signal system, we started with the comparison between Colorectal P and Colorectal Liver. Approximately 70% of patients in both groups have the mutated suppressor gene *APC* (Fig. 6B), which was detected as one of the common driver factors for the two groups (Data S2, Table S4.1.1). As is graphically depicted in the Vogelstein model for colorectal cancer [60], mutated *APC* acts as the main initiator of the cancerous cascade and upregulates the Wnt signaling pathway, which could enhance colorectal cell proliferation [61]. The high mutation rate of *APC* in metastatic patients suggests that early tumor cells reserve the mutated *APC* and carry it to the liver. Additionally, the TGF-*β* signaling pathway was also identified as a driving force during the entire process by which a tumor cell performs its mission. However, this pathway only significantly affected a small number of patients with mutated *RHOA* or *BMPR1A* (Fig. 6B, C and Data S6.1). Once the tumor occurs, TGF-*β* changes its face from originally maintaining tissue homeostasis to promote development and invasion of tumors, such as inhibition of apoptosis and promotion of EMT and angiogenesis [62, 63].

As for the distinction between Colorectal_P and Colorectal Liver, the specificity of Colorectal_P is underlined by Ras, PI3K-Akt and MAPK signalings (Fig. 6C and Data S6.2). Four candidate genes, *ERBB2, KRAS, BRAF* and *PIK3R1*, are clearly exclusive (Fig. 6B), indicating a stronger signal response of the receptor tyrosine kinase (RTK)-related pathway in primary tumor cells. Dual pharmacologic inhibition of PI3K and MAPK has set a precedent to enhance the efficacy of anti-EGFR antibody therapy in patients with colorectal cancer [64].

The comparisons between Colorectal_Liver and Liver_P bears two remarkable resemblances to that of Colorectal P and Colorectal_Liver. What first attracts us is *APC* as the only common factor of the three groups (Fig. 6A and Data S2, Table S4.1.3). Low mutation rate as *APC* has in Liver_P, it is mutually exclusive and works with *AXIN1* and *CTNNB1* to regulate the Wnt signaling pathway (Fig. 6B, D and Data S6.3) [65]. This suggests that activation of Wnt will accompany the development of colorectal cancer and remains active even in the microenvironment of the liver. The second is the RTK-related pathway that appears when comparing Colorectal_Liver to Liver_P. It indicates that targeting mutated genes in this pathway is also applicable for Colorectal_Liver, but not for Liver_P patients.

More strikingly, we detected a series of signaling pathways associated with viral infection, such as HPV and human T-cell leukemia virus 1 (HTLV-1) infection, based on different specific genes of Colorectal_Liver or Liver_P (Data S6.4-6.5). Pathways related to hepatitis B and C are also noteworthy because they loom at the beginning of colorectal tumor growth (Fig. 6C). Currently, hepatitis B virus (HBV) and hepatitis C virus (HCV) remain the most important global risk factors for hepatocellular carcinoma [66]. Therfore, abnormal virus-related signaling pathways in colorectal cancer cells might become foretastes of the tendency to liver metastasis.

In general, increasing numbers of pathways specific to the liver are coming to the fore during the process of colorectal cancer cell spread. Metastatic patients suffer from a greater variety of disorders in the signaling system, as the specific signaling pathways disrupted in Colorectal_Liver largely overlap those in Liver P (Data S6.4-6.5). Therefore, we suggest a combination of different target agents necessary for Colorectal Liver patients and concern for liver-specific pathways in the early phase for Colorectal P patients.

### Bone microenvironment facilitates breast and prostate cancer cells to further metastasize

Bone is the most frequent destination for many cancer metastases, notably for tumors originating in the breast and prostate [67, 68]. What characteristics of bone make it an ideal environment for cell migration were investigated based on the comparison between the two types of primary tumors and their respective bone metastases, as well as between the two bone metastases.

It is evident from our results that mutation patterns in primary prostate (Prostate_P) tumors more closely resemble its respective bone metastases (Prostate_Bone) than those in breast since we only detected common gene sets between Prostate_P and Prostate_Bone (Supplementary Fig. S9A). Apart from the prostate cancer pathway represented by *TMPRSS2, HRAS* and *MTOR*, we also identified the endocrine resistance with *NOTCH4* taking its effect on the progression of prostate cancer [69] (Data S6.16). The existing mainstay of endocrine treatment for prostate cancer, especially sex hormone deprivation, contributes to fertile bone microenvironment that might promote bone metastases, which comes down to the evidence of endocrine-therapy-refractory prostate cancer [70, 71]. Despite unavailable results in the comparison between Breast_P and Breast_Bone, we noticed the pathway related to endometrial cancer, another sex hormone-driven cancer [72], but it was not significantly enriched by *CDH1* and *TP53* (Data S6.10). The outcome gives us an enlightenment to explore the connection between breast and endometrial cancer or other cancer groups with underlying ties, which will be discussed in the next part.

Moving on to the specificity of primary breast tumors and respective bone metastases (Breast Bone versus Breast_P), our attention is drawn to the calcium signaling pathway (Supplementary Fig. S9C, right). This pathway exerts a pivotal role in many cellular mechanisms related to breast cancer, such as cancer progression, metastasis, drug resistance and pain, especially after breast cancer cells migrate to bones [34]. This feature is also for prostate cancer because accumulation of intracellular calcium in prostate cancer cells can potentially favor bone metastasis development [73]. The awareness of calcium signaling as an early sign in patients with breast or prostate cancer might facilitate diagnosis and treatment before diseases become advanced.

Finally, we searched for signals displaying predilection for the bone microenvironment and analyzed the differences between breast and prostate tumor cells homing to bone. What we want to emphasize is the *PIK3CA* and *TMPRSS2* genes with a high frequency of mutations (Supplementary Fig. S9A, B), which are also respectively detected as specific genes of Breast_Bone and Prostate_Bone (Data S2, Table S4.3.3). While *PIK3CA* potentially serves as a future target for drug therapies of metastases in breast cancer [74], *TMPRSS2-ERG* increases bone tropism of prostate cancer cells and metastasis development [75]. Additionally, we notice *IKBKE* is positive in a very little percentage of Breast_Bone samples, but it exhibits mutual exclusivity with *PIK3CA* and *AKT1* (Supplementary Fig. S9B). The three genes control some special pathways, such as the Toll-like receptor and VEGF signaling (Supplementary Fig. S9D and Data S6.14).

In addition to the above two cases, common drivers detected for Lung_P and Lung_Brain also indicate organ tropism in metastasis because signaling pathways associated with brain diseases occur in primary NSCLC (see Supplementary Text 2.3). We can also observe an inclusion relation in the specific pathways of Lung Brain and Brain P. Therefore, it could add a further complication to antimetastasis since tumors reborn in distant regions are more heterogeneous than those born in the first place.

### Metastasis-promoting mechanisms for comparable cancers

In last section, we noticed that some genes specific in one type of cancer are prone to be enriched, in functionality or location, in the pathways related to the same smattering of other types of cancer. Therefore, we selected four groups of comparative objects: (1) breast carcinoma, ovarian and endometrial cancer; (2) bladder cancer, prostate cancer and renal cell carcinoma; (3) esophagogastric carcinoma and gastrointestinal stromal tumor and (4) melanoma and non-melanoma skin cancer to explore whether there is a certain parallelism within tumors before or after metastasis for each group (Fig. 1C, Part 5).

### Comparison of three gynecologic cancers

Starting with gynecologic neoplasms that have exhibited some interconnections above, we are mainly concerned about three prevalent female tumors: breast, endometrium and ovary. We carried out EntCDP within primary and metastatic groups to search for commonality among them and used ModSDP to perform comprehensive pairwise comparisons.

Not only are breast and endometrial cancer linked, but all three cancers, including ovarian cancer, have common pathways (Fig. 7A, D). Consistent with the above conclusion, we detected metastatic signals, cell adhesion molecules and their representative genes, *CDH1, CTLA4* and *CD276*, at the primary stage (Fig. 7A and Data S7.18). For patients with metastatic tumors, some signaling pathways involved in viral infections and endocrine function (Fig. 7D and Data S7.19) have been identified, giving an impetus to explore potential strategies for resistance mitigation to endocrine therapies and the carcinogenic mechanism of viral factors.

**Figure 7.**
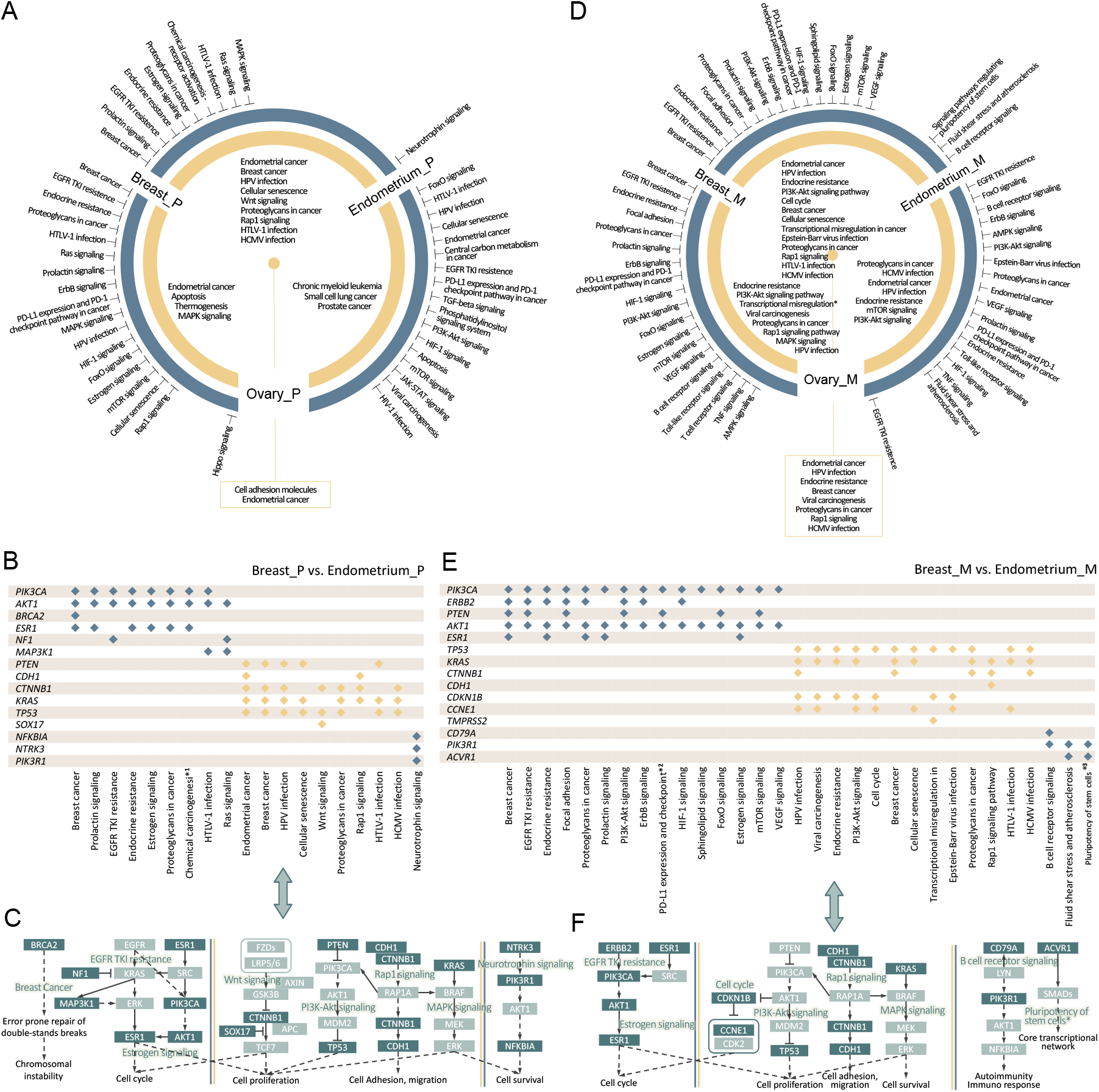
Pathway analysis for three gynecologic neoplasms in terms of primary and metastatic tumors. (A, D) Circos plot of significant common (yellow) or specific (blue) pathways in the three primary (A) or metastatic (D) cancers. The center of the circle points out signaling pathways the three cancers have in common. Both sides of each severed arc record the signaling pathways of the datasets at the end of the arc. (B, E) Subordinate relationships between identified genes and significant pathways in comparison between Breast_P and Endometrium_P (B) or Breast_M and Endometrium_M (E). (C, F) Identified genes and their functional relationships in several typical pathways in comparison between Breast P and Endometrium P (C) or Breast M and Endometrium M (F). The double arrow between and (C) / (E) and (F) points to the results of the same compared groups.

For breast carcinoma, endometrial cancer and ovarian cancer, there is a large degree of similarity between the outcomes of metastatic and primary groups, especially with respect to specificity (Fig. 7B, E). As we discussed in Part 1, all three gynecological cancers have common driver gene sets or pathways between primary and metastatic tumors (Fig. 3A). Actually, this similarity boils down to genetic similarity (Fig. 7B, E and Data S2, Table S5.1.1-5.1.3), except some metastases featured pathways (Fig. 7D), such as the focal adhesion, VEGF signaling, fluid shear stress and atherosclerosis.

For pairwise comparison, breast cancer patients have more specificity than the other two types, followed by endometrial cancer (Fig. 7A, D). The specificity of different comparative pairs overlaps in terms of different cancers in the same state or different states of the same cancer, which is represented by EGFR TKI resistance, estrogen signaling, PI3K-Akt signaling, etc. (Fig. 7C, F) This suggests that different combinations of genes can disrupt the same signaling pathway. We highlight the Hippo signaling pathway that plays an important role in governing ovarian physiology, fertility and pathology [76], which was identified as specificity of Ovarian_P and controlled by *LATS1, CTNNB1* and *NF2* (Data S2, Table S5.1.4-5.1.6 and Data S7.6). Recent studies have proven that loss of heterozygosity of the large tumor suppressor kinase 1 (LATS1) chromosomal region predisposes to ovarian cancer [77], and it regulates ovarian cancer cell invasion as a critical component of the Hippo signaling pathway [78].

### Comparison of three cancers related to males

We proceed to move on to three male cancers, bladder cancer, prostate cancer and renal cell carcinoma, and investigate whether they have similar findings to gynecologic tumors.

The results show that, their primary tumors do not have much in common, as contrasted to metastatic tumors (Supplementary Fig. S10A, D). Bladder_M, Prostate_M and Renal_M are manipulated by PI3K-Akt and MAPK signaling [79] of cellular proliferation, apoptosis and invasion regulation, with mutated *FGFR3, TP53* and *MET* (Supplementary Fig. S10B and Data S7.35). This indicates that the behavior of these tumor cells tends to be consistent after metastasis. Unlike gynecologic cancers, primary and metastatic tumors of the three male cancers are not apparently similar. Therefore, cancer treatment may be gender-specific and should be approached differently in targeted drug and analogy experiments.

Here, we focus on prostate cancer since it is the second leading cause of cancer death in men after lung cancer [79]. While *GSK3B* and *SPOP* were detected to regulate Hedgehog signaling for Prostate P relative to Bladder_P, driver factors for metastasis are mainly attributed to *CDKN1B, PTEN, BRAF* and *PIK3R1* as well as some important signaling pathways they disrupt (Data S2, Table S5.2.3 and Supplementary Fig. S10A, D, E). The study of Hedgehog signaling in prostate development has revealed its important roles in ductal morphogenesis and in epithelial growth regulation recapitulated in prostate cancer [80]. On the other hand, the comparison between prostate cancer and renal cell carcinoma brings some unexpected discoveries. Three EPH-related receptors, *EPHA3, EPHA5* and *EPHB1*, were recognized together as key members of the axon guidance pathway (Table 3 and Supplementary Fig. S10E, F) that leads to alterations in the proliferative, migratory and invasive potential of prostate cancer [81].

**Table 3.**
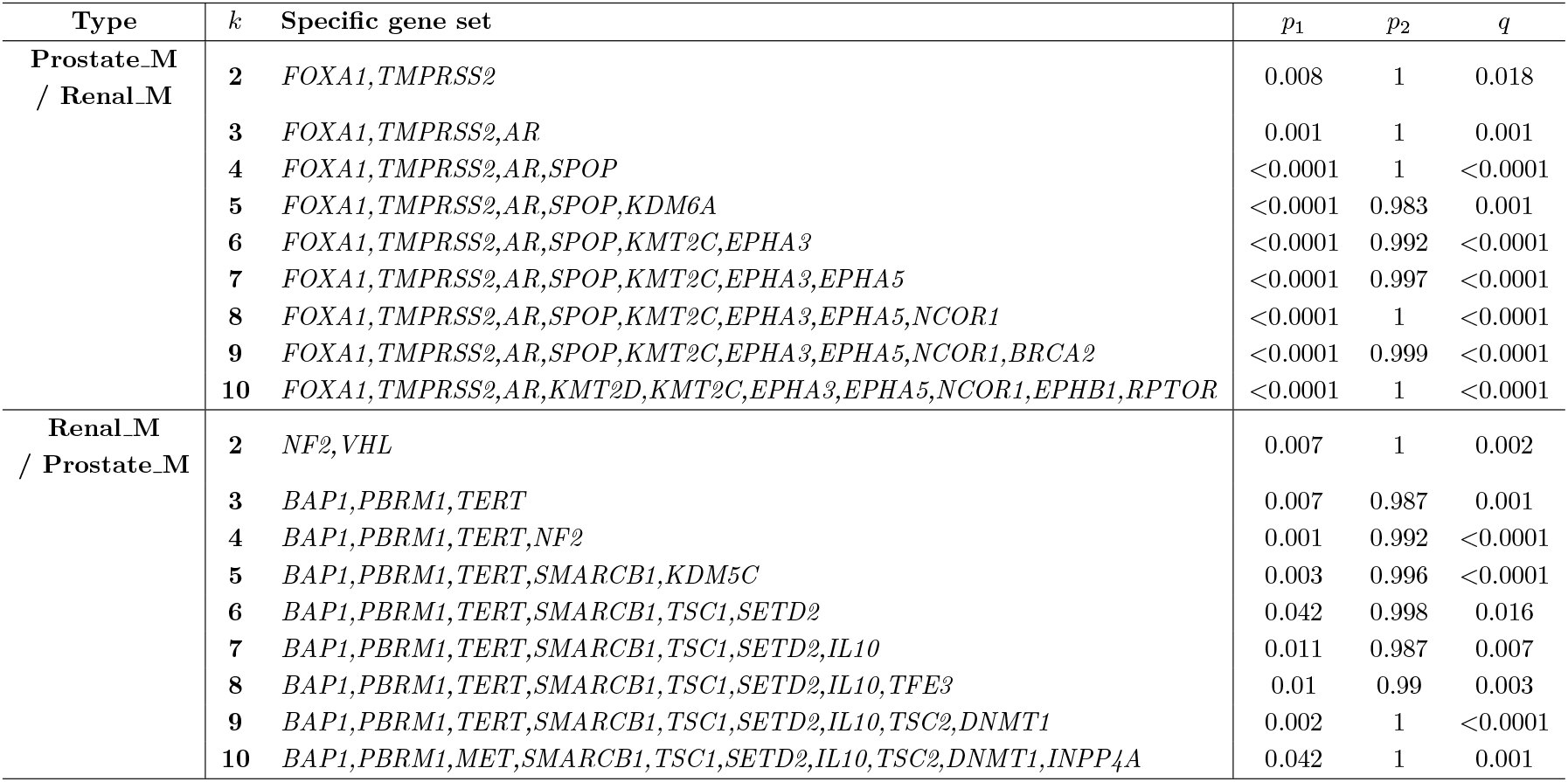
Prostate_M and Renal_M specific mutated driver gene sets relative to each other.

For the other two groups, we can only see similarities between esophagogastric carcinoma and gastrointestinal stromal tumor, and the specificity of melanoma and non-melanoma skin relative to each other. Their special performances are fully described in Supplementary Text 2.4. To sum up, the similarities shown by these groups of cancers suggest that we can draw reference from each other in the study of their pathogenesis and medications. In contrast, cancers in some groups that are very different from each other show their own characteristics under this comparison approach and require more nuanced treatment.

## Discussion and conclusion

Transition of the signaling system from primary to metastatic tumors prompts our thoughts on whether there is a clear purpose and destination for escaping tumor cells and what efforts tumor cells have made to colonize and adapt to the new environment. To settle some of the doubts, we gathered gene mutation data in both primary and metastatic tumors of 17 cancers from cBioPortal and discussed common or specific signaling pathways in categorized cancer types used two improved models, EntCDP and ModSDP. The two models improve the defects of the original model and enhance the accuracy of the search for signaling pathways. EntCDP solves the problem of inaccurate searches caused by the large variation in mutation rate across different datasets, while ModSDP adjusts the coverage and exclusivity of signaling pathways to a relatively equal status. The accuracy and efficiency of the models are guaranteed for both simulated and biological data. In addition, we retained the advantages of the original model, that is, we can discover driver genes with lower mutation rates [19]. At the same time, the signaling pathways we identified are constituted by more mutually exclusive alterations, which exert stronger selective pressure on tumors.

Our comprehensive signaling analysis that emphasizes metastases in combination with the form of contrasts and comparisons has rarely been considered in previous studies. EntCDP and ModSDP are appropriately used in the five perspectives according to the questions we want to investigate. For example, when comparing the signal changes of tumor cells from the same origin invading different regions, we only use ModSDP; in addition, to explain what common factors make some organs attract tumor cells for residence, we only employ EntCDP. On the other hand, the contents of the five parts are not totally independent. Three in four typical metastatic cases (Part 4) are chosen from the already analyzed primary tumors and their metastatic tropisms (Part 2 and 3), the results of which support previous conclusions such as the important role of calcium signaling in Breast Bone.

Many meaningful conclusions revealed by this study will be helpful for the understanding of the extent of signal changes during tumor metastasis. Targeted therapy regimens normally differ when a cancer patient is diagnosed at the primary or metastatic stage, because advanced patients generally have an accumulation of genetic mutations and greater tumor burden. Some cancers such as non-melanoma skin cancer and renal cell carcinoma are typical examples of signal changes during the process of metastasis, in which the Toll-like receptor signaling and the glucose/energy metabolism pathway have played an indelible role, respectively. However, our results show that breast carcinoma seems an exception since it is the only cancer for which both Breast_P and Breast_M have no specific signals relative to each other (Fig. 3A). Other two gynecologic tumors, endometrial and ovarian cancer, also retain more of the characteristics of primary tumors after becoming metastases. Particularly, endocrine hormone change and virus infection are suggested as common risk factor for the three female cancers. Cancers related to males, in contrast, seem inconsistent over time. For example, the main signaling pathway of prostate cancer is dominated by the Hedgehog pathway (disturbed by *GSK3B* and *SPOP*) in situ and the EPH-related pathway (represented by *EPHA3, EPHA5* and *EPHB1*) after distant metastasis.

Although the specificity of metastatic breast cancer as a whole relative to the primary one was not identified, some interesting results are obtained when we compared five different metastases from breast carcinoma regarding their special signal manifestations relative to the primary cancer as well as pairwise investigated these metastases. Breast_Liver is the most heterogeneous compared with other metastases from breast carcinoma since it has the most varied signals relative to Breast_P. We focus on lymph node because Breast Lymph Node shows no specificity to Breast P, which is of the same case with Lung Lymph Node. Combined with the results of the Melanoma Lymph Node, we suppose the FoxO signaling is a powerful driving force for tumor cells to settle in the lymph nodes. An opposing perspective turns on three organs attracting circulating tumor cells for residence. We identified cushing syndrome, microRNAs in cancer and the MAPK signaling as potential factors yielding three frequent sites for cancer metastasis, the liver, lung and brain, separately. More striking results are signaling pathways associated with glioma and brain disease and found in the comparison between Melanoma_Brain and Breast_Brain.

In general, smart cancer cells seem to anticipate the fate of migration due to the lack of space and food, so they attempt to switch on metastasis signals at the stage of primary tumors. Once tumor cells settle down, they live by disrupting or activating pathways related to their new environment. Such conclusion can be verified by the three well known and frequently reported metastatic patterns, because we detect hepatic signals in Colorectal P and Colorectal Liver, as well as central nervous system disease pathways in Lung_P and Lung_Brain. Additionally, we highlight abnormal virus-related signaling pathways as an early sign of hepatic metastasis from colorectal cancer, FoxO signaling for brain metastasis from NSCLC and calcium signaling for bone metastasis from breast or prostate cancers.

In summary, comprehensive investigations on transition of pan-cancer signaling systems from primary to metastatic tumors are conducted here, and many interesting results are obtained, some of which may deserve further investigations. We expect that this study will provide a valuable resource for cancer researches, and transform our knowledge about signals in cancer metastasis into alternative clinical practice for advanced patients.

## Materials and Methods

### EntCDP: entropy-based common driver pathway identification model

Considering the impact of different cancer mutation rates, we introduce entropy to the objective function of EntCDP for identification of common mutated driver gene set *M* containing *k* gene across multiple cancer types or subtypes. The form of EntCDP is shown as follow:

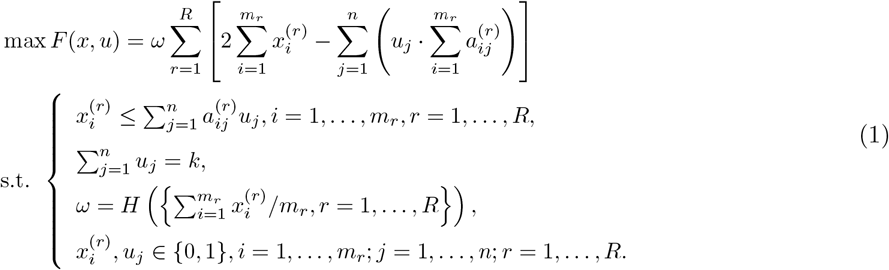

EntCDP takes *R* gene mutation matrices 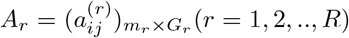 of *m*_*r*_ samples with *G*_*r*_ genes as input (Fig. 2A). 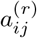 indicates the mutation status of the *j*^*th*^ gene of the *i*^*th*^ sample, with a value of 1 (mutation) or 0 (no mutation). First, the number of columns is expanded to |*G*| =| ∪ *G*_*r*_|, so that the same column of each matrix corresponds to the same gene. If the corresponding gene for the column is not previously available, the column is set to all zeros. To realize the search of gene set by binary programming, *u*_*j*_ is used as an indicator that can judge whether column *j* of the mutation matrix *A*_*r*_(*r* = 1, …, *R*) belongs to the required submatrix *S*_*r*_, whose columns represent gene in *M*, and 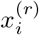 is another indicator that answers whether the entries in the *i*^*th*^ row in *S*_*r*_ are all zero or not.

The objective function of EntCDP is the product of two parts: *ω* · *C*_*m*_(*M*). While *ω* is an entropy-based weight, *C*_*m*_(*M*) describes the total coverage and exclusivity of *M* among *R* cancers, and is actually the objective function of the ComMDP. The initial idea is to assign as much coverage and exclusivity as possible to each cancer type while ensuring that the differences between them were not too large. Specifically, 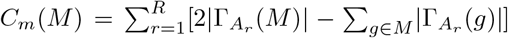, where 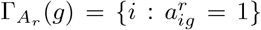 denotes the set of samples of the *r*-th cancer type with gene *g* mutated, and 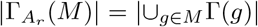 represents the coverage of *M* in *A*_*r*_. Importantly, the weight *ω* is related to coverage rates across different cancers. *H*(*) is the information entropy of coverage rates and defined below:

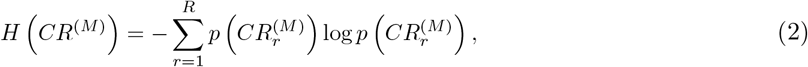

where 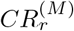 is the coverage rate of the gene set *M* in cancer type *r*, namely 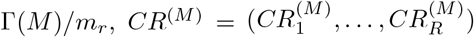, and 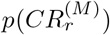 is normalized *CR*^(*M*)^ with form 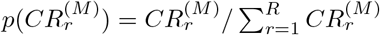. In this case, as *ω* increases, the variation in the coverage of *M* across different cancers decreases. Consequently, the ultimately identified gene set strive to achieve high coverage across all cancers (Fig. 2A, right), more closely aligning with the biological driver pathways than ComMDP (Fig. 2A, left).

In addition, we find that the following relationship holds:

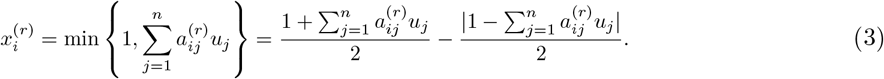

Then, the model can be further reduced to an optimization problem that contains only *n* variables:

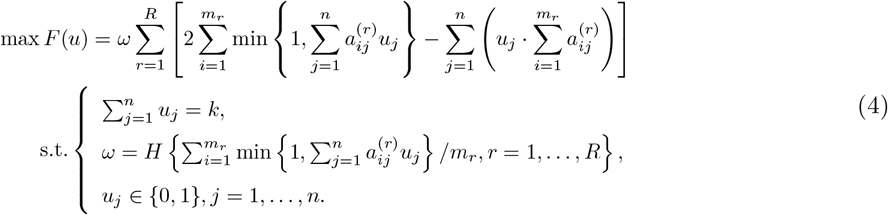

We solve the EntCDP (eq. 4) using genetic algorithms.

### ModSDP: modified specific driver pathway identification model

To find a specific mutated driver gene set *M* for the *R* cancer types relative to the *T* types, we proposed a novel specific pathway identification model called ModSDP that partly compensates for the sacrifice of exclusivity in SpeMDP. The binary linear form of ModSDP is shown in eq. 5.

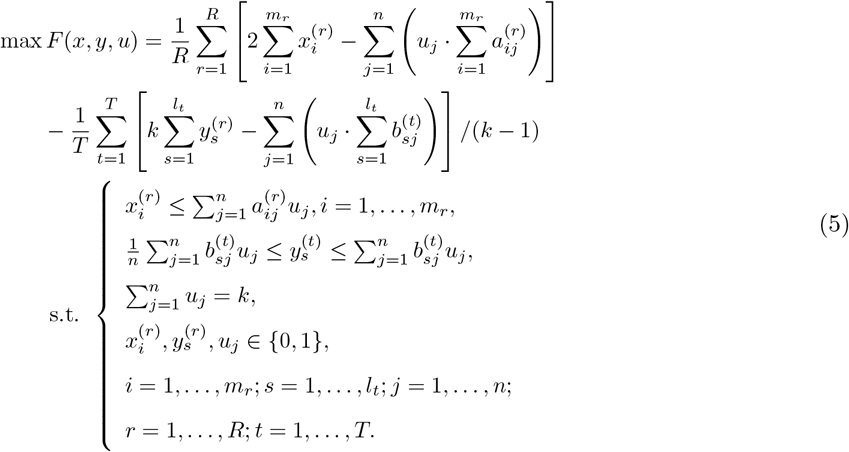

ModSDP aims to identify the gene set *M* that exhibits large coverage and high exclusivity in each cancer of group *A* but lacked these properties in the group *B*. It takes two sets of gene mutation matrices 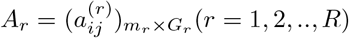 samples with *G*_*r*_ genes and 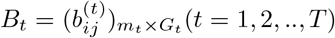 samples with *G*_*t*_ as input (Fig. 2D). Similarly, the number of columns of all matrices is expanded to the same. 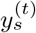 indicates whether the entries in row *s* in the required submatrix of *M* are all zero. Other variables are defined in the same way as EntCDP.

The modified objective function of ModSDP is roughly the difference between the coverage and exclusivity weights of group *A* and group *B*, which is equal to 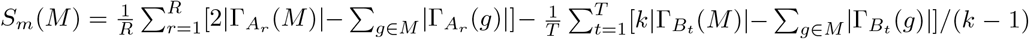. Compared with SpeMDP, the coefficient before coverage (i.e., 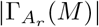 and 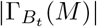) of group *A* in ModSDP was adjusted from *k* to 2, while the coefficient of group *B* is globally divided by *k* − 1. We aim to balance the coverage and exclusivity of the identified gene set for group *A*, while also aligning the total weight scale between group *A* and *B*. The detailed principle of this improvement is analyzed in Supplementary Text 1.1.

We maximize its binary form (eq. 5) by IBM ILOG CPLEX optimizer, a high-performance mathematical programming tool that can be used in combination with multiple programming software to solve linear programming and binary integer programming problem.

### Permutation test

A gene set identified by models should undergo a permutation test to check whether it truly fits the high coverage and high exclusivity of a signaling pathway. For EntCDP, we want to detect a set of genes which had significantly high mutual exclusivity and large coverage in two or more cancer types simultaneously; but for ModSDP, we wanted to identify a driver gene set specific to one or a group of cancer types (say, *A*) versus another group of cancer types (say, *B*). In other words, we required the detected genes to have significantly high mutual exclusivity and large coverage in the group *A* but not in *B*.

The idea of the permutation test related to pathway identification was mentioned in [82], which defined the *p*-value of the test as the number of times that the objective function value of *M* ^′^ obtained by randomly shuffling optimal *M* is more than that of *M* among 1000 successive iterations. Random shuffling here refers to the random transposition order of all samples for each gene in *M* but does not change the number of samples who have that gene mutated, so that all genes in each sample no longer exhibit original mutation patterns.

We performed a permutation test to assess the significance of any result obtained by EntCDP or ModSDP. We permutated the mutations independently among samples to preserve the mutation frequency of each gene. Two kinds of significance were calculated: (i) individual one (such as the p-values *p*_1_, *p*_2_ in Fig 3B, D, F) measuring the significance of a gene set in a certain mutation matrix, where 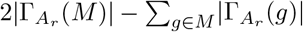 or 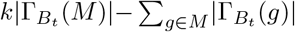 was used as the statistic; (ii) overall one (such as the p-value *p* in Fig 3B, D, F) measuring the significance of a gene set by viewing all the mutation matrices as a whole, where the weight *ω* · *C*_*m*_(*M*) and the weight *S*_*m*_(*M*) were used as the statistics for EntCDP and ModSDP, respectively.

We sorted results of each comparison, that is, significant driver gene sets with the number of genes, if any, from 2 to 10 into a individual table in Data S2.

### Genetic algorithm (GA)

Due to the nonlinearity of EntCDP, it is not suitable for above optimizer, but can be solved with the help of genetic algorithms.

GA, a heuristic algorithm, simulates biological evolution and searches for the best individuals by constantly retaining and updating the fittest individuals in a population. When applying GA to EntCDP, each individual essentially represents a certain gene set, and a bit vector of an individual indicates whether some genes are selected into the submatrix, so the hypothesis space of individuals is 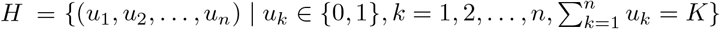. The fitness function is defined as the objective function of EntCDP, and the fitness value of the *i*^*th*^ individual, *r*_*i*_, is the ascending rank of its corresponding objective value. Further, the probability of the individual selected as the one of the parents is defined as 2*r*_*i*_*/P* (*P* + 1), where *P* is the population size. Next-generation populations are generated from *P* pairs of parents that are randomly generated by individuals selected according to the above probability. Each individual inherits the overlapping binary coding site from both parents, and a few of the remaining sites are randomly selected and encoded to 1 to guarantee that this individual satisfies the definition of the hypothesis space. Subsequently, the individual develops a mutation from 1 to 0 at one site and simultaneously from 0 to 1 an another site with a certain probability.

The above procedure is considered one iteration, and we use 1000 iterations. ‘Local search’ is performed to avoid local optimal solutions when the current solution after five generations does not improve. In this case, the optimal solution is subjected to multiple rounds of mutations at two sites as described above. If such a mutation improves the current solution, a new individual after the mutation is retained, that is, the iteration jumps from the local optimal solution. Otherwise, the iteration continues until the solution remains unchanged after 10 iterations, and the optimal individual and the optimal value are found in the final population.

### Pathway Enrichment Analysis

For the results from EntCDP or ModSDP, we combined all the genes in the corresponding table regardless of *k* for more information, and assigned them to pathways (KEGG PATHWAY) on the website of David Bioinformatics Resources (https://david.ncifcrf.gov/home.jsp). Outcomes of pathway enrichment analysis for each gene set are available in Data S4-8 for five parts respectively.

Therefore, an identified signaling pathway was forged by doubling statistical tests in which a gene set is put through the permutation test and a significantly enriched pathway was screened out after testing the significant gene set in enrichment analysis.

### Simulation data

To systematically evaluate the performance of EntCDP and ModSDP, we first construct a series of simulated gene mutation matrices from two perspectives (Supplementary Text 1.2 ∼ 1.4).

On the one hand, we compare EntCDP with ComMDP to test whether EntCDP can identify the gene set with more uniform coverage across multiple simulated gene mutation dataset than ComMDP. Additionally, we compare ModSDP with SpeMDP to test whether ModSDP can identify the gene set that is more mutually exclusive than SpeMDP, though at the expense of very little coverage. On the other hand, we also construct the simulation data to evaluate the accuracy and efficiency of both models.

The simulated gene mutation matrix is created by embedding mutation modules (i.e., pseudo gene sets) to an all-zero matrix, implementing ‘single mutation’ or ‘double mutation’ within the mutation modules, and introducing ‘background mutations’ outside the modules. Basically, ‘single mutation’ means that only one gene in each selected sample is randomly chosen to be mutated at a higher mutation rate, while ‘double mutation’ means that mutations occur with a certain probability in two randomly selected genes simultaneously. Additionally, ‘background mutation’ indicates that genes not in the module are mutated in at most three samples. We designed 8 sets of simulation data (Simulation 1 ∼ 8, abbreviated to Sim1 ∼ 8) for EntCDP and 5 sets of simulation data for ModSDP (Sim9 ∼ 13).

### Cohort overview

All biological data used in this study are available on cBioPortal for Cancer Genomics (http://cbioportal.org/msk-impact). We select gene mutation profiling of 17 cancers from 7734 samples and divide each cancer dataset into two groups based on primary and metastatic tumors except for hepatocellular carcinoma (HCC) and glioma whose metastatic tumors are rarely present (Fig 1A).

To investigate the differences in tumor metastasis in different sites, we further divide some metastatic cancer data into multiple subtypes (Fig 1B). The name and size of each dataset are detailed in Data S1. To be concise, we use the suffixes ‘ P’ and ‘ M’ to denote the primary and metastatic cancers, respectively (for example, Colorectal M represents metastases originating from colorectal cancer, and correspondingly we use Colorectal P to represent primary colorectal cancer, etc.). Furthermore, for one tumor metastasizing to others, such as breast carcinoma metastasizing to lung, liver, bone, lymph node and chest wall, we denote them as Breast Lung, Breast Liver, Breast Bone, Breast Lymph Node and Breast Chest Wall, and so on.

## Supporting information

Supplementary File - Supplementary text, figures, tables, etc.

Data S1: biological data discription

Data S2: tables of results

Data S3: analysis results for part1

Data S4: analysis results for part2

Data S5: analysis results for part3

Data S6: analysis results for part4

Data S7: analysis results for part5

## Acknowledgments

This work was supported by the funding from the National Key Research and Development Program of China (2022YFA1004800).

## Author contributions statement

Author contributions: J.Z. conceived and supervised the study. W.Z. gathered the data and performed the analyses. J.Z. and W.Z. improved the methods and interpreted the results. All authors wrote, revised and approved the manuscript.

## Additional information

Competing financial interests: The authors declare no competing financial interests.

