## Supplementary File - Supplementary text, figures, tables, etc. for "Pan-cancer study on transition of signaling systems from primary to metastatic tumors"

##### This PDF file includes:

Supplementary Text
Figs. S1 to S11
Table S1
Legends for Data S1 to S7
SI References

##### Other supporting materials for this manuscript include the following:

Data S1 to S7

#### Supplementary Text

#### 1. Methods

**1.1 Principle for ModSDP.** Here, we attempt to explain the principle or our notion of the structure of ModSDP.

First, it is reasonable to limit the weight of each cancer of group $B$ in ModSDP to no less than zero, which is to avoid the case where group $A$ takes quite small positive weight and group $B$ occupies quite large negative weight so that the identified gene set belong to neither group $A$ nor $B$. However, this notion is not achieved by the constraint of SpeMDP because additional restrictions would reduce the search scope of possible gene sets and the small negative weight of group $B$ is acceptable. So attention is finally paid to objective function, and we expand the coverage of group $B$ by $k$ times. As a result, newly defined weight of group $B$ is always positive.

To unify the order of magnitude, the coefficients of group $A$ should be changed simultaneously. Upon further analysis, we argue that the adjustment of coefficients of group $A$ is not essential, so the imbalance of coverage and exclusive of group $A$ are easy to overcome by restoring $k$ to the initial 2. On the other hand, the exclusivity of group $B$ must sacrifice at the cost of such imbalance as our analysis above, so it cannot be restored. However, once adjust for group $A$, the coefficient of group $B$ naturally need to adjust to reach a balance with that of group $A$. To ensure that the weight of group $B$ remain non-negative, we considered reducing the combinations of coverage and exclusive at the same time. The rationality of dividing by $k-1$ lies in that the coverage of group $B$ expand $k$ times while the exclusivity remain unchanged, and the original model SpeMDP is also a special case when $k=2$.

**1.2 How to design the simulation data to test the identification ability of the models.** Before applying our models to biological datasets, we had tested the effectiveness of the two models on simulation data beforehand. The form of the simulation data should resemble real data in terms of sparsity and sample size in order to convince our results on biological datasets.

In general, the simulation data takes the method of embedding the 'mutant module' in the zero matrix. These 'mutant module' is usually expected to be identified by models. Each module is a binary 0-1 matrix, where every entry are placed to 1 in a certain mutational mode, which means there might be a genetic mutation in one sample. There are three types of pattern: single mutation, double mutation and background mutation. Specifically, 'single mutation' means that only one gene in each selected sample is randomly chosen to be mutated with a higher mutation rate $p$, and if the event occurs, the other genes in this sample are mutated with a small mutation rate $p_0$. $p$ and $p_0$ represent coverage and mutual exclusivity respectively, and by default $p=0.85$ and $p_0=0.04$. Similarly, 'double mutation' means mutation occurs with certain probability in two randomly selected genes simultaneously. Moreover, background mutation implies that genes not in the module are mutated in at most three samples. In general, genes outside those modules are mutated by 'background mutation'.

When measuring the rationality and validity of the model, we usually use simulation matrices varying in sample size, mutation rate and mutant pattern. When making comparative analysis with ComMDP and SpeMDP, problem-driven simulation datasets are constructed to compare the order and efficiency of identification.

**1.3 Simulation data for EntCDP.** In addition to the Sim1 in text, Sim2 and Sim3 are constructed to illustrate the advantages of EntCDP over ComMDP. In both cases, module 2 has more uniform coverage than module 1 in different cancers, and therefore has greater biological significance. The calculation results are shown in Table S1.

Sim2: A large gap between the sample size of two cancers based on Sim1 (Fig. S1A). On the left side of Fig. S1A, module 1 covers 85% and 40% of samples in both cancers, and the proportions for module 2 are 65% and 70% on the right. Using ComMDP, the objective function values of module 1 and module 2 are calculated as $(2\times17-20)+(2\times4-5)=17$ and $(2\times13-16)+(2\times7-8)=16$, thus ComMDP identified module 1. Using EntCDP (Fig. S1A, right), since $H\left(CR_1^{(module\,1)}, CR_2^{(module\,1)}\right)=0.9044$, $H\left(CR_1^{(module\,2)}, CR_2^{(module\,2)}\right)=0.9994$, the objective function values of module 1 and 2 are therefore calculated as $17\times0.9044=15.3745$ and $16\times0.9990=15.9842$, respectively. Therefore, EntCDP chooses to identify the module 2.

Sim3: Identification of gene sets with four genes in three cancers. On the right side of Fig. S1B, module 1 covers 85%, 80%, and 33% of samples in the three cancers, while the proportion for module 2 is 65%, 60% and 60% on the right. Using ComMDP (Fig. S1B, left), the objective function values of module 1 and module 2 are calculated as $(2\times17-20)+(2\times8-10)+(2\times5-7)=23$ and $(2\times13-16)+(2\times7-9)+(2\times9-11)=22$, so ComMDP identifies module 1. Using EntCDP (Fig. S1B, right), since $H\left(CR_1^{(1)}, CR_1^{(2)}, CR_1^{(3)}\right)=1.4846$, $H\left(CR_2^{(1)}, CR_2^{(2)}, CR_2^{(3)}\right)=1.5821$, the objective function values of module 1 and module 2 are calculated as $23\times1.4846=34.1458$ and $22\times1.5821=34.8062$ respectively. Therefore, EntCDP chooses to identify the module 2.

Sim4 $\sim$ 8 are set below to evaluate the identification ability of EntCDP.

Sim4: Zero matrices $A_1\sim A_3$ represent three types of cancers, with column number of 900 and row number of 500, 550 and 600 respectively, indicating that the number of genes is 900 and the number of samples of each cancer is 500, 550 and 600 respectively. For each cancer $(r=1,2,3)$, nine groups of mutant modules are embedded in $A_r, r=1,2,3$, and the number of columns (i. e., $k$) was $2,3,...,10$ (i. e., $I$), respectively. For each $I$, $M_r^{(I)}$ occupy the same columns in the corresponding $A_r$. Each module is implemented 'single mutation' as described in Appendix 2.2 with the mutation rate fixed at 0.85 and the proportion of samples who are selected to have such mutation pattern is 0.8, 0.7 and 0.75 for each cancer. Genes outside those modules are mutated by 'background mutation' pattern. The simulation is repeated five times in succession. In Sim4, all genes corresponding to the nine groups of modules can serve as common signaling pathways for the three cancers, and we expect EntCDP would identify 9 gene sets in turn by sequential deletion of the identified modules from 2 to 9.

The identification results are shown in text Fig. 2B. EntCDP can stably identify signaling pathways with the number of genes within 7. When $k = 8$, the identified gene set sometimes comprises seven genes in the embedded module plus one gene outside the module, and the outside gene is found to be background gene. Inaccuracy also exists when $k = 9$ and 10, in which the maximum number of errors is 2. The explanation for the above faults is: the coverage and exclusivity are reflected by gene mutation arrangement; larger number of genes in module eliminates the contribution of each gene to the two characteristics, which may result in the use of a single mutant gene in the module being less effective than the background mutant gene. This is also virtually because of a few non-exclusivity of the simulation itself adding to the 'single mutation' pattern.

Sim5: Unlike Sim4, larger range of sample size is settled for the three cancers with 200, 500, 800, respectively. Under this construction, gene sets corresponding to the 9 groups of modules can also serve as common signaling pathways of 3 cancers.

As shown in Fig. S2B, the accuracy of Sim5 is similar to Sim4. EntCDP can also identify pathways with small number of genes (2∼7), but failed in the large number of genes (8∼10) with non-significant results at $k = 9$ and 10. The reason for this phenomenon is that when the sample size is large, the sample size of each gene mutated in modules increases accordingly, thus enhancing the role of each gene on the coverage; once a gene in the module is replaced by a background mutation gene, the overall coverage weight significantly reduces, so it is hard to pass the permutation test.

Sim6: The number and size of cancers are same as that in Sim4, but we add a small number of 'double mutation' genes in embedded modules with the proportion of 2% and 4%. Under this construction, gene sets corresponding to 9 modules can also be viewed as common pathways for 3 cancers. When the proportion of 'double mutation' is 2%, EntCDP can stably recognize the signaling pathway with less than or equal to 7 genes, but when the proportion is 4%, EntCDP fails to recognize modules containing less than 7 genes (Fig. S2B). After the addition of 'double mutation', there are 3 wrong genes outside the embedded modules and non-significant cases when $k = 10$, indicating that 'double mutation' will increase the instability of identification.

Sim7: Zero matrices $A_1 \sim A_5$ represent five types of cancer. The number of genes for each cancer is 900 and the number of samples are 300, 350, 400, 450 and 500, respectively. We embed five groups of mutant modules $M_r^{(I)}(r, I = 1, ..., 5)$ containing 6 genes in $A_1 \sim A_5$. Each module is subjected to 'single mutation' with the mutation rate fixed at 0.85, and the proportion of the sample selected to mutated is $0.75 - \phi(r) * I, \phi(r) = 0.01r$. The simulation was repeated five times in succession.

Under this construction, the corresponding gene sets of five groups can serve as common signaling pathways for five cancers, but the average coverage of pathways decreases and the coverage difference expands in sequence, weakening the two characteristics of pathways. It is expected that the EntCDP can identify the five gene sets sequentially. The identification results are shown in text Fig. 2C, in which the bias in the order of identification suggests that GA algorithms are occasionally trapped in the local optimum.

Sim8: Unlike Sim7, larger range of sample size are settled for the five cancers with 200, 400, 600, 800 and 1000, respectively. It is also expected that the EntCDP can identify five gene sets sequentially. The result illustrated in Fig. S2C is similar to Sim7, which indicates that the accuracy of the identification is not affected by the sample size.

**1.4 Simulation data for ModSDP.** Sim11 $\sim$ 13 are set below to evaluate the identification ability of ModSDP.

Sim11: Zero matrices $A$ and $B_1 \sim B_3$ represent four types of cancer with column number of 900 and row number of 500, 400 and 600 respectively. We embed $M^{(I)}$ and $M_r^{(I)}(t = 1, 2, 3, I = 1, 2, ..., 9)$ in group $A$ and $B$ respectively, whose number of column is $I + 1$ and mutation rate is 0.85. For each $M^{(I)}$, 85% of samples are chosen to be conduct 'single mutation'. For $M_r^{(I)}$, we perform 'double mutation' on 50%, 55% and 60% of samples with cancers $B_t, t = 1, 2, 3$, respectively. The simulation is repeated five times in succession. As can be seen in Fig. S3, ModSDP can completely identify the first 8 modules, but occasionally has wrong identification when $k = 9$ and a maximum of 3 false genes for 10. The reason behind it is the same as Sim4, and it is also related to the stronger coverage of group $B$ than exclusivity in ModSDP.

Sim12: Matrix size is same as that in Sim10 described in text. We embed $M^{(I)}$ and $M_t^{(I)}(t = 1, 2, 3, I = 1, 2, 3, 4)$ in $A$ and $B_t$ respectively, whose number of column are six. For each $M^{(I)}$, 85% of samples are chosen to be conduct 'single mutation', and its mutation rate is $0.95 - 0.05 * I$. Whereas for $M_r^{(I)}$, 60% samples are uniformly selected to conduct 'double mutation' with mutation rate of $0.9 - \lambda(t) * I, \lambda(t) = 0.01 * t$. Under this construct, all four gene sets can serve as specific pathways of $A$. Sequentially deleting of the identified modules, the ModSDP can identify the gene sets corresponding to 4 modules as expected (Fig. S4A), indicating its excellent identification ability in terms of different combinations of mutation rates.

Sim13: Based on Sim12, one and two types of cancer are added for group $A$ and $B$ respectively, namely $A_1, A_2$ and $B_1 \sim B_5$, with column number of 400 and 500, as well as 400, 400, 500, 500 and 600. For $M_r^{(I)}$ embedded in $A_r$, the mutation rate is $0.95 - \phi(r) * I, \phi(1) = 0.04, \phi(2) = 0.05(r = 1, 2, I = 1, 2, 3, 4)$ and for $M_t^{(I)}$ embedded in $B_t$, the mutation rate is $0.9 - \lambda(t) * I, \lambda(t) = 0.01 * t(t = 1, ..., 5, I = 1, 2, 3, 4)$. Under this construction, all gene sets corresponding to the four groups can also serve as specific signaling pathways for group $A$. As well as Sim12, the results (Fig. S4B) indicated that ModSDP has great identification accuracy and efficiency facing more cancer types.

#### 2. Extended Description of Results

##### 2.1 Supplement to Part 2.

***Different signals in tumor cells originating from NSCLC.*** As one of the subtypes of lung cancer, NSCLC is known as malignant tumors that seriously threaten human health, and it is the metastatic phase that contributes to most of the mortalities (1). We considered five migratory tendency destinations including bone, brain, liver, fluid pleura and lymph node to figure out the role of NSCLC-derived signaling pathways in invasion and metastasis.

Similar to analysis on breast cancer, Lung_P dataset is taken to explore the signal change pre and post metastasis. However, we only identify specific genes for primary tumors which correspond to the results discussed in text Part 1. Nevertheless, the specificity of metastatic relative to the primary cancer can also be seen in comparison of the five types of metastases (Fig. S5B and Fig. S6A).

Metastatic tumors growing in lymph node or pleura seem to preserve more characteristics of primary NSCLC since we detect the non-small cell lung cancer pathway when respectively comparing the two metastases to other subtypes (Fig. S6B, D and Data S4.24-4.30), while the other three metastases are not the case. Furthermore, significant results can be observed in case of Lung_Lymph Node relative to Lung_Pleura (Fluid), but not vice versa, indicating extraordinary heterogeneity in lymph nodes. In this case, the chemokine signaling pathway (2) might promote the formation of metastases in the regional lymph nodes from lung via activation of downstream genes on PI3K-Akt and RAF-MEK-ERK (Data S4.27). Some other enriched pathways also show up in the comparison of Lung_Lymph Node relative to Lung_Bone, Lung_Brain and Lung_Liver, with 9, 25 and 14 overlaps between them (Fig. S6C). There are 7 overlapping pathways belonging to all the four cases, among which the FoxO signaling is notable. Mutations of *EGFR* and *BRAF* on it (Fig. S6C) may increase the metastatic potential of NSCLC cell towards to lymph nodes (3).

Pleura has a unique geographical advantage for the invasion of NSCLC cells, so we continue to investigate the relationship between driver oncogene alterations and metastatic pattern of lung based on the results of pleura. *EGFR* on the proteoglycans in cancer pathway (Fig. S6D, E) stands out for its high mutation rates on patients who have NSCLC and secondary tumor lesions in the pleura. This coincides with the recent result that *EGFR*-mutated tumors preferentially spread to the pleura (4).

In addition, a recent discovery looked into the mechanisms behind how IL6 promotes metastatic cells colonization from NSCLC to brain via the JAK2/STAT3 signaling pathway (5). In our study, we also identify the JAK-STAT signaling pathway for Lung_Brain relative to Lung_Bone and Lung_Pleura (Fluid) (Data S4.18, 4.20).

##### 2.2 Supplement to Part 3.

***Brain as the preferred site for cancer metastases.*** Most brain metastases is credited to lung, breast and melanoma (6), so the mutation data collected from the three origins (text Fig. 1B and Fig. S7A) may help to understand the genome of brain metastases.

Similar to the analysis for lung cancer, we combine enrichment results of two cases and identified five latent common signaling pathways for the three subtypes (Fig. S7B, C). In addition to pathways related to particular cancers, MAPK signaling pathway is highlighted (Fig. S7C). Targeted inhibitors of its pathway components, such as RAF, MEK or ERK have been employed as a strategy to block aberrant MAPK signaling in lung (7), melanoma (8) and breast (9) cancers with brain metastases.

Furthermore, we are also curious about breast and melanoma-derived metastases because of their abundant significant pathways in common, which indicates an underlying connection, for instance, metastatic patterns or driver factors, between the two organs. Besides the direct regulation of glioma-centred pathway network such as Alzheimer disease (including *HRAS*, *NRAS*, *CDK4*, *AKT3*, $k = 8$), mTOR signaling (including *TP53*, *BRAF* and *FLCN*, $k = 6$) that has been reported to predict brain metastasis (10) can also be detected for Breast_Brain and Melanoma_Brain (Data S2: Table S3.3.2 and Data S5.11).

##### 2.3 Supplement to Part 4.

***Brain microenvironment facilitates NSCLC cancer cells to further metastasize.*** Brain metastasis (BM) is associated with poor survival outcomes and poses distinct clinical challenges in patients with NSCLC (7, 11). This pattern in Appendix 2.2 is not the clearest picture, so we paint it in details and construct the pathway map in comparison with Lung_P and Brain_P.

With the evolution of NSCLC, mutated *ERBB2*, *EGFR*, *BRAF*, *KRAS*, *MET*, *EML4* and *KIF5B*, common genes of Lung_P and Lung_Brain (Data S2: Table S4.2.1), overactivate the MAPK, Ras and ErbB signaling pathways that are related to NSCLC (7) (Fig. S8A, C and Data S6.6). In addition to these seven driver factors, oncogene driver *ROS1*, the eighth genes detected in gene set when $k = 8$, has proven to be a drug target for treatment of brain metastases in *ROS1*-positive NSCLC (11). Additionally, commonality can also be observed in the detection of central nervous system disease pathways, such as Alzheimer disease (12), and metastasis-related pathways, such as focal adhesion and adherent junction (13) (Fig. S8C) with *KIF5B*, *BRAF*, *KRAS* and *ERBB2*, *EGFR*, *BRAF*, *MET* as representatives (Data S6.6). When it comes to the specificity, we detect a pathway called platinum drug resistance for Lung_P (Fig. S8C). This coincides the evidence that platinum resistance limits the clinical application of platinum compounds, the most active anticancer agents used for lung cancer (14). Fluid shear stress is another specific pathway (Fig. S8C), implying cancer cells' preparation for vascular invasion and and their ability to overcome fluid shear stress in blood vessels (15).

After arriving at the microenvironment of brain, tumor cells develop more mutations involved in cerebral diseases, which raises some doubts about brain metastases increasing the risk of central nervous system (CNS) disorders. Similar to results of

Colorectal_Liver and Liver_P, specificity of Lung_Brain outweighs that of Brain_P, among which we focus on the FoxO signaling, because this pathway is the only intersection among the four results in Fig. S8C, D. The function of FoxO signaling is controlling aging and EMT of cancer cells (16, 17). As a whole, it should be responsible for the imbalanced state of NSCLC and brain.

At the level of genes, we summarize that *STK11*, *PIK3CA* and *ATM* come into play at the stage of primary NSCLC, and when cerebral metastases occur, *AKT3* and *IL7R* join in the effort; *BRAF* plays the bridge role which predestines the metastatic tendency and adaption from NSCLC to brain (Fig. S8B).

#### 2.4 Supplement to Part 5.

***Comparison of esophagogastric and Gastrointestinal stromal tumor as well as non-melanoma melamona and skin cancer.*** We also consider esophagogastric carcinoma and gastrointestinal stromal tumors of similar function as well as melanoma and non-melanoma skin cancer of similar location. Unexpectedly, the two groups have totally different performances. The former only has similarity but no specificity, while the latter is completely opposite.

With the former, less common signaling pathways between Esophagogastric_M and GastrointestinalStromal_M means dramatic changes in late-stage metastatic patients (Fig. S11A and Data S7.36-7.37). *TP53* is both vulnerable and destructive in the digestive system, since it is identified as a driver gene for both groups and act as a guardian for tumor cell growth and metastasis (Fig. S11B, C and Data S2, Table S5.3.1-5.3.2). Although we cannot directly detect significant TGF$\beta$ signaling, two of its key members *TGFBR1* and *TGFBR2* are searched together as common factors of the two metastatic cancers (Fig. S11C, E), alerting us to its critical role in digestive tract cancer cell invasion. Despite no specific pathways for primary and metastatic groups (Fig. S11A), same signaling pathways disturbed by different mutations from both results are what we recommend as therapeutic priorities.

However, no common pathways being identified for melanoma and non-melanoma skin cancer indicates that there is a remarkable gap between the two cancers in mutation characteristics and disease treatment (Fig. S11F). Either for primary or metastatic tumors, melanoma has more complicated heterogeneity than skin cancer. Only when skin cancer as metastasis can it shows own specificity relative to melanoma, which is in accord with the unique differences between Skin_P and Skin_M in Part 1. cGMP-PKG signaling for melanoma (18) and adipocytokine signaling for skin cancer (19) (Fig. S11F, H, J) can be envisaged as therapeutic targets for respective treatments. *GAN11* and *GNAQ* have a relatively wide range of mutation rates in melanoma patients with a mutually exclusive pattern (Fig. S11G, I and Data S2, Table S5.4.1-5.4.2). Their prevailing in uveal melanomas has been reported extensively (20, 21).

### Supplementary Figures

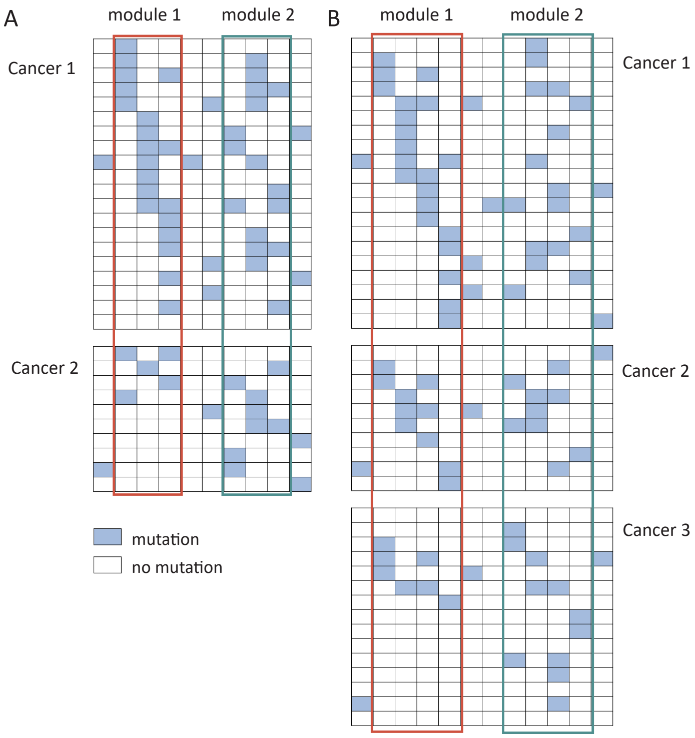


**Fig. S1.** Schematic diagram of Sim2 and Sim3 display the identification advantage of EntCDP, simulating (A) two cancers with large differences in sample size and (B) three cancers with uniform sample size. Each grid corresponds to a gene in a sample. Module 2 is more likely to be a common signaling pathway among all cancers than module 1.

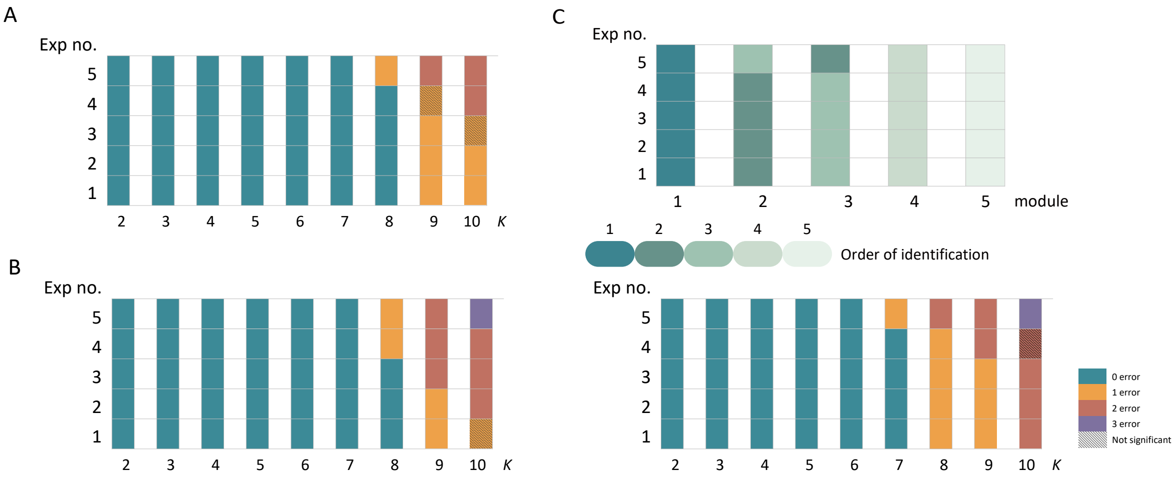


**Fig. S2.** (A, B) Results of Sim5 and 6. For $k = 2$ to 10 with a constant mutation rate in case of (A) large differences in sample size and (B) adding 2% (left) and 4% (right) 'double mutation' pattern. Each small color block illustrates one experiment and different colors represent different identification accuracy. Blocks covered by an oblique line indicate that the gene set identified by EntCDP cannot pass the permutation test. (C) Results of Sim8 for identifying five modules with $k = 5$ and decreasing proportion of mutated samples in case of large differences in sample size. The green series indicates the full identification of the five genes in each module, and the color from dark to light implies the recognition order from front to back. Exp no.: number of experiments. $k$: number of genes in each module.

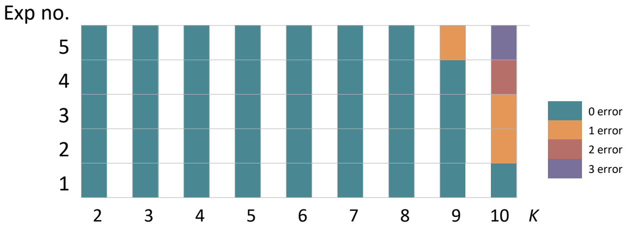


**Fig. S3.** Results of Sim11 for $k = 2$ to 10 with a constant mutation rate in case of uniform sample size. Refer to Fig. S2 for illustration.

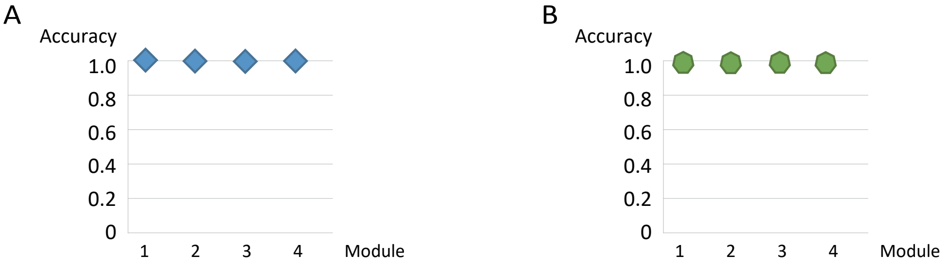


**Fig. S4.** Results of Sim12 and 13 for identifying 4 modules with $k = 5$ and decreasing proportion of mutated samples in case of (A) 1 vensus 3 and (B) 2 vensus 5 types of simulated cancers. Quadrilateral (A) and heptagon (B) are used to label the number of times the ModSDP completely identifies embedded modules in order.

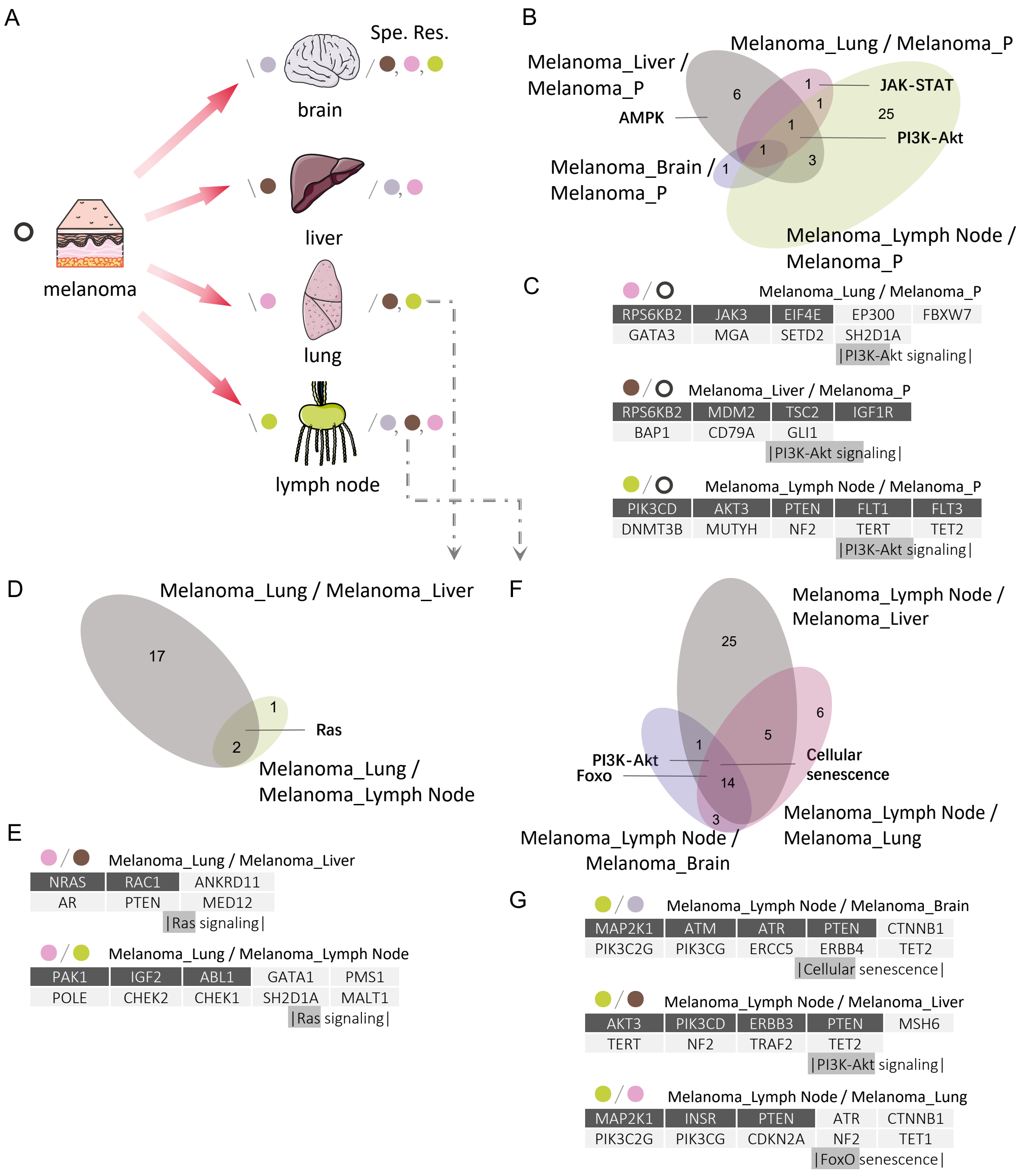


**Fig. S5.** Annotations of symbols for melanoma developing other metastases are similar to those in text Fig. 3.

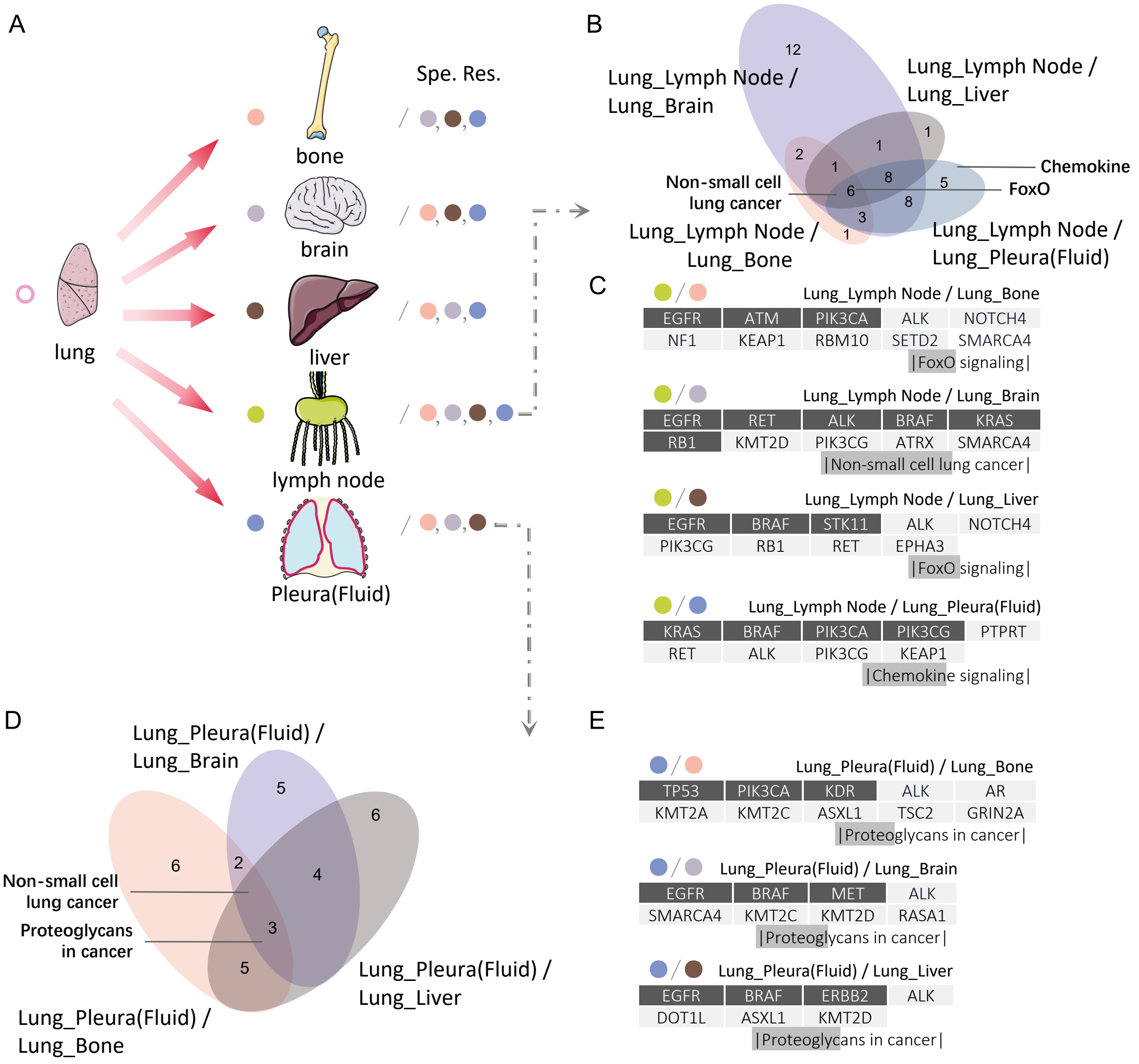


**Fig. S6.** Annotations of symbols for NSCLC developing other metastases are similar to those in text Fig. 3.

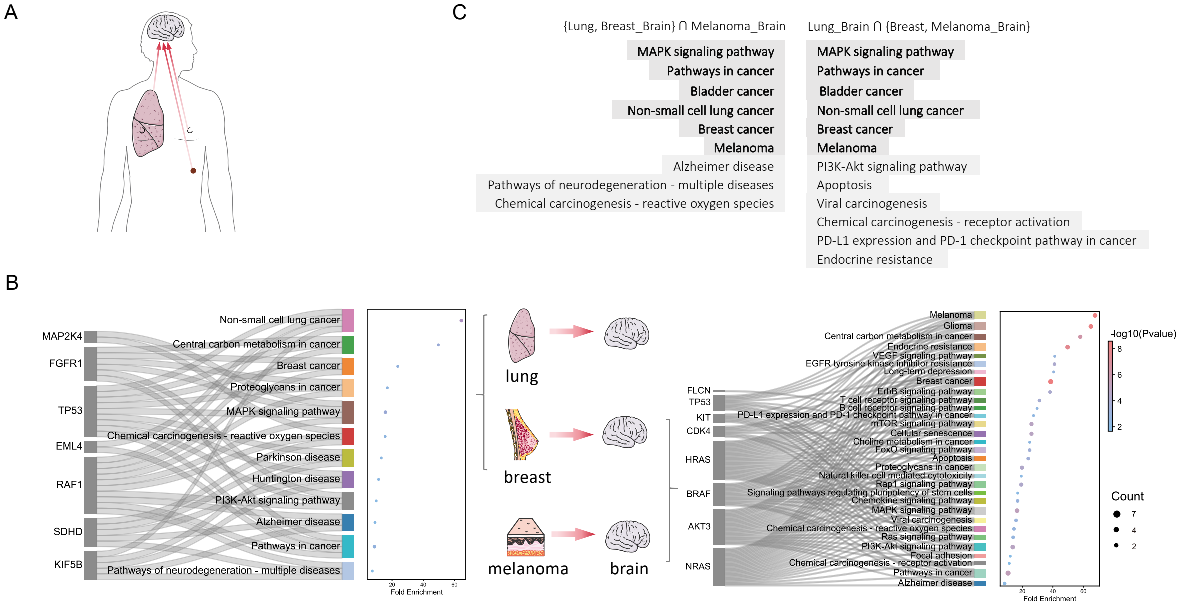


**Fig. S7.** Annotations of symbols for brain metastases are similar to those in text Fig. 4.

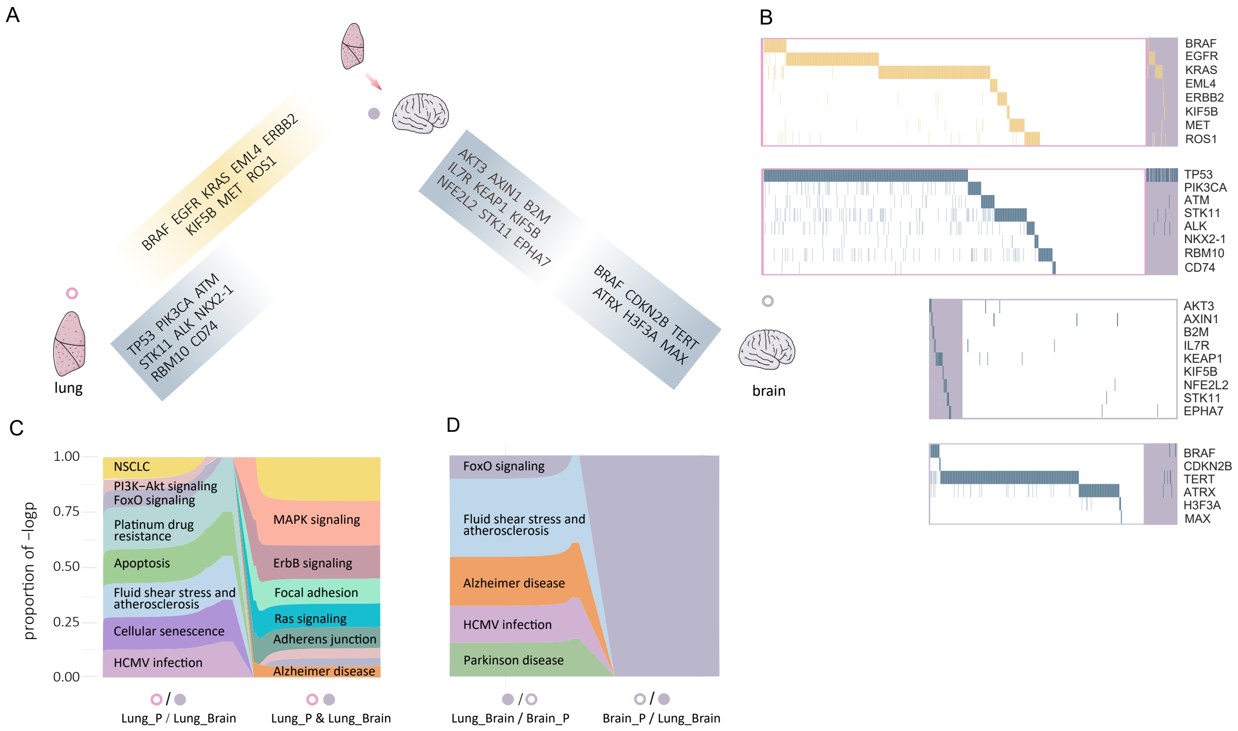


**Fig. S8.** Annotations of symbols for NSCLC with brain metastases are similar to those in text Fig. 5.

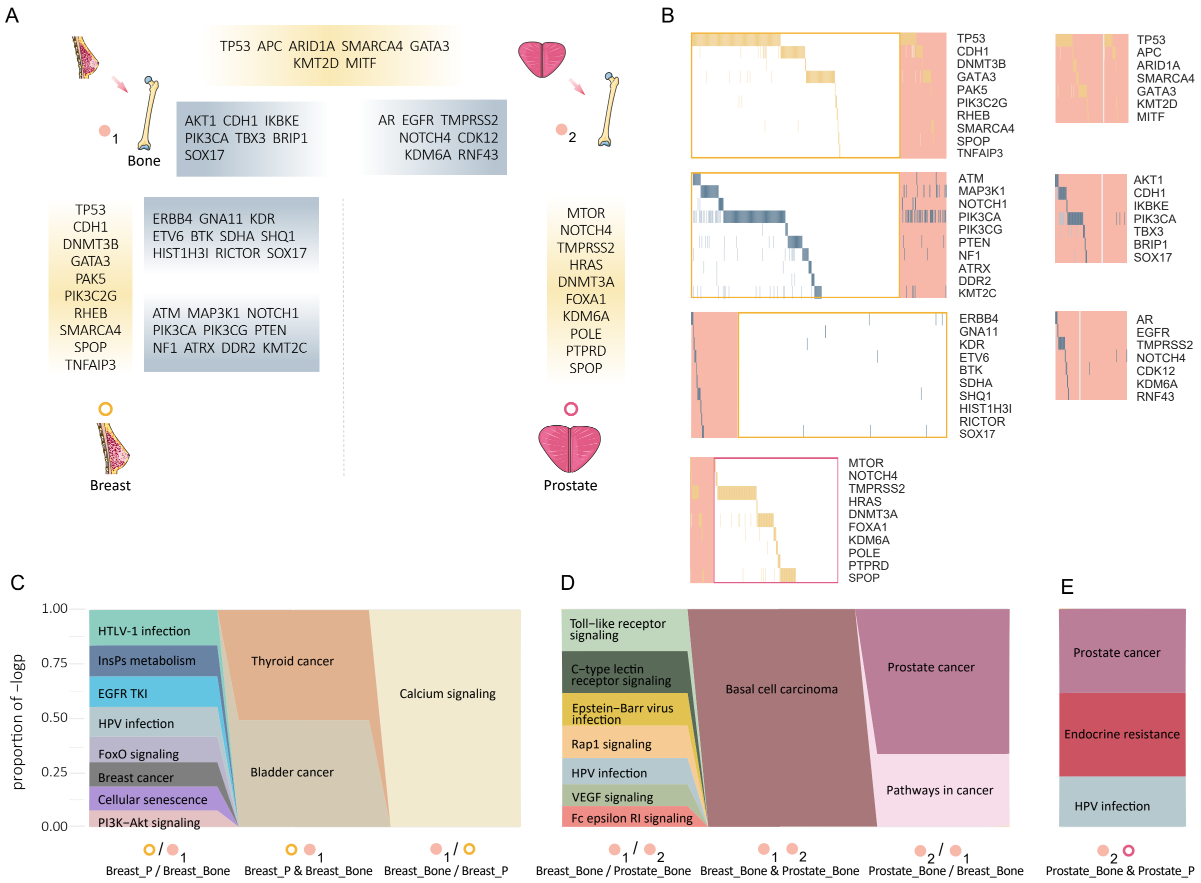


**Fig. S9.** Annotations of symbols for breast and prostate cancers with bone metastases are similar to those in text Fig. 5. The difference is that we focus on the comparison between the two types of primary tumors and respective bone metastases, as well as between the two metastases.

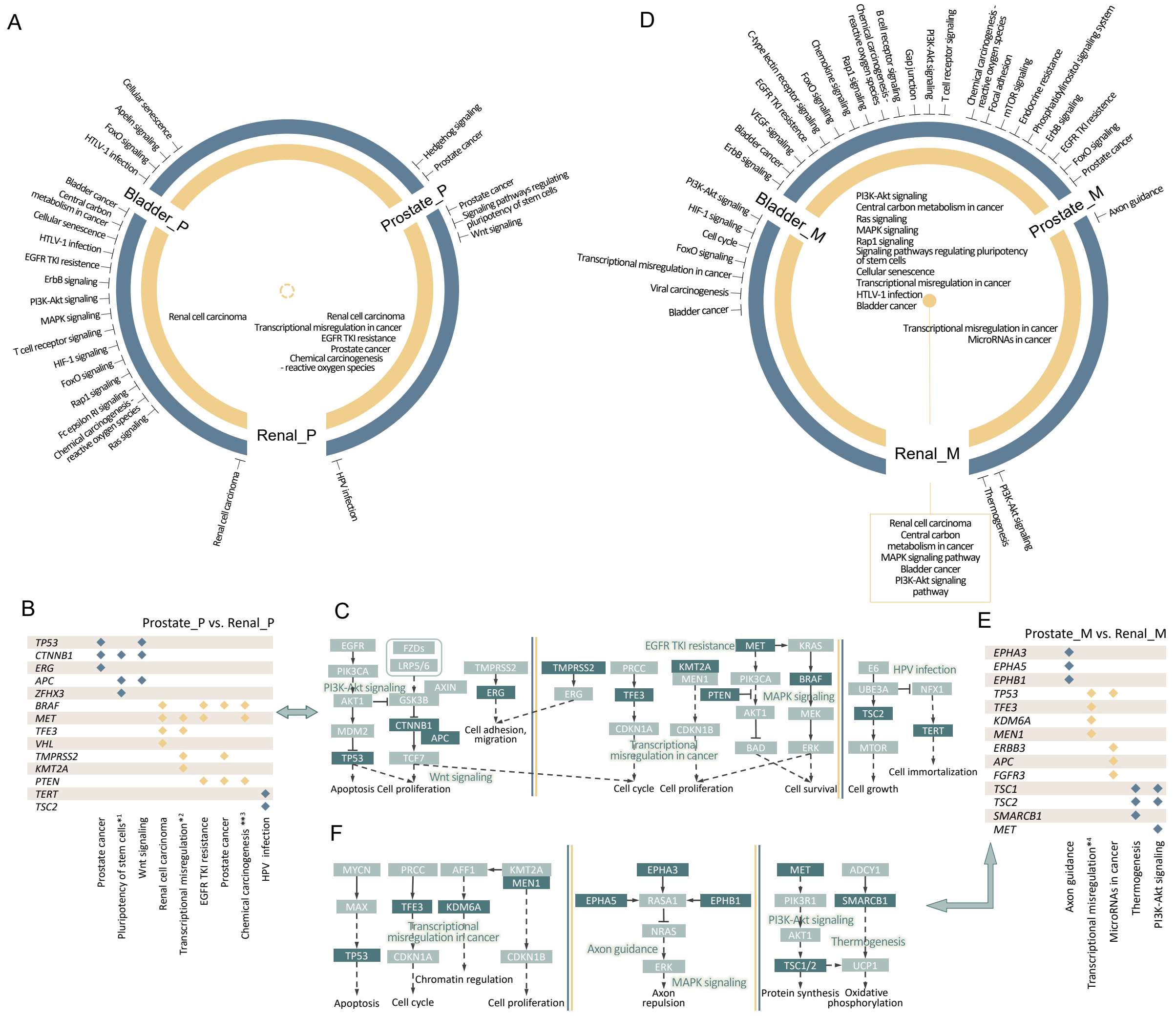

**Fig. S10.** Annotations of symbols for three male cancers are similar to those in text Fig. 6. The dotted circle at the center of a circle means there are no significant signaling pathways that the three tumors have in common. *1: Signaling pathways regulating pluripotency of stem cells; *2,*4: Transcriptional misregulation in cancer; **3: Chemical carcinogenesis - reactive oxygen species.

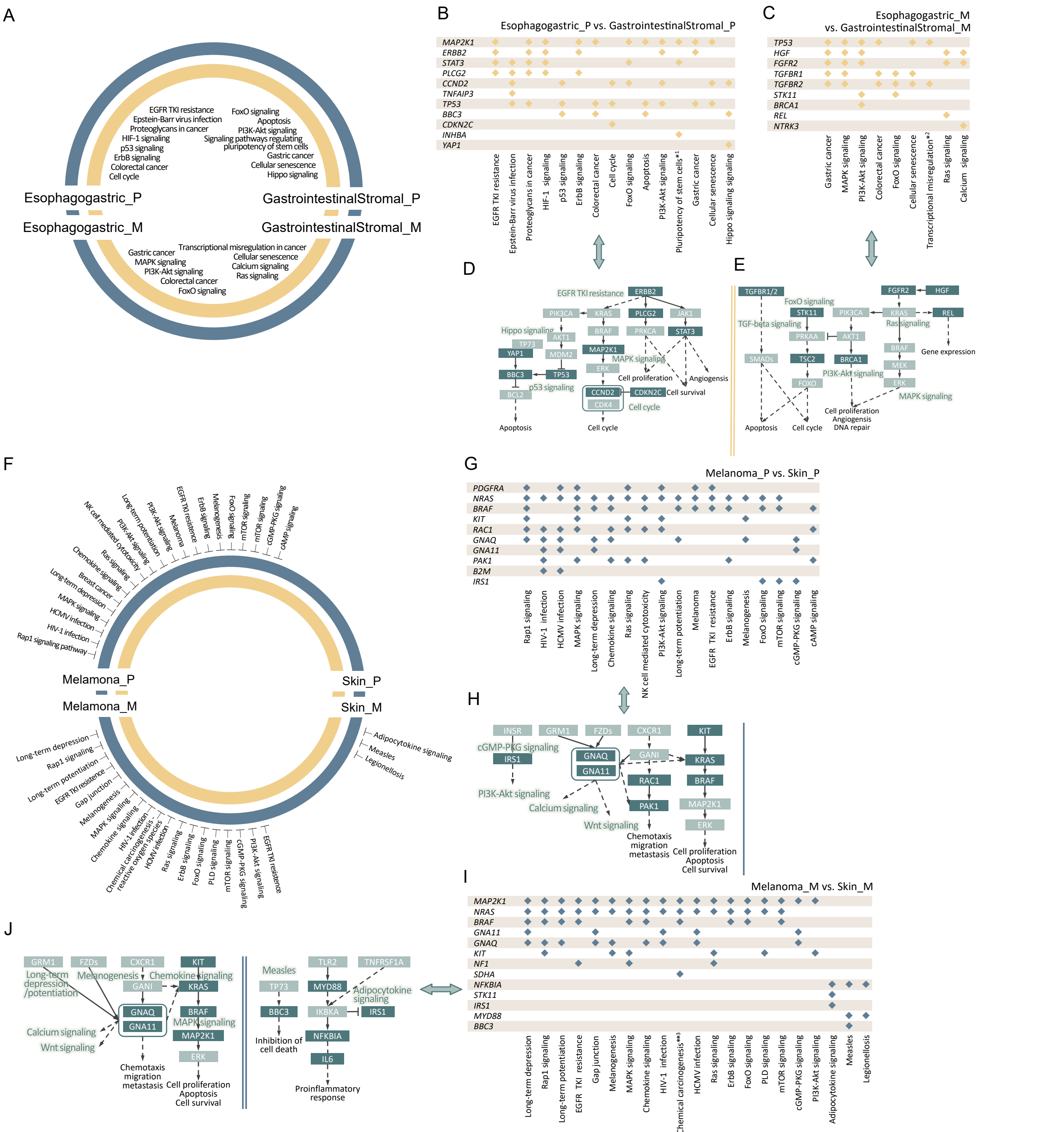

**Fig. S11.** Annotations of symbols for esophagogastric carcinoma and Gastrointestinal stromal tumor as well as melanoma and skin cancer are similar to those in text Fig. 6. *1: Signaling pathways regulating pluripotency of stem cells; *2: Transcriptional misregulation in cancer; **3: Chemical carcinogenesis - reactive oxygen species.

#### Supplementary Tables

**Table S1. Results of Sim1 $\sim$ 3**

| | ComMDP | | | EntCDP | | | | |
|---|---|---|---|---|---|---|---|---|
| Sim1 | obj_val_1[1] | 17 | ✓ | H_1[2] | 0.8886 | obj_val_1 | 15.9952 | |
| | obj_val_2 | 16 | | H_2 | 0.9997 | obj_val_2 | 15.1062 | ✓ |
| Sim2 | obj_val_1 | 17 | ✓ | H_1 | 0.9044 | obj_val_1 | 15.3745 | |
| | obj_val_2 | 16 | | H_2 | 0.9994 | obj_val_2 | 15.9842 | ✓ |
| Sim3 | obj_val_1 | 23 | ✓ | H_1 | 1.4846 | obj_val_1 | 34.1469 | |
| | obj_val_2 | 22 | | H_2 | 1.5821 | obj_val_2 | 34.8065 | ✓ |

[1] Objective function value for to module 1.
[2] Information entropy of the coverage of module 1 in two cancers.
The marks ✓correspond to the gene sets identified by ComMDP or EntCDP.

#### Other supporting materials

**SI Data S1 (biological_dataset_discription.xlsx)**

**SI Data S2 (tables_of_results.pdf)**

**SI Data S3 (part_1_enrichment_analysis.xlsx)**

**SI Data S4 (part_2_enrichment_analysis.xlsx)**

**SI Data S5 (part_3_enrichment_analysis.xlsx)**

**SI Data S6 (part_4_enrichment_analysis.xlsx)**

**SI Data S7 (part_5_enrichment_analysis.xlsx)**
