## Supplementary material for "Pan-cancer study on transition of signaling systems from primary to metastatic tumors": Data S2: tables of results

P.S.: $p_i$ is the $p$-values of the individual significance of each identified gene sets for $i^{th}$ cancer, $p$ represents the $p$-value of overall significance.

### 1 Supplementary Tables: Overall profile of primary and metastatic tumors

#### 1.1 Bladder cancer

Table S 1.1.1. Significant common driver gene set between Bladder_P and Bladder_M

| k | common gene set | p1 | p2 | q |
|---|---|---|---|---|
| 4 | *TERT,MYCN,PTPN11,SMAD3* | 0.004 | 0.029 | <0.0001 |
| 5 | *TERT,AKT2,COP1,EGFL7,PTPN11* | 0.002 | 0.004 | <0.0001 |
| 6 | *TERT,AKT2,COP1,EGFL7,PTPN11,BMPR1A* | 0.002 | 0.004 | <0.0001 |
| 7 | *TERT,AKT2,COP1,EGFL7,PTPN11,BMPR1A,HIST1H3E* | <0.0001 | 0.001 | <0.0001 |
| 8 | *TERT,AKT2,COP1,EGFL7,PTPN11,CD276,HIST1H3E,TAFA2* | <0.0001 | 0.003 | <0.0001 |
| 9 | *TERT,AKT2,COP1,EGFL7,PTPN11,CD276,EIF4E,IGF1,PIBF1* | <0.0001 | 0.002 | <0.0001 |
| 10 | *TERT,AKT2,COP1,EGFL7,PTPN11,CD79B,EIF4E,TAFA2,TNFRSF14,TNIP2* | <0.0001 | 0.004 | <0.0001 |

Table S 1.1.2. Bladder_P specific mutated driver gene sets relative to Bladder_M

| k | specific gene set | p1 | p2 | q |
|---|---|---|---|---|
| 2 | *ATRX,TERT* | 0.003 | 0.994 | 0.003 |
| 3 | *ATRX,TERT,NOTCH3* | 0.001 | 0.975 | 0.002 |
| 4 | *TP53,ATM,ERBB2,FGFR3* | <0.0001 | 0.066 | <0.0001 |
| 5 | *TP53,ATM,ERBB2,FGFR3,KRAS* | <0.0001 | 0.065 | <0.0001 |
| 6 | *TP53,CDKN1A,FGFR3,HRAS,KDM6A,NFE2L2* | <0.0001 | 0.884 | <0.0001 |
| 7 | *TP53,CDKN1A,FGFR3,HRAS,KDM6A,NFE2L2,FOXA1* | <0.0001 | 0.895 | <0.0001 |
| 8 | *TP53,CDKN1A,FGFR3,HRAS,KDM6A,NFE2L2,FOXA1,KRAS* | <0.0001 | 0.799 | <0.0001 |
| 9 | *TP53,CDKN1A,FGFR3,HRAS,KDM6A,NFE2L2,FOXA1,KRAS,FGF19* | <0.0001 | 0.822 | <0.0001 |
| 10 | *TP53,CDKN1A,FGFR3,HRAS,KDM6A,NFE2L2,FOXA1,KRAS,FGF19,MALT1* | <0.0001 | 0.784 | <0.0001 |

*Bladder_M has no specific mutated driver gene sets relative to Bladder_P when k=2 $\sim$ 10.

#### 1.2 Breast carcinoma

Table S 1.2.1. Significant common driver gene set between Breast_P and Breast_M

| k | common gene set | p1 | p2 | q |
|---|---|---|---|---|
| 2 | *TP53,GATA3* | <0.0001 | <0.0001 | <0.0001 |
| 3 | *TP53,GATA3,CDH1* | <0.0001 | <0.0001 | <0.0001 |
| 4 | *TP53,GATA3,CDH1,ESR1* | <0.0001 | <0.0001 | <0.0001 |
| 5 | *TP53,GATA3,CDH1,CDC73,CDKN1B* | <0.0001 | <0.0001 | <0.0001 |
| 6 | *TP53,GATA3,CDH1,CDC73,CDKN1B,ESR1* | <0.0001 | <0.0001 | <0.0001 |
| 7 | *TP53,GATA3,CDH1,CDC73,CDKN1B,ESR1,RHEB* | <0.0001 | <0.0001 | <0.0001 |
| 8 | *TP53,GATA3,CDH1,CDC73,CDKN1B,ESR1,RHEB,PTPN11* | <0.0001 | <0.0001 | <0.0001 |
| 9 | *TP53,GATA3,CDH1,CDC73,CDKN1B,RHEB,CCNE1,PTPN11,SMARCA4* | <0.0001 | <0.0001 | <0.0001 |
| 10 | *TP53,GATA3,CDH1,CDC73,CDKN1B,RHEB,CCNE1,PTPN11,SMARCA4,NFE2L2* | <0.0001 | <0.0001 | <0.0001 |

*Breast_M has no specific mutated driver gene sets relative to Breast_P when k=2 $\sim$ 10.
*Breast_P has no specific mutated driver gene sets relative to Breast_M when k=2 $\sim$ 10.

#### 1.3 Colorectal cancer

Table S 1.3.1. Significant common driver gene set between Colorectal_P and Colorectal_M

| k | common gene set | p1 | p2 | q |
|---|---|---|---|---|
| 2 | *APC,RNF43* | <0.0001 | <0.0001 | <0.0001 |
| 3 | *APC,RNF43,H3F3A* | <0.0001 | <0.0001 | <0.0001 |
| 4 | *APC,RNF43,H3F3A,PIM1* | <0.0001 | <0.0001 | <0.0001 |
| 5 | *APC,RNF43,PIM1,HIST1H3H,PPP6C* | <0.0001 | <0.0001 | <0.0001 |
| 6 | *APC,RNF43,ASB17,HIST1H3H,CEBPA,PPP6C* | <0.0001 | <0.0001 | <0.0001 |
| 7 | *APC,RNF43,ASB17,HIST1H3H,PIM1,H3F3A,YTHDC1* | <0.0001 | <0.0001 | <0.0001 |
| 8 | *APC,RNF43,ASB17,HIST1H3H,PIM1,PPP6C,YTHDC1,CUL1* | <0.0001 | <0.0001 | <0.0001 |
| 9 | *APC,RNF43,ASB17,HIST1H3H,PIM1,EIF4A2,YTHDC1,FOXA1,H3F3A* | <0.0001 | <0.0001 | <0.0001 |
| 10 | *APC,RNF43,RAB35,HIST1H3H,PIM1,EIF4A2,SDHAF2,FOXA1,H3F3A,SDHD* | <0.0001 | <0.0001 | <0.0001 |

Table S 1.3.2. Colorecta_M specific mutated driver gene sets relative to Colorectal_P

| k | Specific gene set | p1 | p2 | q |
|---|---|---|---|---|
| 2 | *TP53,AXL* | 0.007 | 0.604 | 0.047 |
| 4 | *TP53,AXL,RBM10,RUNX1* | <0.0001 | 0.919 | <0.0001 |
| 5 | *TP53,BRAF,RBM10,BRD4,HNF1A* | 0.012 | 0.999 | <0.0001 |
| 6 | *TP53,BRAF,RBM10,BRD4,HNF1A,NBN* | 0.005 | 0.999 | <0.0001 |
| 7 | *TP53,BRAF,RBM10,BRD4,HNF1A,NBN,NRAS* | 0.003 | 1 | <0.0001 |
| 8 | *TP53,BRAF,RBM10,BRD4,HNF1A,NBN,NRAS,RUNX1* | 0.002 | 1 | <0.0001 |
| 9 | *TP53,BRAF,RBM10,BRD4,HNF1A,NBN,NRAS,RUNX1,STK11* | <0.0001 | 1 | <0.0001 |
| 10 | *TP53,BRAF,RBM10,BRD4,HNF1A,NBN,NRAS,RUNX1,BMPR1A,NEGR1* | <0.0001 | 1 | <0.0001 |

*Colorectal_P has no specific mutated driver gene sets relative to Colorectal_M when k=2 $\sim$ 10.

#### 1.4 Endometrial cancer

Table S 1.4.1. Significant common driver gene set between Endometrial_P and Endometrial_M

| k | common gene set | p1 | p2 | q |
|---|---|---|---|---|
| 2 | *TP53,ARID1A* | <0.0001 | <0.0001 | <0.0001 |
| 3 | *TP53,AKT1,PTEN* | <0.0001 | <0.0001 | <0.0001 |
| 4 | *TP53,AKT1,PTEN,MUTYH* | <0.0001 | <0.0001 | <0.0001 |
| 5 | *TP53,AKT1,PTEN,MUTYH,CD276* | <0.0001 | <0.0001 | <0.0001 |
| 6 | *TP53,AKT1,PTEN,MUTYH,CD276,LRRC20* | <0.0001 | <0.0001 | <0.0001 |
| 7 | *TP53,AKT1,PTEN,MUTYH,CD276,LRRC20,CDKN2A* | <0.0001 | <0.0001 | <0.0001 |
| 8 | *TP53,AKT1,PTEN,MUTYH,CD276,LRRC20,CDKN2A,PAK1* | <0.0001 | <0.0001 | <0.0001 |
| 9 | *TP53,AKT1,PTEN,MUTYH,CD276,LRRC20,CDKN2A,PAK1,SMARCB1* | <0.0001 | <0.0001 | <0.0001 |
| 10 | *TP53,AKT1,PTEN,MUTYH,CD276,LRRC20,CDKN2A,PAK1,SMARCB1,HIST1H1C* | <0.0001 | <0.0001 | <0.0001 |

Table S 1.4.2. Endometrial_M specific mutated driver gene sets relative to Endometrial_P

| k | specific gene set | p1 | p2 | q |
|---|---|---|---|---|
| 3 | *CTNNB1,GLI1,KRAS* | 0.012 | 0.743 | 0.018 |
| 4 | *CTNNB1,GLI1,KRAS,PAK5* | 0.005 | 0.854 | 0.006 |
| 5 | *CTNNB1,GLI1,KRAS,XPO1,AR* | 0.007 | 0.992 | <0.0001 |
| 6 | *CTNNB1,GLI1,KRAS,XPO1,AR,RICTOR* | 0.012 | 1 | <0.0001 |

*Endometrial_P has no specific mutated driver gene sets relative to Endometrial_M when k=2 $\sim$ 10.

#### 1.5 Esophagogastric carcinoma

Table S 1.5.1. Significant common driver gene set between Esophagogastric_P and Esophagogastric_M

| k | common gene set | p1 | p2 | q |
|---|---|---|---|---|
| 4 | *TP53,PLCG2,TGFBR1,TNFAIP3* | <0.0001 | 0.03 | <0.0001 |
| 5 | *TP53,PLCG2,TGFBR1,TNFAIP3,IDH1* | <0.0001 | 0.034 | <0.0001 |
| 6 | *TP53,PLCG2,TGFBR1,TNFAIP3,IDH1,ERG* | <0.0001 | 0.026 | <0.0001 |
| 7 | *TP53,PLCG2,TGFBR1,TNFAIP3,IDH1,ERG,SMAD3* | <0.0001 | 0.01 | <0.0001 |
| 8 | *TP53,PLCG2,TGFBR1,TNFAIP3,IDH1,ERG,FGFR2,MAP2K1* | <0.0001 | 0.007 | <0.0001 |
| 9 | *TP53,PLCG2,TGFBR1,TNFAIP3,IDH1,ERG,FGFR2,SMAD3,CD276* | <0.0001 | <0.0001 | <0.0001 |
| 10 | *TP53,PLCG2,TGFBR1,TNFAIP3,IDH1,ERG,FGFR2,HGF,MAP2K1,SUFU* | <0.0001 | <0.0001 | <0.0001 |

Table S 1.5.2. Esophagogastric_P and Esophagogastric_M specific mutated driver gene sets relative to each other N

| Type | k | Specific gene set | p1 | p2 | q |
|---|---|---|---|---|---|
| **Esophagogastric_P / Esophagogastric_M** | 2 | *TP53,ERBB2* | 0.024 | 0.787 | 0.03 |
| | 3 | *TP53,ERBB2,PIK3CA* | <0.0001 | 0.79 | <0.0001 |
| | 9 | *ARID1A,BRAF,CDKN2A,CTNNB1,EPHB1,KRAS,NTRK3,RHOA,SMAD4* | 0.045 | 0.999 | 0.005 |
| | 10 | *ARID1A,BRAF,CDKN2A,CTNNB1,EPHB1,KRAS,NTRK3,RHOA,SMAD4,ANKRD11* | <0.0001 | 1 | <0.0001 |
| **Esophagogastric_M / Esophagogastric_P** | 10 | *ARAF,ARID2,DICER1,GNAS,IRS1,MYC,NKX2-1,PGR,PTCH1,TGFBR2* | 0.034 | 1 | 0.001 |

#### 1.6 Gastrointestinal stromal tumor

Table S 1.6.1. Significant common driver gene set between Gastrointestinal_P and Gastrointestinal_M

| k | common gene set | p1 | p2 | q |
|---|---|---|---|---|
| 4 | *KIT,NF1,PDGFRA,SDHA* | <0.0001 | 0.026 | <0.0001 |
| 5 | *KIT,NF1,PDGFRA,SDHA,EGFR* | <0.0001 | 0.027 | <0.0001 |
| 6 | *KIT,NF1,PDGFRA,SDHA,EGFR,B2M* | <0.0001 | 0.036 | <0.0001 |
| 7 | *KIT,NF1,PDGFRA,SDHA,EGFR,B2M,TGFBR2* | <0.0001 | 0.004 | <0.0001 |
| 8 | *KIT,NF1,PDGFRA,SDHA,EGFR,B2M,CARD11,RASA1* | <0.0001 | <0.0001 | <0.0001 |
| 9 | *KIT,NF1,PDGFRA,SDHA,EGFR,B2M,TGFBR2,RASA1,SDHB* | <0.0001 | <0.0001 | <0.0001 |
| 10 | *KIT,NF1,PDGFRA,SDHA,EGFR,B2M,TGFBR2,RASA1,SDHB,CARD11* | <0.0001 | <0.0001 | <0.0001 |

Table S 1.6.2. Gastrointestinal_P and Gastrointestinal_M specific mutated driver gene sets relative to each other

| Type | k | Specific gene set | p1 | p2 | q |
|---|---|---|---|---|---|
| **Gastrointestinal_P / Gastrointestinal_M** | 2 | *KIT,PDGFRA* | <0.0001 | 0.234 | 0.001 |
| | 3 | *KIT,PDGFRA,NF1* | <0.0001 | 0.103 | <0.0001 |
| **Gastrointestinal_M / Gastrointestinal_P** | 8 | *ANKRD11,FLT4,KRAS,MTOR,PPP2R1A,RB1,SETD2,ALK* | 0.43 | 1 | 0.002 |
| | 9 | *ANKRD11,CLCN5,DOT1L,FLT4,KRAS,MTOR,PPP2R1A,RB1,SETD2* | 0.027 | 1 | <0.0001 |
| | 10 | *ANKRD11,CLCN5,DOT1L,FLT4,KRAS,MTOR,PPP2R1A,RB1,SETD2,PHOX2B* | 0.013 | 1 | <0.0001 |

#### 1.7 Head and Neck carcinoma

Table S 1.7.1. Significant common driver gene set between HeadNeck_P and HeadNeck_M

| k | common gene set | p1 | p2 | q |
|---|---|---|---|---|
| 4 | *TP53,PIK3CA,PIK3R1,PTEN* | 0.028 | 0.026 | 0.002 |
| 5 | *TP53,PIK3CA,PIK3R1,PTEN,AKT1* | 0.01 | 0.008 | 0.001 |
| 6 | *TP53,PIK3CA,PIK3R1,PTEN,AKT1,CTCF* | 0.015 | 0.005 | <0.0001 |
| 7 | *TP53,PIK3CA,PIK3R1,PTEN,AKT1,CTCF,EWSR1* | 0.006 | 0.001 | <0.0001 |
| 8 | *TP53,PIK3CA,PIK3R1,PTEN,AKT1,CTCF,EWSR1,B2M* | 0.002 | 0.015 | <0.0001 |
| 9 | *TP53,U2AF1,PIK3R1,PTEN,AKT1,CTCF,EWSR1,ARID1A,RAF1* | 0.002 | <0.0001 | <0.0001 |
| 10 | *TP53,U2AF1,PIK3R1,PTEN,AKT1,CTCF,EWSR1,ARID1A,RAF1,AXIN1* | 0.001 | <0.0001 | <0.0001 |

Table S 1.7.2. HeadNeck_P and HeadNeck_M specific mutated driver gene sets relative to each other

| Type | k | Specific gene set | p1 | p2 | q |
|---|---|---|---|---|---|
| **HeadNeck_P / HeadNeck_M** | 2 | *ARID1A,TERT* | 0.045 | 1 | 0.004 |
| | 3 | *ARID1A,TERT,AXIN2* | 0.004 | 1 | <0.0001 |
| | 4 | *ARID1A,TERT,AXIN2,MYCN* | 0.005 | 1 | <0.0001 |
| | 6 | *ARID1A,TERT,AXIN2,MYCN,SYK,RAF1* | <0.0001 | 0.996 | <0.0001 |
| | 7 | *AXIN2,TERT,BCL6,MITF,MYCN,SYK,SETD2* | 0.001 | 1 | <0.0001 |
| | 8 | *AXIN2,TERT,BCL6,MITF,MYCN,SYK,SETD2,PLK2* | <0.0001 | 1 | <0.0001 |
| | 9 | *AXIN2,TERT,BCL6,MITF,MYCN,SYK,SETD2,PLK2,CDKN1B* | <0.0001 | 1 | <0.0001 |
| | 10 | *AXIN2,TERT,BCL6,MITF,MYCN,SYK,SETD2,PLK2,CDKN1B,ATF1* | <0.0001 | 1 | <0.0001 |
| **HeadNeck_M / HeadNeck_P** | 7 | *ATR,NFKBIA,RB1,MSH2,BMPR1A,ERBB3,KDM6A* | 0.034 | 0.999 | 0.002 |
| | 9 | *ATR,NFKBIA,RB1,BLM,KMT2A,PIK3CA,PIK3CD,SMARCA4,WT1* | 0.032 | 1 | <0.0001 |
| | 10 | *ATR,NFKBIA,RB1,BLM,BMPR1A,PIK3CA,PIK3CD,SMARCA4,WT1,KMT2A* | 0.018 | 1 | 0.001 |

#### 1.8 Non-small-cell Lung cancer

Table S 1.8.1. Significant common driver gene set between Lung_P and Lung_M

| k | common gene set | p1 | p2 | q |
|---|---|---|---|---|
| 2 | *TP53,KRAS* | <0.0001 | <0.0001 | <0.0001 |
| 3 | *TP53,KRAS,EML4* | <0.0001 | <0.0001 | <0.0001 |
| 4 | *TP53,KRAS,EML4,CD74* | <0.0001 | <0.0001 | <0.0001 |
| 5 | *TP53,KRAS,EML4,CD74,CCDC6* | <0.0001 | <0.0001 | <0.0001 |
| 6 | *TP53,KRAS,EML4,CD74,CCDC6,KIF5B* | <0.0001 | <0.0001 | <0.0001 |
| 7 | *TP53,KRAS,EML4,CD74,CCDC6,KIF5B,EIF1AX* | <0.0001 | <0.0001 | <0.0001 |
| 8 | *TP53,KRAS,EML4,CD74,CCDC6,KIF5B,EIF1AX,SLC1A2* | <0.0001 | <0.0001 | <0.0001 |
| 9 | *TP53,KRAS,EML4,CD74,CCDC6,KIF5B,EIF1AX,SLC1A2,DYSF* | <0.0001 | <0.0001 | <0.0001 |
| 10 | *TP53,KRAS,EML4,CD74,CCDC6,KIF5B,EIF1AX,SLC1A2,NCOA4,NDUFAF6* | <0.0001 | <0.0001 | <0.0001 |

Table S 1.8.2. Lung_P specific mutated driver gene sets relative to Lung_M

| k | Specific gene set | p1 | p2 | q |
|---|---|---|---|---|
| 6 | *EGFR,KMT2D,PDGFRA,RBM10,STK11,ROS1* | <0.0001 | 0.061 | <0.0001 |
| 8 | *EGFR,KMT2D,PDGFRA,RBM10,STK11,RASA1,BRAF,ROS1* | <0.0001 | 0.052 | <0.0001 |
| 9 | *EGFR,KMT2D,PDGFRA,RBM10,STK11,RASA1,BRAF,ROS1,NKX2-1* | <0.0001 | 0.096 | <0.0001 |
| 10 | *EGFR,KMT2D,PDGFRA,RBM10,STK11,RASA1,BRAF,ROS1,NKX2-1,AR* | <0.0001 | 0.388 | <0.0001 |

*Lung_M has no specific mutated driver gene sets relative to Lung_P when k=2 $\sim$ 10.

#### 1.9 Melanoma

Table S 1.9.1. Significant common driver gene set between Melanoma_P and Melanoma_M

| k | common gene set | p1 | p2 | q |
|---|---|---|---|---|
| 3 | *GNA11,GNAQ,TERT* | <0.0001 | 0.021 | <0.0001 |
| 4 | *GNA11,GNAQ,NRAS,BRAF* | <0.0001 | 0.002 | <0.0001 |
| 5 | *GNA11,GNAQ,NRAS,BRAF,KIT* | <0.0001 | <0.0001 | <0.0001 |
| 6 | *GNA11,GNAQ,NRAS,BRAF,KIT,RICTOR* | <0.0001 | <0.0001 | <0.0001 |
| 7 | *GNA11,GNAQ,NRAS,BRAF,KIT,CHEK2,MAP3K1* | <0.0001 | <0.0001 | <0.0001 |
| 8 | *GNA11,GNAQ,NRAS,BRAF,KIT,CHEK2,HNF1A,PIM1* | <0.0001 | <0.0001 | <0.0001 |
| 9 | *GNA11,GNAQ,NRAS,BRAF,KIT,CHEK2,HNF1A,PIM1,RICTOR* | <0.0001 | <0.0001 | <0.0001 |
| 10 | *GNA11,GNAQ,NRAS,BRAF,KIT,CHEK2,HNF1A,PIM1,RICTOR,PAK1* | <0.0001 | <0.0001 | <0.0001 |

Table S 1.9.2. Melanoma_M specific mutated driver gene sets relative to Melanoma_P

| k | Specific gene set | p1 | p2 | q |
|---|---|---|---|---|
| 2 | *ATRX,TERT* | 0.019 | 0.997 | <0.0001 |
| 3 | *BRAF,NF1,NRAS* | <0.0001 | 0.905 | <0.0001 |
| 4 | *ATRX,TERT,BAP1,TRAF2* | <0.0001 | 0.993 | <0.0001 |
| 5 | *ATRX,TERT,BAP1,TRAF2,PPP6C* | <0.0001 | 0.997 | <0.0001 |
| 6 | *ATRX,TERT,BAP1,TRAF2,CASP8,PTEN* | <0.0001 | 0.997 | <0.0001 |
| 7 | *ATRX,TERT,BAP1,TRAF2,CASP8,PTEN,DICER1* | 0.002 | 0.999 | <0.0001 |
| 8 | *ATRX,TERT,BAP1,TRAF2,CASP8,PTEN,DICER1,SETD2* | 0.012 | 1 | 0.001 |

*Melanoma_P has no specific mutated driver gene sets relative to Melanoma_M when k=2 $\sim$ 10.

#### 1.10 Ovarian cancer

Table S 1.10.1. Significant common driver gene set between Ovarian_P and Ovarian_M

| k | common gene set | p1 | p2 | q |
|---|---|---|---|---|
| 2 | *TP53,ARID1A* | <0.0001 | 0.005 | <0.0001 |
| 3 | *TP53,ARID1A,KRAS* | <0.0001 | <0.0001 | <0.0001 |
| 4 | *TP53,ARID1A,KRAS,NRAS* | <0.0001 | <0.0001 | <0.0001 |
| 5 | *TP53,ARID1A,KRAS,NRAS,PAK5* | <0.0001 | <0.0001 | <0.0001 |
| 6 | *TP53,ARID1A,KRAS,NRAS,PAK5,BRAF* | <0.0001 | <0.0001 | <0.0001 |
| 7 | *TP53,ARID1A,KRAS,NRAS,PAK5,BRAF,BTK* | <0.0001 | <0.0001 | <0.0001 |
| 8 | *TP53,ARID1A,KRAS,NRAS,PAK5,BRAF,BTK,PPM1D* | <0.0001 | <0.0001 | <0.0001 |
| 9 | *TP53,ARID1A,KRAS,NRAS,PAK5,BRAF,BTK,PPM1D,EYA2* | <0.0001 | <0.0001 | <0.0001 |
| 10 | *TP53,ARID1A,KRAS,NRAS,FLCN,BRAF,BTK,PPM1D,EYA2,CDC73* | <0.0001 | <0.0001 | <0.0001 |

Table S 1.10.2. Ovarian_P and Ovarian_M specific mutated driver gene sets relative to each other

| **Type** | **k** | **Specific gene set** | **p1** | **p2** | **q** |
|---|---|---|---|---|---|
| **Ovarian_P / Ovarian_M** | 5 | *LATS2,MST1R,NF2,PIK3CA,ROS1* | 0.02 | 0.748 | 0.048 |
| | 6 | *LATS1,MST1R,NF2,PIK3CA,ROS1,ERBB3* | 0.008 | 0.759 | 0.028 |
| | 7 | *LATS1,MST1R,NF2,PIK3CA,ROS1,ERBB3,LATS2* | 0.002 | 0.666 | 0.012 |
| | 8 | *LATS1,MST1R,NF2,PIK3CA,ROS1,ERBB3,LATS2,ARID1B* | <0.0001 | 0.945 | 0.002 |
| | 9 | *LATS1,MST1R,NF2,PIK3CA,ROS1,ERBB3,LATS2,ARID1B,ATRX* | <0.0001 | 0.982 | <0.0001 |
| | 10 | *LATS1,MST1R,NF2,PIK3CA,ROS1,ERBB3,LATS2,ARID1B,ATRX,KMT2A* | <0.0001 | 0.999 | <0.0001 |
| **Ovarian_M / Ovarian_P** | 9 | *APC,AXL,IRS2,KDM5A,MAP3K1,NF1,NOTCH1,PTPRS,SPEN* | 0.027 | 1 | <0.0001 |
| | 10 | *APC,AXL,IRS2,KDM5A,MAP3K1,NF1,NOTCH1,PTPRS,SPEN,BRD4* | 0.039 | 1 | 0.001 |

#### 1.11 Pancreatic cancer

Table S 1.11.1. Significant common driver gene set between Pancreatic_P and Pancreatic_M

| k | common gene set | p1 | p2 | q |
|---|---|---|---|---|
| 2 | *KRAS,MEN1* | <0.0001 | <0.0001 | <0.0001 |
| 3 | *KRAS,MEN1,BRAF* | <0.0001 | <0.0001 | <0.0001 |
| 4 | *KRAS,MEN1,BRAF,CTNNB1* | <0.0001 | <0.0001 | <0.0001 |
| 5 | *KRAS,MEN1,BRAF,CTNNB1,AURKB* | <0.0001 | <0.0001 | <0.0001 |
| 6 | *KRAS,MEN1,BRAF,CTNNB1,AURKB,TSC1* | <0.0001 | <0.0001 | <0.0001 |
| 7 | *KRAS,MEN1,BRAF,CTNNB1,AURKB,TSC1,FAT1* | <0.0001 | <0.0001 | <0.0001 |
| 8 | *KRAS,MEN1,BRAF,CTNNB1,AURKB,TSC1,FAT1,NTRK3* | <0.0001 | <0.0001 | <0.0001 |
| 9 | *KRAS,MEN1,BRAF,CTNNB1,AURKB,TSC1,FAT1,NTRK3,VHL* | <0.0001 | <0.0001 | <0.0001 |
| 10 | *KRAS,MEN1,BRAF,CTNNB1,AURKB,TSC1,ARAF,HIST1H1C,VHL,ZNF678* | <0.0001 | <0.0001 | <0.0001 |

Table S 1.11.2. Pancreatic_P and Pancreatic_M specific mutated driver gene sets relative to each other

| **Type** | **k** | **Specific gene set** | **p1** | **p2** | **q** |
|---|---|---|---|---|---|
| **Pancreatic_P / Pancreatic_M** | 2 | *TP53,GNAS* | 0.001 | 0.688 | 0.001 |
| | 3 | *TP53,GNAS,PTPRT* | <0.0001 | 0.974 | <0.0001 |
| | 4 | *TP53,GNAS,PTPRT,ATRX* | <0.0001 | 0.884 | <0.0001 |
| | 5 | *TP53,GNAS,PTPRT,ATRX,VHL* | <0.0001 | 0.918 | <0.0001 |
| | 6 | *TP53,GNAS,PTPRT,ATRX,VHL,KMT2C* | <0.0001 | 0.976 | <0.0001 |
| | 7 | *TP53,GNAS,PTPRT,ATRX,VHL,KMT2C,AXIN1* | <0.0001 | 0.994 | <0.0001 |
| | 8 | *TP53,GNAS,PTPRT,ATRX,VHL,KMT2C,AXIN1,RNF43* | <0.0001 | 0.993 | <0.0001 |
| | 9 | *TP53,GNAS,PTPRT,ATRX,JAK3,KMT2C,AXIN1,RNF43,PLK2* | <0.0001 | 1 | <0.0001 |
| | 10 | TP53,GNAS,PTPRT,ATRX,JAK3,KMT2C,AXIN1,RNF43,PLK2,NOTCH1 | <0.0001 | 1 | <0.0001 |
| **Pancreatic_M / Pancreatic_P** | 4 | *DAXX,FANCA,KMT2D,PIK3CA* | 0.047 | 0.953 | 0.03 |

#### 1.12 Prostate cancer

Table S 1.12.1. Significant common driver gene set between Prostate_P and Prostate_M

| k | common gene set | p1 | p2 | q |
|---|---|---|---|---|
| 2 | *SPOP,TMPRSS2* | <0.0001 | 0.004 | <0.0001 |
| 3 | *SPOP,TMPRSS2,FOXA1* | <0.0001 | <0.0001 | <0.0001 |
| 4 | *SPOP,TMPRSS2,FOXA1,AR* | <0.0001 | 0.002 | <0.0001 |
| 5 | *SPOP,TMPRSS2,FOXA1,KDM6A,PTEN* | <0.0001 | <0.0001 | <0.0001 |
| 6 | *SPOP,TMPRSS2,FOXA1,KDM6A,AR,MTOR* | <0.0001 | <0.0001 | <0.0001 |
| 7 | *SPOP,TMPRSS2,FOXA1,KDM6A,AR,MTOR,EPHA3* | <0.0001 | <0.0001 | <0.0001 |
| 8 | *SPOP,TMPRSS2,FOXA1,KDM6A,AR,MTOR,EPHA3,PGR* | <0.0001 | <0.0001 | <0.0001 |
| 9 | *SPOP,TMPRSS2,FOXA1,KDM6A,AR,MTOR,EPHA5,PGR,HRAS* | <0.0001 | <0.0001 | <0.0001 |
| 10 | *SPOP,TMPRSS2,FOXA1,KDM6A,AR,MTOR,EPHA5,PGR,HRAS,EPHA3* | <0.0001 | <0.0001 | <0.0001 |

Table S 1.12.2. Prostate_M specific mutated driver gene sets relative to Prostate_P

| k | Specific gene set | p1 | p2 | q |
|---|---|---|---|---|
| 4 | *TP53,APC,ERBB4,KMT2D* | 0.03 | 0.972 | 0.004 |
| 5 | *TP53,AR,ARID1A,FLT4,PIK3CA* | 0.014 | 0.911 | 0.005 |
| 6 | *TP53,APC,ARID1A,KMT2D,ERBB4,SOX17* | 0.022 | 0.951 | 0.01 |
| 9 | *TP53,ARID5B,EPHB1,KMT2D,NTRK2,SOX17,PIK3R1,SPEN,TET2* | 0.038 | 0.999 | 0.001 |
| 10 | *TP53,ARID5B,EPHB1,KMT2D,ERBB4,SOX17,PIK3R1,SPEN,TET2,ARID1A* | 0.014 | 0.998 | <0.0001 |

*Prostate_P has no specific mutated driver gene sets relative to Prostate_M when k=2 $\sim$ 10.

#### 1.13 Renal cell carcinoma

Table S 1.13.1. Significant common driver gene set between Renal_P and Renal_M

| k | common gene set | p1 | p2 | q |
|---|---|---|---|---|
| 2 | *NF2,VHL* | 0.01 | <0.0001 | <0.0001 |
| 3 | *NF2,VHL,MET* | 0.002 | <0.0001 | <0.0001 |
| 4 | *NF2,VHL,MET,TFE3* | <0.0001 | <0.0001 | <0.0001 |
| 5 | *NF2,VHL,MET,TFE3,TSC2* | <0.0001 | <0.0001 | <0.0001 |
| 6 | *NF2,VHL,MET,TFE3,TSC2,KMT2A* | <0.0001 | <0.0001 | <0.0001 |
| 7 | *NF2,VHL,MET,TFE3,TSC2,KMT2A,SMARCB1* | <0.0001 | <0.0001 | <0.0001 |
| 8 | *NF2,VHL,MET,TFE3,TSC2,KMT2A,SMARCB1,ELOC* | <0.0001 | <0.0001 | <0.0001 |
| 9 | *NF2,VHL,MET,TFE3,TSC2,KMT2A,B2M,ELOC,CIC* | <0.0001 | <0.0001 | <0.0001 |
| 10 | *NF2,VHL,MET,TFE3,TSC2,KMT2A,B2M,ELOC,SMARCB1,FYN* | <0.0001 | <0.0001 | <0.0001 |

Table S 1.13.2. Renal_P and Renal_M specific mutated driver gene sets relative to each other

| Type | k | Specific gene set | p1 | p2 | q |
|---|---|---|---|---|---|
| **Renal_P / Renal_M** | 7 | *ALK,PBRM1,PIK3C2G,PIK3CB,PIK3CG,MTOR,NCOR1* | 0.043 | 1 | <0.0001 |
| | 8 | *ALK,PBRM1,PIK3C2G,PIK3CB,PIK3CG,MTOR,NCOR1,PTPRT* | 0.041 | 1 | <0.0001 |
| | 9 | *ALK,PBRM1,PIK3C2G,PIK3CB,PIK3CG,MTOR,NCOR1,PTPRT,NOTCH3* | 0.01 | 1 | <0.0001 |
| | 10 | *ALK,PBRM1,PIK3C2G,PIK3CB,PIK3CG,PTPRT,AXL,KMT2C,PAK5,SPEN* | 0.001 | 1 | <0.0001 |
| **Renal_M / Renal_P** | 2 | *PTEN,BAP1* | 0.04 | 0.965 | 0.016 |
| | 6 | *HNF1A,MED12,MSH6,PTEN,SF3B1,JAK3* | 0.04 | 1 | <0.0001 |
| | 7 | *HNF1A,MED12,MSH6,PTEN,SF3B1,ATRX,AKT2* | 0.015 | 0.999 | <0.0001 |
| | 8 | *HNF1A,MED12,MSH6,PTEN,SF3B1,ATRX,CALR,RECQL4* | 0.009 | 0.999 | <0.0001 |
| | 9 | *HNF1A,MED12,MSH6,PTEN,SF3B1,ATRX,SH2B3,JAK3,KRAS* | 0.004 | 1 | <0.0001 |
| | 10 | *HNF1A,MED12,MSH6,PTEN,SF3B1,AKT2,CALR,HLA-A,RECQL4,SH2B3* | 0.001 | 1 | <0.0001 |

#### 1.14 Skin cancer, non malanoma

*There is no significant common driver gene set between Skin_P and Skin_M when k=2 ∼ 10.

Table S 1.14.1. Skin_P and Skin_M specific mutated driver gene sets relative to each other

| **Type** | **k** | **Specific gene set** | **p1** | **p2** | **q** |
|---|---|---|---|---|---|
| **Skin_P / Skin_M** | 5 | *ANKRD11,BCOR,FLT3,MSH2,SPOP* | 0.044 | 0.999 | 0.001 |
| | 9 | *ANKRD11,BCOR,FLT3,MSH2,H3F3A,IKBKE,TBL3,WT1,CSF3R* | 0.028 | 1 | <0.0001 |
| | 10 | *ANKRD11,BCOR,FLT3,MSH2,H3F3A,IKBKE,TBL3,WT1,CSF3R,CDKN2B* | 0.011 | 1 | <0.0001 |
| **Skin_M / Skin_P** | 8 | *FBXW7,FGFR1,ICOSLG,IRF4,MED12,MYD88,STAG2,VTCN1* | 0.039 | 1 | 0.005 |
| | 9 | *FBXW7,FGFR1,ICOSLG,IRF4,MED12,MYD88,STAG2,VTCN1,SDHAF2* | 0.031 | 1 | 0.004 |
| | 10 | *FBXW7,FGFR1,ICOSLG,IRF4,MED12,MYD88,STAG2,VTCN1,SDHAF2,NFKBIA* | 0.019 | 1 | 0.002 |

#### 1.15 Thyroid cancer

Table S 1.15.1. Significant common driver gene set between Thyroid_P and Thyroid_M

| k | common gene set | p1 | p2 | q |
|---|---|---|---|---|
| 3 | *KRAS,RET,TERT* | 0.03 | <0.0001 | <0.0001 |
| 4 | *KRAS,RET,TERT,ARID1A* | 0.007 | <0.0001 | <0.0001 |
| 5 | *KRAS,RET,TERT,ARID1A,MAP2K4* | 0.004 | <0.0001 | <0.0001 |
| 6 | *KRAS,RET,BRAF,HRAS,NRAS,NF1* | <0.0001 | <0.0001 | <0.0001 |
| 7 | *KRAS,RET,BRAF,HRAS,NRAS,NF1,PTEN* | <0.0001 | <0.0001 | <0.0001 |
| 8 | *KRAS,RET,BRAF,HRAS,NRAS,NF1,PTEN,TP63* | <0.0001 | <0.0001 | <0.0001 |
| 9 | *KRAS,RET,BRAF,HRAS,NRAS,RB1,PTEN,PTPRT,TBX3* | <0.0001 | <0.0001 | <0.0001 |
| 10 | *KRAS,RET,BRAF,HRAS,NRAS,NF1,DAXX,PTPRT,APC,STK11* | <0.0001 | <0.0001 | <0.0001 |

Table S 1.15.2. Thyroid_P and Thyroid_M specific mutated driver gene sets relative to each other

| Type | k | Specific gene set | p1 | p2 | q |
|---|---|---|---|---|---|
| **Thyroid_P / Thyroid_M** | 7 | *CREBBP,EIF1AX,ERCC4,MAX,NF2,NFE2L2,KMT2C* | 0.031 | 0.96 | 0.009 |
| | 9 | *CREBBP,EIF1AX,ERCC4,MAX,NF2,NFE2L2,CDKN1B,DICER1,DIS3* | 0.014 | 0.996 | <0.0001 |
| | 10 | *CREBBP,EIF1AX,BCOR,PTEN,GRIN2A,NFE2L2,HIST1H3H,DIS3,HLA-A,SOX17* | 0.049 | 0.98 | 0.018 |
| **Thyroid_M / Thyroid_P** | 4 | *DNMT3A,BRAF,CCDC6,ETV6* | 0.031 | 0.99 | 0.002 |
| | 5 | *DNMT3A,BRAF,CCDC6,ETV6,IRS1* | 0.021 | 0.985 | 0.002 |
| | 6 | *DNMT3A,DAXX,TERT,ARID1B,PTEN,SPEN* | <0.0001 | 0.932 | <0.0001 |
| | 7 | *DNMT3A,DAXX,TERT,PIK3CA,PPP2R1A,SYK,TET2* | 0.005 | 0.998 | <0.0001 |
| | 8 | *DNMT3A,DAXX,TERT,PIK3CA,EP300,SYK,TET2,IRS1* | 0.003 | 1 | <0.0001 |
| | 9 | *DNMT3A,DAXX,TERT,PIK3CA,EP300,SYK,TET2,IRS1,NCOA4* | <0.0001 | 0.999 | <0.0001 |
| | 10 | *DNMT3A,ATM,TERT,ETV6,MSH6,PTEN,SPEN,IRS1,KMT2D,TP63* | <0.0001 | 0.983 | <0.0001 |

### 2 Supplementary Tables: Abnormal signals may predict the seeding site of metastasis

#### 2.1 Breast carcinoma

*Breast_P and other metastatic cancers from Breast*

Table S 2.1.1. Breast_P and Breast_Bone specific mutated driver gene sets relative to each other

| Type | k | Specific gene set | p1 | p2 | q |
|---|---|---|---|---|---|
| **Breast_P / Breast_Bone** | 2 | *PIK3CA,PTEN* | 0.01 | 0.503 | 0.021 |
| | 3 | *PIK3CA,PTEN,BRCA2* | <0.0001 | 0.186 | 0.007 |
| | 4 | *PIK3CA,PTEN,BRCA2,NF1* | 0.002 | 0.444 | 0.006 |
| | 10 | *PIK3CA,PTEN,ATRX,NF1,ATM,DDR2,KMT2C,MAP3K1,NOTCH1,PIK3CG* | 0.041 | 0.996 | <0.0001 |
| **Breast_Bone / Breast_P** | 10 | *BTK,ERBB4,ETV6,GNA11,HIST1H3I,KDR,RICTOR,SDHA,SHQ1,SOX17* | 0.02 | 0.999 | <0.0001 |

Table S 2.1.2. Breast_P specific mutated driver gene sets relative to Breast_Chest Wall

| k | Specific gene set | p1 | p2 | q |
|---|---|---|---|---|
| 2 | *PIK3CA,PTEN* | 0.014 | 1 | 0.006 |
| 3 | *PIK3CA,PTEN,BRCA2* | <0.0001 | 0.86 | <0.0001 |
| 4 | *PIK3CA,PTEN,BRCA2,AKT1* | <0.0001 | 0.48 | <0.0001 |
| 5 | *PIK3CA,PTEN,BRCA2,AKT1,NF1* | <0.0001 | 0.742 | <0.0001 |
| 6 | *PIK3CA,PTEN,BRCA2,AKT1,NF1,MAP3K1* | 0.028 | 0.957 | 0.012 |
| 9 | *PIK3CA,PTEN,BRCA2,AKT1,NF1,MAP3K1,ESR1,GRIN2A,KMT2C* | 0.041 | 0.949 | 0.023 |
| 10 | *PIK3CA,PTEN,BRCA2,AKT1,NF1,MAP3K1,ESR1,GRIN2A,KMT2C,NOTCH1* | 0.026 | 0.949 | 0.008 |

*Breast_Chest Wall has no specific mutated driver gene sets relative to Breast_P when k=2 $\sim$ 10.
*Breast_Lymph Node has no specific mutated driver gene sets relative to Breast_P when k=2 $\sim$ 10.

Table S 2.1.3. Breast_P and Breast_Liver specific mutated driver gene sets relative to each other

| Type | k | Specific gene set | p1 | p2 | q |
|---|---|---|---|---|---|
| **Breast_P / Breast_Liver** | 2 | *PIK3CA,PTEN* | 0.011 | 0.845 | 0.008 |
| | 3 | *PIK3CA,PTEN,BRCA2* | <0.0001 | 0.933 | <0.0001 |
| | 4 | *PIK3CA,PTEN,BRCA2,NF1* | 0.002 | 0.996 | <0.0001 |
| | 5 | *PIK3CA,PTEN,BRCA2,NF1,ATRX* | <0.0001 | 0.999 | <0.0001 |
| | 6 | *PIK3CA,PTEN,BRCA2,NF1,ATRX,ATM* | 0.001 | 1 | <0.0001 |
| | 7 | *PIK3CA,PTEN,BRCA2,NF1,ATRX,ATM,AKT1* | <0.0001 | 1 | <0.0001 |
| | 8 | *PIK3CA,PTEN,BRCA2,NF1,ATRX,ATM,AKT1,DDR2* | <0.0001 | 1 | <0.0001 |
| | 9 | *PIK3CA,PTEN,BRCA2,NF1,ATRX,ATM,AKT1,DDR2,FLT1* | <0.0001 | 0.999 | <0.0001 |
| | 10 | *PIK3CA,PTEN,BRCA2,NF1,ATRX,ATM,AKT1,DDR2,FLT1,MED12* | <0.0001 | 1 | <0.0001 |
| **Breast_Liver / Breast_P** | 3 | *ERBB2,ESR1,FGFR4* | 0.002 | 0.591 | 0.025 |
| | 4 | *ERBB2,ESR1,FGFR4,BLM* | 0.002 | 0.56 | 0.006 |
| | 5 | *ERBB2,ESR1,FGFR4,BLM,SHQ1* | <0.0001 | 0.518 | 0.002 |
| | 6 | *ERBB2,ESR1,FGFR4,BLM,SHQ1,FGFR4* | <0.0001 | 0.813 | <0.0001 |
| | 7 | *ERBB2,ESR1,FGFR4,BLM,SHQ1,FGFR4,KDR* | <0.0001 | 0.809 | <0.0001 |
| | 8 | *ERBB2,ESR1,FGFR4,BLM,SHQ1,FGFR4,KDR,CCND2* | <0.0001 | 0.802 | 0.001 |
| | 9 | *ERBB2,ESR1,EP300,BLM,GATA1,ERCC5,MAPK1,MYOD1,ROS1* | <0.0001 | 0.668 | <0.0001 |
| | 10 | *ERBB2,ESR1,EP300,BLM,SHQ1,CCND2,KDR,MAPK1,ROS1,GATA1* | <0.0001 | 0.696 | <0.0001 |

Table S 2.1.4. Breast_P and Breast_Lung specific mutated driver gene sets relative to each other

| Type | k | Specific gene set | p1 | p2 | q |
|---|---|---|---|---|---|
| Breast_P / Breast_Lung | 2 | *PIK3CA,PTEN* | 0.006 | 0.157 | 0.041 |
| | 3 | *PIK3CA,PTEN,BRCA2* | <0.0001 | 0.614 | 0.001 |
| | 4 | *PIK3CA,PTEN,BRCA2,AKT1* | <0.0001 | 0.481 | <0.0001 |
| | 5 | *PIK3CA,PTEN,BRCA2,AKT1,MAP3K1* | 0.017 | 0.944 | 0.007 |
| | 6 | *PIK3CA,PTEN,BRCA2,AKT1,MAP3K1,NF1* | 0.024 | 0.977 | 0.013 |
| | 7 | *PIK3CA,PTEN,BRCA2,AKT1,MAP3K1,NF1,ATM* | 0.012 | 0.966 | 0.004 |
| | 8 | *PIK3CA,PTEN,BRCA2,AKT1,MAP3K1,NF1,ATM,GRIN2A* | 0.007 | 0.97 | 0.004 |
| | 9 | *PIK3CA,PTEN,BRCA2,AKT1,MAP3K1,NF1,ATM,GRIN2A,MED12* | 0.003 | 0.988 | 0.003 |
| | 10 | *PIK3CA,PTEN,BRCA2,AKT1,MAP3K1,NF1,ATM,GRIN2A,MED12,DDR2* | 0.002 | 0.974 | <0.0001 |
| Breast_Lung / Breast_P | 8 | *BRIP1,DOT1L,ELOC,EPHA7,FGFR1,INSR,MST1,CSF3R* | 0.049 | 0.998 | 0.003 |
| | 9 | *BRIP1,DOT1L,ELOC,EPHA7,FGFR1,INSR,MST1,IRS1,RAD51B* | 0.044 | 1 | 0.002 |
| | 10 | *BRIP1,DOT1L,ELOC,EPHA7,FGFR1,INSR,MST1,IRS1,RAD51B,CSF3R* | 0.024 | 0.996 | 0.001 |

Table S 2.1.5. Breast_P specific mutated driver gene sets relative to Breast_Lymph Node

| k | Specific gene set | p1 | p2 | q |
|---|---|---|---|---|
| 3 | *BRCA2,PIK3CA,PTEN* | <0.0001 | 0.331 | 0.001 |
| 4 | *BRCA2,PIK3CA,PTEN,AKT1* | <0.0001 | 0.26 | <0.0001 |
| 5 | *BRCA2,PIK3CA,PTEN,AKT1,NF1* | 0.001 | 0.488 | 0.003 |
| 6 | *BRCA2,PIK3CA,PTEN,AKT1,NF1,MAP3K1* | 0.023 | 0.82 | 0.012 |
| 10 | *BRCA2,PIK3CA,PTEN,AKT1,NF1,MAP3K1,ARID1A,ARID1B,KMT2C,NOTCH1* | 0.046 | 0.999 | 0.001 |

##### *Other metastatic cancers from Breast*

Table S 2.1.6. Breast_Bone specific mutated driver gene sets relative to Breast_Chest Wall

| k | Specific gene set | p1 | p2 | q |
|---|---|---|---|---|
| 3 | *AKT1,PIK3CA,RUNX1* | 0.003 | 0.377 | 0.390 |
| 5 | *AKT1,PIK3CA,RUNX1,ESR1,RB1* | 0.014 | 0.724 | 0.013 |
| 6 | *AKT1,PIK3CA,RUNX1,ESR1,RB1,PTEN* | 0.004 | 0.735 | 0.007 |
| 7 | *AKT1,PIK3CA,RUNX1,ESR1,RB1,PTEN,ASXL1* | 0.006 | 0.751 | 0.005 |
| 8 | *DNMT1,PIK3CA,RUNX1,ESR1,RB1,PTEN,ASXL1,NSD1* | 0.001 | 0.976 | 0.001 |
| 9 | *DNMT1,PIK3CA,RUNX1,ESR1,RB1,PTEN,ASXL1,NSD1,BRIP1* | 0.001 | 0.976 | <0.0001 |
| 10 | *DNMT1,PIK3CA,RUNX1,ESR1,RB1,PTEN,ASXL1,NSD1,BRIP1,GRIN2A* | <0.0001 | 0.979 | <0.0001 |

*Breast_Chest Wall has no specific mutated driver gene sets relative to Breast_Bone when k=2 ∼ 10.

Table S 2.1.7. Breast_Bone and Breast_Liver specific mutated driver gene sets relative to each other

| Type | k | Specific gene set | p1 | p2 | q |
|---|---|---|---|---|---|
| **Breast_Bone / Breast_Liver** | 6 | *ASXL1,BTK,CBFB,GRIN2A,IKBKE,RICTOR* | 0.019 | 0.974 | 0.026 |
| | 7 | *ASXL1,BTK,CBFB,GRIN2A,IKBKE,RICTOR,PAK1* | 0.011 | 0.967 | 0.025 |
| | 8 | *ASXL1,BTK,CBFB,GRIN2A,IKBKE,RICTOR,EPHA7,FGF19* | 0.023 | 0.972 | 0.014 |
| | 9 | *ASXL1,BTK,CBFB,GRIN2A,IKBKE,RICTOR,MSH2,NTRK3,SOX17* | 0.028 | 1 | <0.0001 |
| | 10 | *ASXL1,BTK,CBFB,GRIN2A,IKBKE,RICTOR,MSH2,NTRK3,SOX17,ERBB4* | 0.011 | 0.999 | 0.001 |
| **Breast_Liver / Breast_Bone** | 10 | *BLM,ERBB3,ESR1,FAT1,KMT2C,KMT2D,NF1,NOTCH2,SMARCB1,SPEN* | 0.037 | 1 | <0.0001 |

*Breast_Lung has no specific mutated driver gene sets relative to Breast_Bone when k=2 ∼ 10.
*Breast_Chest Wall has no specific mutated driver gene sets relative to Breast_Liver when k=2 ∼ 10.
*Breast_Chest Wall has no specific mutated driver gene sets relative to Breast_Lymph Node when k=2 ∼ 10.
*Breast_Lung has no specific mutated driver gene sets relative to Breast_Liver when k=2 ∼ 10.
*Breast_Lymph Node has no specific mutated driver gene sets relative to Breast_Liver when k=2 ∼ 10.

Table S 2.1.8. Breast_Bone specific mutated driver gene sets relative to Breast_Lung

| k | Specific gene set | p1 | p2 | q |
|---|---|---|---|---|
| 8 | ERBB2,ESR1,GRIN2A,IKBKE,NCOR1,NSD1,PIK3CA,DNMT1 | 0.043 | 0.941 | 0.015 |
| 9 | ERBB2,ESR1,GRIN2A,IKBKE,NCOR1,NSD1,PIK3CA,DNMT1,BTK | 0.023 | 0.919 | 0.013 |
| 10 | ERBB2,ESR1,GRIN2A,IKBKE,NCOR1,NSD1,PIK3CA,PIK3CB,TERT,KDR | 0.003 | 0.973 | 0.002 |

Table S 2.1.9. Breast_Bone and Breast_Lymph Node specific mutated driver gene sets relative to each other

| Type | k | Specific gene set | p1 | p2 | q |
|---|---|---|---|---|---|
| Breast_Bone / Breast_Lymph Node | 4 | AKT1,BTK,ESR1,MAP3K1 | 0.049 | 0.979 | 0.013 |
| | 7 | AKT1,BTK,ESR1,MAP3K1,IKBKE,PARP1,BRCA1 | 0.027 | 1 | <0.0001 |
| | 8 | AKT1,BTK,ESR1,MAP3K1,IKBKE,PARP1,PAX5,BRCA1 | 0.011 | 1 | <0.0001 |
| | 9 | AKT1,BTK,ESR1,MAP3K1,IKBKE,FUBP1,PAX5,BRCA1,SOX17 | <0.0001 | 1 | <0.0001 |
| | 10 | AKT1,BTK,ESR1,MAP3K1,IKBKE,PARP1,PAX5,BRCA1,KLF4,RPS6KA4 | <0.0001 | 1 | <0.0001 |
| Breast_Lymph Node / Breast_Bone | 2 | TP53,IKZF1 | 0.043 | 1 | 0.013 |

Table S 2.1.10. Breast_Liver specific mutated driver gene sets relative to Breast_Chest Wall

| k | Specific gene set | p1 | p2 | q |
|---|---|---|---|---|
| 2 | ESR1,TP53 | <0.0001 | 0.149 | <0.0001 |

Table S 2.1.11. Breast_Chest Wall and Breast_Lung specific mutated driver gene sets relative to each other

| Type | k | Specific gene set | p1 | p2 | q |
|---|---|---|---|---|---|
| Breast_Chest Wall / Breast_Lung | 9 | CHEK2,ERG,FLT4,FOXA1,JAK1,KDM6A,KMT2A,TBX3,HIST1H1C | 0.035 | 1 | 0.037 |
| | 10 | CHEK2,ERG,FLT4,FOXA1,JAK1,KDM6A,KMT2A,TBX3,NOTCH3,EIF4A2 | 0.016 | 1 | 0.026 |
| Breast_Lung / Breast_Chest Wall | 9 | AKT2,AMER1,ASXL1,ATR,BRIP1,CIC,DOT1L,INSR,PTEN | 0.025 | 1 | 0.021 |
| | 10 | AKT2,AMER1,ASXL1,ATR,BRIP1,CIC,DOT1L,INSR,PTEN,TSC2 | 0.025 | 1 | 0.047 |

Table S 2.1.12. Breast_Lymph Node specific mutated driver gene sets relative to Breast_Chest Wall

| k | Specific gene set | p1 | p2 | q |
|---|---|---|---|---|
| 5 | *ERBB2,ESR1,PTEN,PIK3CA,KDM6A* | 0.036 | 0.741 | 0.044 |
| 6 | *ERBB2,ESR1,PTEN,PIK3CA,POLE,BRIP1* | 0.021 | 0.731 | 0.034 |
| 7 | *ERBB2,ESR1,PTEN,PIK3CA,POLE,BRIP1,DNMT3A* | 0.029 | 0.732 | 0.032 |
| 8 | *ERBB2,ESR1,PTEN,PIK3CA,POLE,BRIP1,DNMT3A,EPHA5* | 0.012 | 0.857 | 0.013 |
| 9 | *ERBB2,ESR1,PTEN,PIK3CA,POLE,BRIP1,DNMT3A,EPHA5,GPS2* | 0.01 | 0.906 | 0.005 |
| 10 | *ERBB2,ESR1,PTEN,PIK3CA,POLE,BRIP1,DNMT3A,EPHA5,GPS2,ARID2* | 0.004 | 0.86 | 0.007 |

Table S 2.1.13. Breast_Liver specific mutated driver gene sets relative to Breast_Lung

| k | Specific gene set | p1 | p2 | q |
|---|---|---|---|---|
| 2 | *TP53,ESR1* | <0.0001 | 0.266 | <0.0001 |
| 7 | *ARID2,EP300,ERBB2,ESR1,KMT2A,NCOR1,PIK3CA* | 0.03 | 0.965 | 0.016 |
| 8 | *ARID2,EP300,ERBB2,ESR1,KMT2A,NCOR1,PIK3CA,CARD11* | 0.012 | 0.969 | 0.007 |
| 9 | *ARID2,EP300,ERBB2,ESR1,KMT2A,NCOR1,PIK3CA,CARD11,DNMT1* | 0.01 | 0.969 | 0.006 |
| 10 | *ARID2,EP300,ERBB2,ESR1,KMT2A,NCOR1,PIK3CA,CARD11,DNMT1,BLM* | 0.006 | 0.965 | 0.003 |

Table S 2.1.14. Breast_Liver specific mutated driver gene sets relative to Breast_Lymph Node

| k | Specific gene set | p1 | p2 | q |
|---|---|---|---|---|
| 2 | *TP53,ESR1* | <0.0001 | 0.469 | <0.0001 |
| 3 | *ERBB2,ESR1,NF1* | 0.015 | 0.956 | 0.003 |
| 4 | *ERBB2,ESR1,NF1,FOXA1* | 0.043 | 0.961 | 0.01 |
| 6 | *ERBB2,ERBB3,NF1,FOXA1,DNMT1,ESR1* | 0.036 | 0.997 | 0.004 |
| 8 | *ERBB2,ERBB3,NF1,FOXA1,DNMT1,ESR1,DICER1,SHQ1* | 0.021 | 0.999 | <0.0001 |
| 9 | *ERBB2,ERBB3,NF1,FOXA1,DNMT1,ESR1,ERBB4,ERRFI1,FAT1* | 0.018 | 0.99 | 0.003 |
| 10 | *ERBB2,ERBB3,NF1,FOXA1,DNMT1,ESR1,CDKN1B,SHQ1,FAT1,PALB2* | 0.002 | 0.998 | <0.0001 |

Table S 2.1.15. Breast_Chest Wall, Liver specific mutated driver gene sets relative to Breast_Lymph Node

| k | Specific gene set | p1 | p2 | p3 | q |
|---|---|---|---|---|---|
| **10** | *AKT1,ARAF,BAP1,CDKN1B,ERBB3,ESR1,FOXA1,NOTCH3,PIK3R1,SHQ1* | 0.033 | 0.011 | 0.919 | <0.0001 |

*Breast_Lymph Node has no specific mutated driver gene sets relative to Breast_Chest Wall, Liver when k=2 ∼ 10.
*Breast_Lung has no specific mutated driver gene sets relative to Breast_Lymph Node when k=2 ∼ 10.
*Breast_Lymph Node has no specific mutated driver gene sets relative to Breast_Lung when k=2 ∼ 10.

#### 2.2 Non-small-cell Lung cancer

*Lung_P and other metastatic cancers from Lung*

Table S 2.2.1. Lung_P specific mutated driver gene sets relative to Lung_Bone

| k | Specific gene set | p1 | p2 | q |
|---|---|---|---|---|
| 2 | *TP53,KRAS* | <0.0001 | 0.055 | <0.0001 |
| 3 | *TP53,KRAS,EML4* | <0.0001 | 0.077 | <0.0001 |
| 4 | *TP53,KRAS,EML4,EGFR* | <0.0001 | 0.082 | <0.0001 |
| 7 | *TP53,ALK,ATM,CD74,PIK3CA,RBM10,STK11* | <0.0001 | 0.226 | <0.0001 |
| 8 | *TP53,ALK,ATM,CD74,PIK3CA,RBM10,STK11,NKX2-1* | <0.0001 | 0.259 | <0.0001 |
| 9 | *TP53,ALK,ATM,MED12,PIK3CA,RBM10,STK11,NKX2-1,SETD2* | <0.0001 | 0.398 | <0.0001 |
| 10 | *TP53,ALK,ATM,MED12,PIK3CA,RBM10,STK11,NKX2-1,SETD2,CTNNB1* | <0.0001 | 0.443 | <0.0001 |

*Lung_Bone has no specific mutated driver gene sets relative to Lung_P when k=2 $\sim$ 10.
*Lung_Brain has no specific mutated driver gene sets relative to Lung_P when k=2 $\sim$ 10.
*Lung_Liver has no specific mutated driver gene sets relative to Lung_P when k=2 $\sim$ 10.
*Lung_Pleura(Fluid) has no specific mutated driver gene sets relative to Lung_P when k=2 $\sim$ 10.
*Lung_Lymph Node has no specific mutated driver gene sets relative to Lung_P when k=2 $\sim$ 10.

Table S 2.2.2. Lung_P specific mutated driver gene sets relative to Lung_Brain

| k | Specific gene set | p1 | p2 | q |
|---|---|---|---|---|
| 2 | *TP53,KRAS* | <0.0001 | 0.795 | <0.0001 |
| 3 | *TP53,KRAS,EML4* | <0.0001 | 0.613 | <0.0001 |
| 4 | *TP53,EGFR,KRAS,MET* | <0.0001 | 0.614 | <0.0001 |
| 5 | *TP53,EGFR,KRAS,MET,BRAF* | <0.0001 | 0.675 | <0.0001 |
| 6 | *TP53,EGFR,KRAS,MET,BRAF,EML4* | <0.0001 | 0.465 | <0.0001 |
| 7 | *TP53,EGFR,KRAS,MET,BRAF,EML4,CD74* | <0.0001 | 0.48 | <0.0001 |
| 8 | *TP53,ALK,ATM,NKX2-1,PIK3CA,CD74,RBM10,STK11* | <0.0001 | 0.488 | <0.0001 |
| 9 | *TP53,EGFR,KRAS,MET,BRAF,EML4,CD74,ERBB2,STK11* | <0.0001 | 0.541 | <0.0001 |
| 10 | *TP53,ALK,ATM,NKX2-1,PIK3CA,RBM10,SETD2,STK11,CTNNB1,MED12* | <0.0001 | 0.832 | <0.0001 |

Table S 2.2.3. Lung_P specific mutated driver gene sets relative to Lung_Liver

| k | Specific gene set | p1 | p2 | q |
|---|---|---|---|---|
| 2 | *TP53,KRAS* | <0.0001 | 0.346 | <0.0001 |
| 3 | *TP53,KRAS,EML4* | <0.0001 | 0.189 | <0.0001 |
| 4 | *TP53,KRAS,EGFR,MET,* | <0.0001 | 0.457 | <0.0001 |
| 5 | *TP53,KRAS,EGFR,MET,EML4* | <0.0001 | 0.32 | <0.0001 |
| 6 | *TP53,KRAS,EGFR,MET,EML4,BRAF* | <0.0001 | 0.222 | <0.0001 |
| 7 | *TP53,ALK,ATM,CD74,PIK3CA,RBM10,STK11* | <0.0001 | 0.799 | <0.0001 |
| 8 | *TP53,ALK,ATM,CD74,PIK3CA,RBM10,STK11,NKX2-1* | <0.0001 | 0.771 | <0.0001 |
| 9 | *TP53,ALK,ATM,CD74,PIK3CA,RBM10,STK11,NKX2-1,MED12* | <0.0001 | 0.847 | <0.0001 |
| 10 | *TP53,ALK,ATM,CD74,PIK3CA,RBM10,STK11,NKX2-1,MED12,CTNNB1* | <0.0001 | 0.955 | <0.0001 |

Table S 2.2.4. Lung_P specific mutated driver gene sets relative to Lung_Pleura(Fluid)

| **k** | **Specific gene set** | **p1** | **p2** | **q** |
|---|---|---|---|---|
| 2 | *TP53,KRAS* | <0.0001 | 0.426 | <0.0001 |
| 3 | *TP53,KRAS,MET* | <0.0001 | 0.45 | <0.0001 |
| 4 | *TP53,KRAS,MET,EGFR* | <0.0001 | 0.098 | <0.0001 |
| 5 | *TP53,KRAS,MET,EGFR,BRAF* | <0.0001 | 0.072 | <0.0001 |
| 7 | *TP53,ALK,NKX2-1,STK11,RBM10,SMAD4,SETD2* | <0.0001 | 0.613 | <0.0001 |
| 8 | *TP53,ALK,NKX2-1,STK11,PIK3CA,RBM10,ATM,CD74* | <0.0001 | 0.081 | <0.0001 |
| 9 | *TP53,ALK,NKX2-1,STK11,PIK3CA,RBM10,ATM,SETD2,MED12* | <0.0001 | 0.312 | <0.0001 |
| 10 | *TP53,ALK,NKX2-1,STK11,PIK3CA,RBM10,ATM,SETD2,MED12,CTNNB1* | <0.0001 | 0.252 | <0.0001 |

Table S 2.2.5. Lung_P specific mutated driver gene sets relative to Lung_Lymph Node

| **k** | **Specific gene set** | **p1** | **p2** | **q** |
|---|---|---|---|---|
| 3 | *EGFR,STK11,KMT2D* | <0.0001 | 0.078 | <0.0001 |
| 4 | *EGFR,STK11,KMT2D,RBM10* | <0.0001 | 0.341 | <0.0001 |
| 5 | *EGFR,STK11,FAT1,RBM10,ROS1* | <0.0001 | 0.89 | <0.0001 |
| 6 | *EGFR,STK11,FAT1,RBM10,ROS1,BRAF* | <0.0001 | 0.798 | <0.0001 |
| 8 | *EGFR,STK11,FAT1,RBM10,ROS1,BRAF,ALK,KMT2D* | <0.0001 | 0.234 | <0.0001 |
| 9 | *TP53,STK11,AMER1,ARAF,ATM,NKX2-1,RBM10,SETD2,SMAD4* | <0.0001 | 0.99 | <0.0001 |
| 10 | *TP53,STK11,AMER1,ARAF,ATM,NKX2-1,RBM10,SETD2,SMAD4,ARID1A* | 0.001 | 0.998 | <0.0001 |

#### *Other metastatic cancers from Lung*

Table S 2.2.6. Lung_Bone and Lung_Brain specific mutated driver gene sets relative to each other

| Type | k | Specific gene set | p1 | p2 | q |
|---|---|---|---|---|---|
| **Lung_Bone / Lung_Brain** | 5 | *DNMT1,ERBB2,MET,STK11,YAP1* | 0.019 | 0.941 | 0.017 |
| | 6 | *DNMT1,ERBB4,MET,STK11,YAP1,CBL* | <0.0001 | 0.979 | <0.0001 |
| | 7 | *DNMT1,KRAS,MET,TOP1,YAP1,CBL,PAK5* | <0.0001 | 1 | <0.0001 |
| | 8 | *DNMT1,KRAS,MET,EPHA3,YAP1,CBL,PAK5,SRC* | <0.0001 | 1 | <0.0001 |
| | 9 | *DNMT1,KRAS,MET,EPHA3,YAP1,CBL,PAK5,SRC,NOTCH3* | <0.0001 | 1 | <0.0001 |
| | 10 | *DNMT1,KRAS,MET,EPHA3,YAP1,CBL,PAK5,SRC,ERCC4,IRS2* | <0.0001 | 1 | <0.0001 |
| **Lung_Brain / Lung_Bone** | 3 | *TP53,FGFR1,WT1* | 0.001 | 1 | <0.0001 |
| | 4 | *TP53,FGFR1,AR,ERBB2* | 0.004 | 1 | 0.002 |
| | 5 | *NFE2L2,ALK,ATM,PDGFRA,ZFHX3* | 0.013 | 0.943 | 0.011 |
| | 6 | *NFE2L2,ALK,ATM,PDGFRA,ZFHX3,RAD54L* | 0.002 | 0.95 | 0.002 |
| | 7 | *NFE2L2,ARID5B,CREBBP,IL7R,PIK3CD,ROS1,SETD2* | <0.0001 | 0.905 | <0.0001 |
| | 8 | *NFE2L2,ARID5B,CREBBP,IL7R,PIK3CD,ROS1,SETD2,CENPA* | <0.0001 | 0.885 | <0.0001 |
| | 9 | *NFE2L2,ARID5B,CREBBP,IL7R,PIK3CD,ROS1,SETD2,CENPA,ARID2* | <0.0001 | 0.843 | <0.0001 |
| | 10 | *NFE2L2,ARID5B,CREBBP,IL7R,PIK3CD,ROS1,SETD2,CENPA,ARID2,BTK* | <0.0001 | 0.842 | <0.0001 |

*Lung_Bone has no specific mutated driver gene sets relative to Lung_Lymph Node when k=2 $\sim$ 10.
*Lung_Brain has no specific mutated driver gene sets relative to Lung_Lymph Node when k=2 $\sim$ 10.
*Lung_Liver has no specific mutated driver gene sets relative to Lung_Lymph Node when k=2 $\sim$ 10.
*Lung_Pleura(Fluid) has no specific mutated driver gene sets relative to Lung_Lymph Node when k=2 $\sim$ 10.

Table S 2.2.7. Lung_Bone and Lung_Liver specific mutated driver gene sets relative to each other

| Type | k | Specific gene set | p1 | p2 | q |
|---|---|---|---|---|---|
| **Lung_Bone / Lung_Liver** | 3 | *EPHA3,KRAS,SRC* | 0.025 | 0.994 | <0.0001 |
| | 4 | *EPHA3,KRAS,EPHA7,KMT2D* | 0.002 | 1 | <0.0001 |
| | 5 | *EPHA3,KRAS,EPHA7,KMT2D,ATRX* | 0.001 | 1 | <0.0001 |
| | 6 | *EPHA3,KRAS,IKZF1,KDM5A,ATRX,SRC* | 0.002 | 1 | <0.0001 |
| | 7 | *EPHA3,KRAS,IKZF1,KDM5A,ATRX,SRC,MRE11* | 0.001 | 1 | <0.0001 |
| | 8 | *EPHA3,IKZF1,KRAS,KDM5A,ATRX,SRC,MRE11,STAT5A* | 0.001 | 1 | <0.0001 |
| | 9 | *EPHA3,IKZF1,KRAS,KDM5A,ATRX,SRC,MRE11,STAT5A,TET1* | <0.0001 | 1 | <0.0001 |
| | 10 | *EPHA3,IKZF1,KRAS,KDM5A,ATRX,SRC,MRE11,STAT5A,TET1,XRCC2* | <0.0001 | 1 | <0.0001 |
| **Lung_Liver / Lung_Bone** | 3 | *ATM,EGFR,TERT* | 0.003 | 0.784 | 0.015 |
| | 4 | *ATM,EGFR,TERT,BRCA2* | <0.0001 | 0.896 | 0.001 |
| | 5 | *ATM,EGFR,TERT,BRCA2,RICTOR* | <0.0001 | 0.972 | <0.0001 |
| | 6 | *ATM,EGFR,TERT,BRCA2,RICTOR,TGFBR1* | <0.0001 | 0.989 | <0.0001 |
| | 7 | *ATM,EGFR,TERT,BRCA2,RICTOR,TGFBR1,NCOR1* | <0.0001 | 0.996 | <0.0001 |
| | 8 | *ATM,EGFR,TERT,BRCA2,RICTOR,TGFBR1,NCOR1,FBXW7* | <0.0001 | 0.999 | <0.0001 |
| | 9 | *ATM,EGFR,TERT,BRCA2,RICTOR,TGFBR1,NCOR1,FBXW7,CDKN1B* | <0.0001 | 0.994 | <0.0001 |
| | 10 | *ATM,EGFR,TERT,BRCA2,RICTOR,TGFBR1,NCOR1,FBXW7,CDKN1B,HNF1A* | <0.0001 | 0.997 | <0.0001 |

Table S 2.2.8. Lung_Lymph Node specific mutated driver gene sets relative to Lung_Bone

| **k** | **Specific gene set** | **p1** | **p2** | **q** |
|---|---|---|---|---|
| 2 | *TP53,KRAS* | 0.002 | 0.067 | 0.033 |
| 3 | *TP53,KRAS,EML4* | <0.0001 | 0.077 | <0.0001 |
| 4 | *ALK,ATM,EGFR,KEAP1* | <0.0001 | 0.1 | <0.0001 |
| 5 | *ALK,ATM,EGFR,KEAP1,NF1* | <0.0001 | 0.167 | <0.0001 |
| 6 | *ALK,ATM,EGFR,KEAP1,NF1,SMARCA4* | <0.0001 | 0.398 | <0.0001 |
| 7 | *ALK,ATM,EGFR,KEAP1,NF1,SMARCA4,NOTCH4* | <0.0001 | 0.308 | <0.0001 |
| 8 | *ALK,ATM,EGFR,KEAP1,NF1,SMARCA4,NOTCH4,SETD2* | <0.0001 | 0.463 | <0.0001 |
| 9 | *ALK,ATM,EGFR,KEAP1,NF1,SMARCA4,NOTCH4,SETD2,RBM10* | <0.0001 | 0.667 | <0.0001 |
| 10 | *ALK,ATM,EGFR,KEAP1,NF1,SMARCA4,NOTCH4,SETD2,RBM10,PIK3CA* | <0.0001 | 0.748 | <0.0001 |

Table S 2.2.9. Lung_Bone and Lung_Pleura(Fluid) specific mutated driver gene sets relative to each other

| Type | k | Specific gene set | p1 | p2 | q |
|---|---|---|---|---|---|
| Lung_Bone / Lung_Pleura(Fluid) | 3 | *POLE,AR,STK11* | 0.043 | 0.997 | 0.004 |
| | 5 | *POLE,DIS3,KDM5A,KEAP1,MED12* | 0.024 | 1 | <0.0001 |
| | 6 | *POLE,DIS3,KDM5A,KEAP1,MED12,PRKN* | 0.014 | 1 | <0.0001 |
| | 7 | *POLE,DIS3,KDM5A,KEAP1,MED12,PRKN,INHBA* | 0.007 | 1 | <0.0001 |
| | 8 | *POLE,DIS3,KDM5A,KEAP1,MED12,PRKN,EPHA3,FOXA1* | 0.003 | 1 | <0.0001 |
| | 9 | *POLE,DIS3,KDM5A,KEAP1,MED12,PRKN,EPHA3,FOXA1,PMS2* | 0.002 | 1 | <0.0001 |
| | 10 | *POLE,DIS3,KDM5A,KEAP1,MED12,PRKN,EPHA3,FOXA1,TCF7L2,MYC* | <0.0001 | 1 | <0.0001 |
| Lung_Pleura(Fluid) / Lung_Bone | 2 | *EGFR,ALK* | 0.018 | 1 | 0.015 |
| | 3 | *EGFR,ALK,KMT2C* | 0.001 | 0.507 | 0.021 |
| | 4 | *EGFR,ALK,KMT2C,SMARCA4* | 0.002 | 0.72 | 0.007 |
| | 5 | *TP53,ALK,KMT2C,POLE,APC* | 0.005 | 0.963 | 0.003 |
| | 6 | *TP53,ALK,KMT2C,PIK3CA,AR,GRIN2A* | 0.011 | 0.991 | 0.001 |
| | 7 | *TP53,ALK,KMT2C,PIK3CA,AR,GRIN2A,ASXL1* | 0.007 | 0.999 | 0.001 |
| | 8 | *TP53,ALK,KMT2A,PIK3CA,AR,GRIN2A,ASXL1,KMT2C* | 0.006 | 0.999 | 0.002 |
| | 9 | *TP53,ALK,KMT2A,PIK3CA,AR,GRIN2A,ASXL1,KMT2C,BRCA1* | 0.006 | 1 | <0.0001 |
| | 10 | *TP53,ALK,KMT2A,PIK3CA,AR,GRIN2A,ASXL1,KMT2C,KDR,TSC2* | 0.001 | 1 | <0.0001 |

Table S 2.2.10. Lung_Brian and Lung_Liver specific mutated driver gene sets relative to each other

| Type | k | Specific gene set | p1 | p2 | q |
|---|---|---|---|---|---|
| **Lung_Brain / Lung_Liver** | 5 | *ALK,FAT1,INPP4B,PIK3CG,ZFHX3* | 0.001 | 1 | <0.0001 |
| | 6 | *ALK,FAT1,INPP4B,PIK3CG,ZFHX3,NOTCH4* | <0.0001 | 1 | <0.0001 |
| | 7 | *ALK,FAT1,INPP4B,PIK3CG,ZFHX3,NOTCH4,MYCN* | <0.0001 | 1 | <0.0001 |
| | 8 | *ALK,CARD11,EPHA7,PIK3CG,ZFHX3,JAK1,MAP2K1,NFE2L2* | 0.006 | 1 | <0.0001 |
| | 9 | *ALK,CARD11,EPHA7,AKT3,ZFHX3,JAK1,MAP2K1,NFE2L2,GNA11* | 0.001 | 1 | <0.0001 |
| | 10 | *ALK,CARD11,EPHA7,AKT3,ZFHX3,JAK1,MAP2K1,NFE2L2,GNA11,MDM2* | <0.0001 | 1 | <0.0001 |
| **Lung_Liver / Lung_Brain** | 3 | *ATM,EGFR,MET* | 0.003 | 0.515 | 0.035 |
| | 4 | *ATM,EGFR,MET,IL7R* | 0.008 | 0.94 | 0.006 |
| | 5 | *ATM,EGFR,MET,IL7R,JAK2* | 0.002 | 0.879 | 0.002 |
| | 6 | *ATM,EGFR,MET,IL7R,JAK2,PTCH1* | <0.0001 | 0.899 | <0.0001 |
| | 7 | *ATM,EGFR,MET,IL7R,JAK2,PTCH1,RICTOR* | <0.0001 | 0.933 | <0.0001 |
| | 8 | *ATM,EGFR,ATR,CDKN1B,PTCH1,RICTOR,MGA,PIK3C3* | <0.0001 | 0.694 | <0.0001 |
| | 9 | *ATM,EGFR,ATR,CDKN1B,PTCH1,RICTOR,MGA,PIK3C3,MET* | <0.0001 | 0.611 | <0.0001 |
| | 10 | *ATM,EGFR,ATR,CDKN1B,PTCH1,RICTOR,MGA,PIK3C3,MET,SUZ12* | <0.0001 | 0.763 | <0.0001 |

Table S 2.2.11. Lung_Lymph Node specific mutated driver gene sets relative to Lung_Brain

| k | Specific gene set | p1 | p2 | q |
|---|---|---|---|---|
| 2 | *TP53,KRAS* | <0.0001 | 0.782 | <0.0001 |
| 3 | *EGFR,KRAS,PTPRT* | <0.0001 | 0.181 | 0.004 |
| 4 | *EGFR,KRAS,PTPRT,TERT* | <0.0001 | 0.458 | <0.0001 |
| 7 | *EGFR,KRAS,ALK,BRAF,KEAP1,MET,SMARCA4* | <0.0001 | 0.377 | <0.0001 |
| 8 | *EGFR,KRAS,ALK,BRAF,KEAP1,RET,SMARCA4,RB1* | <0.0001 | 0.587 | <0.0001 |
| 9 | *EGFR,KRAS,ALK,KEAP1,RET,MET,EPHA3,ATM,BRCA2* | <0.0001 | 0.672 | <0.0001 |
| 10 | *EGFR,KRAS,ALK,BRAF,KMT2D,RET,SMARCA4,ATRX,PIK3CG,RB1* | <0.0001 | 0.547 | <0.0001 |

Table S 2.2.12. Lung_Brain and Lung_Pleura(Fluid) specific mutated driver gene sets relative to each other

| | Type | k | Specific gene set | p1 | p2 | q |
|---|---|---|---|---|---|---|
| N | **Lung_Brain / Lung_Pleura(Fluid)** | 3 | *KEAP1,NFE2L2,PTEN* | 0.024 | 0.979 | 0.006 |
| | | 7 | *ARID5B,CREBBP,DDR2,KEAP1,MAP2K1,NFE2L2,PLCG2* | 0.028 | 0.998 | 0.007 |
| | | 8 | *ARID5B,CREBBP,DDR2,KEAP1,MAP2K1,NFE2L2,PLCG2,AKT3* | 0.021 | 0.995 | 0.003 |
| | | 9 | *ARID5B,ARID1A,AXIN1,KEAP1,MAP2K1,NFE2L2,PLCG2,AKT3,IL7R* | 0.002 | 0.974 | <0.0001 |
| | | 10 | *ARID5B,CDKN2A,MDM2,PAX5,MAP2K1,NFE2L2,PLCG2,AKT3,RAD51B,NOTCH3* | 0.008 | 1 | <0.0001 |
| | **Lung_Pleura(Fluid) / Lung_Brain** | 3 | *ALK,EGFR,MET* | <0.0001 | 0.512 | 0.012 |
| | | 4 | *ALK,EGFR,MET,ATR* | <0.0001 | 0.493 | <0.0001 |
| | | 5 | *ALK,EGFR,MET,RASA1,KMT2C* | <0.0001 | 0.814 | <0.0001 |
| | | 6 | *ALK,EGFR,MET,RASA1,KMT2C,BCOR* | <0.0001 | 0.961 | <0.0001 |
| | | 7 | *ALK,EGFR,MET,RASA1,KMT2C,BCOR,SMARCA4* | <0.0001 | 0.952 | <0.0001 |
| | | 8 | *ALK,EGFR,MET,RASA1,KMT2C,KMT2D,SMARCA4,BRAF* | <0.0001 | 0.976 | <0.0001 |
| | | 9 | *ALK,EGFR,MET,RASA1,KMT2C,KMT2D,SMARCA4,BRAF,ATR* | <0.0001 | 0.986 | <0.0001 |
| | | 10 | *ALK,EGFR,MET,FOXL2,KMT2C,KMT2D,SMARCA4,BRAF,RASA1,LATS1* | <0.0001 | 0.998 | <0.0001 |

Table S 2.2.13. Lung_Lymph Node specific mutated driver gene sets relative to Lung_Liver

| k | Specific gene set | p1 | p2 | q |
|---|---|---|---|---|
| 2 | *TP53,KRAS* | <0.0001 | 0.283 | 0.007 |
| 3 | *TP53,KRAS,EML4* | <0.0001 | 0.189 | <0.0001 |
| 4 | *ALK,EGFR,STK11,PTPRD* | 0.002 | 0.16 | 0.042 |
| 5 | *ALK,EGFR,STK11,NOTCH4,PIK3CG* | <0.0001 | 0.537 | <0.0001 |
| 6 | *ALK,EGFR,STK11,NOTCH4,PIK3CG,RB1* | <0.0001 | 0.814 | <0.0001 |
| 7 | *ALK,EGFR,STK11,NOTCH4,PIK3CG,RB1,RET* | <0.0001 | 0.848 | <0.0001 |
| 8 | *ALK,EGFR,STK11,NOTCH4,PIK3CG,RB1,RET,EPHA3* | <0.0001 | 0.79 | <0.0001 |
| 9 | *ALK,EGFR,STK11,NOTCH4,PIK3CG,RB1,RET,EPHA3,BRAF* | <0.0001 | 0.8 | <0.0001 |
| 10 | *ALK,EGFR,STK11,NOTCH4,PIK3CG,RB1,RET,EPHA3,BRAF,ATRX* | <0.0001 | 0.907 | <0.0001 |

Table S 2.2.14. Lung_Liver and Lung_Pleura(Fluid) specific mutated driver gene sets relative to each other

| Type | k | Specific gene set | p1 | p2 | q |
|---|---|---|---|---|---|
| **Lung_Liver / Lung_Pleura(Fluid)** | 5 | *FBXW7,FGFR3,KEAP1,TGFBR2,EIF1AX* | 0.015 | 1 | 0.022 |
| | 7 | *FBXW7,FGFR3,KEAP1,TGFBR2,NCOR1,CTNNB1,SPEN* | 0.03 | 1 | 0.005 |
| | 8 | *FBXW7,FGFR3,KEAP1,TGFBR2,NCOR1,CTNNB1,FGFR2,RHOA* | 0.008 | 0.995 | 0.003 |
| | 9 | *FBXW7,FGFR3,KEAP1,TGFBR2,NCOR1,CTNNB1,FGFR2,RHOA,WT1* | 0.002 | 0.991 | 0.002 |
| | 10 | *FBXW7,FGFR3,CDKN2A,TGFBR1,NCOR1,CTNNB1,MGA,MSH6,SPEN,TP63* | <0.0001 | 1 | <0.0001 |
| **Lung_Pleura(Fluid) / Lung_Liver** | 3 | *ALK,EGFR,KRAS* | <0.0001 | 0.73 | 0.013 |
| | 4 | *ALK,EGFR,KRAS,KMT2D* | <0.0001 | 0.217 | 0.002 |
| | 5 | *ALK,EGFR,KRAS,KMT2D,DOT1L* | <0.0001 | 0.215 | <0.0001 |
| | 6 | *ALK,EGFR,KRAS,KMT2D,DOT1L,ASXL1* | <0.0001 | 0.617 | <0.0001 |
| | 7 | *ALK,EGFR,KRAS,KMT2D,DOT1L,ASXL1,ERBB2* | <0.0001 | 0.518 | <0.0001 |
| | 8 | *ALK,EGFR,KRAS,KMT2D,DOT1L,ASXL1,ERBB2,NF2* | <0.0001 | 0.611 | <0.0001 |
| | 9 | *ALK,EGFR,KRAS,KMT2D,DOT1L,ASXL1,ERBB2,NF2,NOTCH4* | <0.0001 | 0.818 | <0.0001 |
| | 10 | *ALK,ATR,BAP1,BRAF,CIC,NCOA3,PTPRT,SMARCD1,SOX17,U2AF1* | 0.019 | 1 | <0.0001 |

Table S 2.2.15. Lung_Lymph Node specific mutated driver gene sets relative to Lung_Pleura(Fluid)

| k | Specific gene set | p1 | p2 | q |
|---|---|---|---|---|
| 2 | *KRAS,PTPRT* | 0.021 | 0.987 | 0.002 |
| 3 | *KRAS,PTPRT,RET* | 0.008 | 0.992 | 0.001 |
| 4 | *KRAS,PTPRT,RET,TERT* | 0.009 | 1 | <0.0001 |
| 5 | *KRAS,EPHA3,RET,CDKN2A,RB1* | 0.004 | 0.971 | <0.0001 |
| 6 | *KRAS,EPHA3,RET,CDKN2A,RB1,TERT* | 0.001 | 0.997 | <0.0001 |
| 7 | *KRAS,EPHA3,RET,CDKN2A,RB1,TERT,MTOR* | 0.001 | 0.999 | <0.0001 |
| 8 | *KRAS,EPHA3,RET,CDKN2A,RB1,TERT,MTOR,TGFBR2* | <0.0001 | 0.998 | <0.0001 |
| 9 | *KRAS,PTPRT,RET,ALK,BRAF,KEAP1,PIK3CA,PIK3CG,SMAD4* | 0.018 | 0.997 | 0.002 |
| 10 | *KRAS,PTPRT,RET,ALK,BRAF,KEAP1,PIK3CA,PIK3CG,SMAD4,RB1* | 0.019 | 0.996 | <0.0001 |

#### 2.3 Melanoma

*Melanoma_P and other metastatic cancers from Melanoma*

Table S 2.3.1. Melanoma_P and Melanoma_Brain specific mutated driver gene sets relative to each other

| Type | k | Specific gene set | p1 | p2 | q |
|---|---|---|---|---|---|
| Melanoma_P / Melanoma_Brain | 3 | *TP53,GNA11,SF3B1* | 0.006 | 0.827 | 0.017 |
| | 4 | *ATRX,GNA11,NF1,SF3B1* | 0.006 | 0.9 | 0.003 |
| | 5 | *ATRX,GNA11,NF1,SF3B1,EGFR* | 0.001 | 0.947 | <0.0001 |
| | 6 | *ATRX,GNA11,NF1,SF3B1,EGFR,BAP1* | <0.0001 | 0.993 | <0.0001 |
| | 7 | *ATRX,GNA11,NF1,SF3B1,EGFR,BAP1,CTNNB1* | <0.0001 | 0.992 | <0.0001 |
| | 8 | *ATRX,GNA11,NF1,SF3B1,EGFR,BAP1,CTNNB1,PIK3R1* | <0.0001 | 0.992 | <0.0001 |
| | 9 | *ATRX,GNA11,NF1,SF3B1,EGFR,BAP1,CTNNB1,PIK3R1,RAB35* | <0.0001 | 0.994 | <0.0001 |
| | 10 | *ATRX,EIF1AX,NF1,SF3B1,EGFR,BAP1,PIK3CA,PIK3R1,PIK3CG,POLE* | <0.0001 | 2 | <0.0001 |
| Melanoma_Brain / Melanoma_P | 4 | *AXIN2,MITF,RAD51C,TSC2* | 0.035 | 1 | 0.036 |
| | 5 | *AXIN2,MAP2K2,MDM4,RAD51C,TSC2* | 0.002 | 1 | 0.004 |
| | 6 | *AXIN2,FLCN,HGF,MEN1,PALB2,SETD2* | <0.0001 | 1 | <0.0001 |
| | 7 | *AXIN2,FLCN,HGF,MEN1,PALB2,SETD2,RAD51C* | 0.001 | 1 | <0.0001 |
| | 8 | *AXIN2,FLCN,HGF,MEN1,PALB2,SETD2,RAD51C,STAT5A* | <0.0001 | 1 | <0.0001 |
| | 9 | *AXIN2,FLCN,HGF,MEN1,PALB2,SETD2,RAD51C,STAT5A,MAP3K1* | <0.0001 | 2 | <0.0001 |
| | 10 | *AXIN2,FLCN,HGF,MEN1,PALB2,SETD2,RAD51C,ANKRD11,BIRC3,MAP3K1* | <0.0001 | 2 | <0.0001 |

Table S 2.3.2. Melanoma_P and Melanoma_Liver specific mutated driver gene sets relative to each other

| Type | k | Specific gene set | p1 | p2 | q |
|---|---|---|---|---|---|
| **Melanoma_P / Melanoma_Liver** | 5 | *ATRX,EGFR,NF1,PTEN,SF3B1* | 0.003 | 0.971 | <0.0001 |
| | 6 | *ATRX,EGFR,NF1,PTEN,SF3B1,EIF1AX* | 0.001 | 0.94 | <0.0001 |
| | 7 | *ATRX,BRAF,EIF1AX,SF3B1,FGFR2,IRS2,PIK3R1* | 0.001 | 0.86 | 0.003 |
| | 8 | *ATRX,BRAF,EIF1AX,SF3B1,IRS2,KMT2D,NOTCH3,PIK3C2G* | 0.047 | 0.995 | 0.004 |
| | 9 | *ATRX,BRAF,EIF1AX,SF3B1,IRS2,KMT2D,NOTCH3,PIK3C2G,PTEN* | 0.038 | 0.995 | 0.001 |
| | 10 | *ATRX,EGFR,EIF1AX,SF3B1,FAT1,MRE11,NF1,PIK3R1,POLE,PTEN* | <0.0001 | 1 | <0.0001 |
| **Melanoma_Liver / Melanoma_P** | 5 | *BAP1,CD79A,GLI1,IGF1R,IGF1* | 0.009 | 1 | <0.0001 |
| | 6 | *BAP1,CD79A,GLI1,IGF1R,RPS6KB2,TSC2* | 0.004 | 1 | <0.0001 |
| | 7 | *BAP1,CD79A,GLI1,IGF1R,RPS6KB2,MDM2,TSC2* | 0.002 | 1 | <0.0001 |
| | 8 | *BAP1,CD79A,GLI1,IGF1R,RPS6KB2,MDM2,TSC2,SMO* | 0.003 | 1 | <0.0001 |
| | 9 | *BAP1,CD79A,GLI1,IGF1R,RPS6KB2,MDM2,TSC2,CASP8,H3F3A* | <0.0001 | 1 | <0.0001 |
| | 10 | *BAP1,CD79A,GLI1,IGF1R,RPS6KB2,MDM2,TSC2,CASP8,RAI14,GALNT11* | <0.0001 | 1 | <0.0001 |

Table S 2.3.3. Melanoma_P and Melanoma_Lung specific mutated driver gene sets relative to each other

| Type | k | Specific gene set | p1 | p2 | q |
|---|---|---|---|---|---|
| **Melanoma_P / Melanoma_Lung** | 3 | *BAP1,NF1,SF3B1* | 0.01 | 1 | 0.002 |
| | 4 | *BAP1,NF1,SF3B1,EGFR* | 0.001 | 1 | <0.0001 |
| | 5 | *BAP1,NF1,SF3B1,EGFR,ATRX* | <0.0001 | 1 | <0.0001 |
| | 6 | *BAP1,NF1,SF3B1,EGFR,ATRX,CTNNB1* | 0.001 | 1 | <0.0001 |
| | 7 | *BAP1,NF1,SF3B1,EGFR,ATRX,CTNNB1,EIF1AX* | <0.0001 | 0.999 | <0.0001 |
| | 8 | *BAP1,NF1,SF3B1,EGFR,ATRX,CTNNB1,EIF1AX,PIK3R1* | <0.0001 | 1 | <0.0001 |
| | 9 | *BAP1,NF1,SF3B1,EGFR,ATRX,CTNNB1,EIF1AX,PIK3R1,PRDM1* | <0.0001 | 1 | <0.0001 |
| | 10 | *BAP1,NF1,SF3B1,EGFR,ATRX,CTNNB1,EIF1AX,PIK3R1,PRDM1,RARA* | <0.0001 | 1 | <0.0001 |
| **Melanoma_Lung / Melanoma_P** | 7 | *EP300,GATA3,JAK3,MGA,RPS6KB2,SETD2,SH2D1A* | 0.034 | 1 | <0.0001 |
| | 8 | *EP300,GATA3,JAK3,MGA,RPS6KB2,SETD2,SH2D1A,SMAD2* | 0.023 | 1 | <0.0001 |
| | 9 | *EP300,GATA3,JAK3,MGA,RPS6KB2,SETD2,SH2D1A,EIF4E,FBXW7* | 0.039 | 1 | <0.0001 |
| | 10 | *EP300,GATA3,JAK3,MGA,RPS6KB2,SETD2,SH2D1A,EIF4E,FBXW7,PIM1* | 0.011 | 1 | <0.0001 |

Table S 2.3.4. Melanoma_P and Melanoma_Lymph Node specific mutated driver gene sets relative to each other

| Type | k | Specific gene set | p1 | p2 | q |
|---|---|---|---|---|---|
| **Melanoma_P / Melanoma_Lymph Node** | 3 | *SF3B1,GNA11,NF1* | 0.008 | 1 | <0.0001 |
| | 4 | *SF3B1,ATRX,EGFR,NF1* | 0.003 | 1 | <0.0001 |
| | 5 | *SF3B1,ATRX,EGFR,BAP1,RAB35* | <0.0001 | 0.584 | 0.001 |
| | 6 | *SF3B1,ATRX,EGFR,BAP1,EIF1AX,RAB35* | <0.0001 | 0.499 | <0.0001 |
| | 7 | *SF3B1,ATRX,EGFR,BAP1,EIF1AX,RAB35,MET* | <0.0001 | 0.998 | <0.0001 |
| | 8 | *SF3B1,ATRX,EGFR,BAP1,EIF1AX,NF1,PIK3R1,POLE* | <0.0001 | 1 | <0.0001 |
| | 9 | *SF3B1,ATRX,EGFR,BAP1,EIF1AX,NF1,PIK3R1,MST1R,RAB35* | <0.0001 | 1 | <0.0001 |
| | 10 | *SF3B1,ATRX,EGFR,BAP1,EIF1AX,NF1,PIK3R1,MST1R,RAB35,RARA* | <0.0001 | 1 | <0.0001 |
| **Melanoma_Lymph Node / Melanoma_P** | 6 | *FLT3,PIK3CD,PTEN,TERT,TET2,ERBB3* | 0.024 | 1 | <0.0001 |
| | 7 | *FLT3,PIK3CD,PTEN,TERT,TET2,ERBB3,MUTYH* | 0.012 | 1 | <0.0001 |
| | 8 | *FLT3,PIK3CD,PTEN,TERT,TET2,ERBB3,MUTYH,AKT3* | 0.002 | 1 | <0.0001 |
| | 9 | *FLT3,PIK3CD,PTEN,TERT,TET2,MUTYH,AKT3,DNMT3B,FLT1* | 0.005 | 1 | <0.0001 |
| | 10 | *FLT3,PIK3CD,PTEN,TERT,TET2,MUTYH,AKT3,DNMT3B,FLT1,NF2* | 0.003 | 1 | <0.0001 |

#### *Other metastatic cancers from Melanoma*

Table S 2.3.5. Melanoma_Brain and Melanoma_Liver specific mutated driver gene sets relative to each other

| Type | k | Specific gene set | p1 | p2 | q |
|---|---|---|---|---|---|
| Melanoma_Brain / Melanoma_Liver | 6 | *ANKRD11,DNMT1,EPHB1,PGR,PTEN,TSC1* | 0.002 | 1 | <0.0001 |
| | 7 | *ANKRD11,DNMT1,EPHB1,PGR,PTEN,TSC1,DICER1* | <0.0001 | 1 | <0.0001 |
| | 8 | *ANKRD11,DNMT1,EPHB1,PGR,PTEN,TSC1,PIK3R2,PRKAR1A* | <0.0001 | 1 | <0.0001 |
| | 9 | *ANKRD11,DNMT1,EPHB1,PGR,PTEN,TSC1,PIK3R2,PRKAR1A,DICER1* | <0.0001 | 1 | <0.0001 |
| | 10 | *ANKRD11,CSF1R,EPHB1,PGR,PTEN,ERRFI1,PIK3C3,ETV1,IRS2,DICER1* | <0.0001 | 1 | <0.0001 |
| Melanoma_Liver / Melanoma_Brain | 2 | *BAP1,SF3B1* | 0.027 | 0.933 | 0.036 |
| | 3 | *BRCA2,GNA11,GNAQ* | 0.001 | 1 | 0.004 |
| | 4 | *BAP1,CTNNB1,GLI1,SF3B1* | <0.0001 | 0.975 | <0.0001 |
| | 5 | *BAP1,CTNNB1,GLI1,SF3B1,RARA* | <0.0001 | 0.997 | <0.0001 |
| | 6 | *ARID1B,CTNNB1,GNA11,GNAQ,RARA,STAG2* | <0.0001 | 1 | <0.0001 |
| | 7 | *ARID1B,CTNNB1,GNA11,GNAQ,RARA,STAG2,RB1* | <0.0001 | 1 | <0.0001 |
| | 8 | *BAP1,CTNNB1,EIF1AX,GLI1,RARA,SF3B1,CASP8,PIK3CA* | <0.0001 | 1 | <0.0001 |
| | 9 | *BAP1,CTNNB1,EIF1AX,GLI1,RARA,SF3B1,CASP8,PIK3CA,ABL1* | <0.0001 | 1 | <0.0001 |
| | 10 | *BAP1,CTNNB1,EIF1AX,GLI1,RARA,SF3B1,CASP8,PIK3CA,MDM2,ST6GALNAC3* | <0.0001 | 0.999 | <0.0001 |

Table S 2.3.6. Melanoma_Brain and Melanoma_Lung specific mutated driver gene sets relative to each other
N

| Type | k | Specific gene set | p1 | p2 | q |
|---|---|---|---|---|---|
| **Melanoma_Brain / Melanoma_Lung** | 2 | *KMT2D,PTEN* | 0.024 | 0.963 | 0.02 |
| | 3 | *KMT2D,PTEN,CDK12* | 0.016 | 0.995 | 0.005 |
| | 5 | *KMT2D,PTEN,CDK12,NOTCH2,AXIN2* | 0.009 | 1 | <0.0001 |
| | 6 | *KMT2D,PTEN,CDK12,NOTCH2,AXIN2,PRKAR1A* | 0.002 | 1 | <0.0001 |
| | 7 | *KMT2D,PTEN,CDK12,NOTCH2,AXIN2,PRKAR1A,TNFRSF14* | <0.0001 | 1 | <0.0001 |
| | 8 | *KMT2D,PTEN,CDK12,NOTCH2,AXIN2,PRKAR1A,TNFRSF14,SOX17* | <0.0001 | 1 | <0.0001 |
| | 9 | *BIRC3,EZH2,HGF,MAP3K1,MDM4,AXIN2,RAD51C,TNFRSF14,TSC2* | <0.0001 | 1 | <0.0001 |
| | 10 | *KMT2D,ATR,CDK12,NOTCH2,AXIN2,PRKAR1A,TNFRSF14,MAP3K1,PAX5,RAD51C* | 0.003 | 1 | <0.0001 |
| **Melanoma_Lung / Melanoma_Brain** | 2 | *TP53,NRAS* | 0.027 | 0.814 | 0.048 |
| | 3 | *TP53,NRAS,ASXL2* | 0.003 | 0.862 | 0.009 |
| | 7 | *ABL1,ARID1B,CIC,GATA1,GNA11,POLE,SH2D1A* | 0.017 | 1 | <0.0001 |
| | 8 | *ABL1,ARID1B,CIC,GATA1,CHEK1,POLE,SH2D1A,TENT5C* | 0.007 | 1 | <0.0001 |
| | 9 | *ABL1,ARID1B,CIC,GATA1,GNA11,POLE,SH2D1A,TENT5C,AURKB* | <0.0001 | 1 | <0.0001 |
| | 10 | *ABL1,ARID1B,CIC,GATA1,CHEK1,POLE,SH2D1A,TENT5C,TGFBR1,DAXX* | 0.002 | 1 | <0.0001 |

Table S 2.3.7. Melanoma_Brain and Melanoma_Lymph Node specific mutated driver gene sets relative to each other
N

| Type | k | Specific gene set | p1 | p2 | q |
|---|---|---|---|---|---|
| **Melanoma_Brain / Melanoma_Lymph Node** | 4 | *MRE11,PALB2,JAK1,PAK1* | 0.026 | 1 | 0.047 |
| | 5 | *MRE11,PALB2,HRAS,MPL,XRCC2* | 0.026 | 1 | 0.036 |
| | 6 | *MAP2K2,MDC1,HRAS,PRKAR1A,TNFRSF14,TSC2* | 0.043 | 1 | 0.003 |
| | 7 | *MRE11,PALB2,HRAS,PRKAR1A,TNFRSF14,XRCC2,MPL* | 0.005 | 1 | 0.011 |
| | 8 | *MRE11,PALB2,AURKA,PRKAR1A,TNFRSF14,XRCC2,MPL,HIST1H3H* | 0.009 | 1 | 0.008 |
| | 10 | *MRE11,CBL,HRAS,CHEK2,IRS2,LATS2,TNFRSF14,MPL,RICTOR,MAP2K2* | 0.002 | 1 | <0.0001 |
| **Melanoma_Lymph Node / Melanoma_Brain** | 2 | *BRAF,NRAS* | <0.0001 | 0.076 | 0.001 |
| | 4 | *MSH6,PTEN,TERT,DNMT3B* | 0.016 | 1 | 0.015 |
| | 5 | *MSH6,PTEN,TERT,ERBB3,TET2* | 0.039 | 1 | 0.028 |
| | 6 | *CTNNB1,ERBB4,MAP2K1,PIK3C2G,PTEN,EPHB1* | 0.001 | 1 | <0.0001 |
| | 7 | *CTNNB1,ERBB4,MAP2K1,PIK3C2G,PTEN,PIK3CG,FLT1* | <0.0001 | 0.999 | <0.0001 |
| | 8 | *CTNNB1,IGF1R,MAP2K1,PIK3C2G,PTEN,PIK3CG,ATR,CDKN2A* | 0.001 | 0.998 | <0.0001 |
| | 9 | *CTNNB1,IGF1R,MAP2K1,PIK3C2G,PTEN,PIK3CG,ATR,CDKN2A,INSR* | <0.0001 | 0.999 | <0.0001 |
| | 10 | *CTNNB1,ERBB4,MAP2K1,PIK3C2G,PTEN,PIK3CG,ATR,ATM,ERCC5,TET2* | <0.0001 | 1 | <0.0001 |

Table S 2.3.8. Melanoma_Liver and Melanoma_Lung specific mutated driver gene sets relative to each other
N

| Type | k | Specific gene set | p1 | p2 | q |
|---|---|---|---|---|---|
| Melanoma_Liver / Melanoma_Lung | 3 | *GNA11,GNAQ,NSD1* | 0.001 | 1 | 0.001 |
| | 4 | *GNA11,GNAQ,NSD1,DNMT3A* | <0.0001 | 1 | <0.0001 |
| | 5 | *BAP1,CTNNB1,GLI1,RARA,SF3B1* | <0.0001 | 1 | <0.0001 |
| | 6 | *BAP1,CTNNB1,GLI1,RARA,SF3B1,PDGFRA* | <0.0001 | 1 | <0.0001 |
| | 7 | *BAP1,CTNNB1,GLI1,RARA,SF3B1,PDGFRA,HRAS* | <0.0001 | 1 | <0.0001 |
| | 8 | *BAP1,CTNNB1,GLI1,RARA,SF3B1,PDGFRA,HRAS,ST6GALNAC3* | <0.0001 | 1 | <0.0001 |
| | 9 | *BAP1,CTNNB1,GLI1,RARA,SF3B1,PDGFRA,HRAS,ST6GALNAC3,SLC12A6* | <0.0001 | 1 | <0.0001 |
| | 10 | *BAP1,CTNNB1,DNMT3A,SF3B1,EIF4A2,HRAS,ST6GALNAC3,SLC12A6,AURKA,NSD1* | <0.0001 | 1 | <0.0001 |
| Melanoma_Lung / Melanoma_Liver | 4 | *NRAS,BCOR,CIC,IRS2* | 0.006 | 0.994 | 0.001 |
| | 6 | *NRAS,AR,MED12,PTEN,RAC1,ANKRD11* | <0.0001 | 1 | <0.0001 |
| | 7 | *NRAS,AR,MED12,PTEN,RAC1,IKZF1,IRS2* | 0.001 | 1 | <0.0001 |
| | 8 | *FLT4,AR,MED12,PTEN,MALT1,PGR,IRS2,PMS1* | 0.01 | 1 | <0.0001 |
| | 9 | *ATM,AR,MED12,PTEN,MALT1,FOXP1,IRS2,PMS1,IKZF1* | 0.012 | 1 | <0.0001 |
| | 10 | *ATM,AR,MED12,PTEN,MALT1,FOXP1,IRS2,PMS1,IKZF1,PIM1* | 0.002 | 1 | <0.0001 |

Table S 2.3.9. Melanoma_Liver and Melanoma_Lymph Node specific mutated driver gene sets relative to each other N

| Type | k | Specific gene set | p1 | p2 | q |
|---|---|---|---|---|---|
| Melanoma_Liver / Melanoma_Lymph Node | 2 | *GNA11,GNAQ* | 0.043 | 0.982 | 0.043 |
| | 3 | *GNA11,GNAQ,EGFR* | 0.005 | 0.999 | <0.0001 |
| | 4 | *BAP1,CBL,MGA,SF3B1* | <0.0001 | 1 | <0.0001 |
| | 5 | *BAP1,DNMT3A,SF3B1,RARA,RNF43* | <0.0001 | 1 | <0.0001 |
| | 6 | *BAP1,DNMT3A,SF3B1,RARA,RNF43,LATS2* | <0.0001 | 1 | <0.0001 |
| | 7 | *BAP1,DNMT3A,SF3B1,TSC2,RNF43,LATS2,ST6GALNAC3* | <0.0001 | 1 | <0.0001 |
| | 8 | *BAP1,DNMT3A,SF3B1,RARA,RNF43,LATS2,ST6GALNAC3,SUB1* | <0.0001 | 1 | <0.0001 |
| | 9 | *BAP1,DNMT3A,SF3B1,TSC2,RNF43,LATS2,ST6GALNAC3,SUB1,CD79A* | <0.0001 | 1 | <0.0001 |
| | 10 | *BAP1,DNMT3A,SF3B1,RARA,RNF43,LATS2,ST6GALNAC3,SUB1,CD79A,SLC12A6* | <0.0001 | 1 | <0.0001 |
| Melanoma_Lymph Node / Melanoma_Liver | 2 | *BRAF,NRAS* | <0.0001 | 0.661 | <0.0001 |
| | 3 | *BRAF,NRAS,PDGFRA* | <0.0001 | 0.911 | <0.0001 |
| | 4 | *MSH6,PTEN,TERT,DNMT3B* | 0.038 | 0.994 | 0.005 |
| | 5 | *MSH6,PTEN,TERT,DNMT3B,NF2* | 0.017 | 1 | 0.001 |
| | 6 | *MSH6,PTEN,TERT,ATR,TET2,TRAF2* | 0.018 | 0.999 | <0.0001 |
| | 7 | *MSH6,PTEN,TERT,ATR,TET2,AKT3,PIK3CD* | 0.001 | 1 | <0.0001 |
| | 8 | *MSH6,PTEN,TERT,ATR,TET2,AKT3,PIK3CD,TRAF2* | <0.0001 | 1 | <0.0001 |
| | 9 | *MSH6,PTEN,TERT,NF2,TET2,AKT3,PIK3CD,TRAF2,ERBB3* | 0.002 | 1 | <0.0001 |
| | 10 | *MSH6,PTEN,TERT,NF2,TET2,AKT3,PIK3CD,TRAF2,ATR,ZFHX3* | <0.0001 | 1 | <0.0001 |

Table S 2.3.10. Melanoma_Lung and Melanoma_Lymph Node specific mutated driver gene sets relative to each other N

| Type | k | Specific gene set | p1 | p2 | q |
|---|---|---|---|---|---|
| **Melanoma_Lung / Melanoma_Lymph Node** | 6 | *ABL1,GATA1,PMS1,POLE,TCF3,SH2D1A* | 0.006 | 0.998 | <0.0001 |
| | 7 | *ABL1,GATA1,PMS1,POLE,TCF3,TRAF7,CHEK1* | <0.0001 | 1 | <0.0001 |
| | 8 | *ABL1,GATA1,PMS1,POLE,TCF3,TRAF7,CHEK1,AURKB* | <0.0001 | 0.999 | <0.0001 |
| | 9 | *ABL1,GATA1,PMS1,POLE,TCF3,TRAF7,CHEK1,MALT1,FH* | <0.0001 | 0.997 | <0.0001 |
| | 10 | *ABL1,GATA1,PMS1,POLE,CHEK1,PAK1,CHEK2,MALT1,SH2D1A,IGF2* | <0.0001 | 1 | <0.0001 |
| **Melanoma_Lymph Node / Melanoma_Lung** | 2 | *BRAF,NF1* | <0.0001 | 0.985 | <0.0001 |
| | 3 | *ATM,BRAF,NRAS* | <0.0001 | 0.796 | <0.0001 |
| | 6 | *CTNNB1,KMT2D,NOTCH2,PIK3C2G,PTEN,TET1* | <0.0001 | 1 | <0.0001 |
| | 7 | *CTNNB1,KMT2D,NOTCH2,PIK3C2G,PTEN,TET1,ATR* | <0.0001 | 1 | <0.0001 |
| | 8 | *CTNNB1,KMT2D,NOTCH2,PIK3C2G,PTEN,TET1,ERBB4,CDKN2A* | 0.01 | 1 | <0.0001 |
| | 9 | *CTNNB1,KMT2D,NOTCH2,PIK3C2G,PTEN,TET1,ERBB4,CDKN2A,ATR* | 0.014 | 1 | <0.0001 |
| | 10 | *CTNNB1,MAP2K1,NF2,INSR,PIK3C2G,PTEN,TET1,PIK3CG,CDKN2A,ATR* | <0.0001 | 1 | <0.0001 |

### 3 Supplementary Tables: Three main metastatic lesions attract cancer cells to adapt to their tissue microenvironment

#### 3.1 Hepatocellular carcinoma

Table S 3.1.1. Significant common driver gene set between Breast, Lung, Melanoma, Colorectal and Pancreatic_Liver

| k | common gene set | p1 | p2 | p3 | p4 | p5 | q |
|---|---|---|---|---|---|---|---|
| 7 | *TP53,CDH1,EIF1AX,GATA3,GNA11,GNAQ,MEN1* | <0.0001 | 0.026 | 0.014 | 0.011 | <0.0001 | <0.0001 |
| 8 | *TP53,CDH1,EIF1AX,GATA3,GNA11,GNAQ,MEN1,NFE2L2* | <0.0001 | 0.045 | 0.0132 | 0.024 | <0.0001 | <0.0001 |
| 9 | *TP53,CDH1,EIF1AX,GATA3,GNA11,GNAQ,MEN1,EIF4A2,LDLR* | <0.0001 | 0.009 | 0.003 | 0.015 | <0.0001 | <0.0001 |
| 10 | *TP53,CDH1,EIF1AX,GATA3,GNA11,GNAQ,MEN1,EIF4A2,LDLR,CHEK2* | <0.0001 | 0.01 | 0.004 | 0.015 | <0.0001 | <0.0001 |

Table S 3.1.2. Significant common driver gene set between Colorectal, Pancreatic and Prostate_Liver

| k | common gene set | p1 | p2 | p3 | q |
|---|---|---|---|---|---|
| 7 | *TP53,CD79B,EIF1AX,HIST1H3C,MEN1,SMAD2,STK11* | <0.0001 | <0.0001 | 0.042 | <0.0001 |
| 8 | *TP53,CD79B,EIF1AX,HIST1H3C,MEN1,SMAD2,STK11,HIST1H3E* | <0.0001 | <0.0001 | 0.048 | <0.0001 |

Table S 3.1.3. Significant common driver gene set between Colarectal_Liver and Prostate_Liver

| k | common gene set | p1 | p2 | q |
|---|---|---|---|---|
| 8 | *TP53,BRCA1,CD79B,DDR2,EIF1AX,FGFR4,MEN1,SMAD2* | <0.0001 | 0.02 | <0.0001 |
| 10 | *TP53,BRCA1,CD79B,DDR2,EIF1AX,FGFR4,MEN1,SMAD2,COX7C,HIST1H3C* | <0.0001 | 0.008 | <0.0001 |

*There is no significant common among driver gene set between Breast, Lung, Melanoma, Colorectal, Pancreatic, Prostate_Liver when k=2 $\sim$ 10.

Table S 3.1.4. Significant common driver gene set between Breast, Lung and Melanoma_Liver

| k | common gene set | p1 | p2 | p3 | q |
|---|---|---|---|---|---|
| 10 | *TP53,ASXL1,CDH1,DICER1,ESR1,GNA11,GNAQ,IGF1,MEN1,MSH6* | <0.0001 | 0.016 | <0.0001 | <0.0001 |

#### 3.2 Non-small-cell Lung cancer

Table S 3.2.1. Significant common driver gene set between Breast, Colorectal, HeadNeck_Lung

| k | common gene set | p1 | p2 | p3 | q |
|---|---|---|---|---|---|
| 4 | *TP53,GATA3,PIK3CA,PIK3R1* | <0.0001 | <0.0001 | 0.029 | <0.0001 |
| 5 | *TP53,GATA3,PIK3CA,PIK3R1,MDC1* | <0.0001 | 0.001 | 0.007 | <0.0001 |
| 6 | *TP53,GATA3,PIK3CA,PIK3R1,MDC1,B2M* | <0.0001 | <0.0001 | 0.003 | <0.0001 |
| 7 | *TP53,GATA3,PIK3CA,PIK3R1,MDC1,B2M,LATS1* | <0.0001 | <0.0001 | <0.0001 | <0.0001 |
| 8 | *TP53,GATA3,PIK3CA,PIK3R1,MDC1,B2M,LATS1,CCND1* | <0.0001 | <0.0001 | <0.0001 | <0.0001 |
| 9 | *TP53,GATA3,PIK3CA,PIK3R1,MDC1,B2M,LATS1,CCND1,HIST1H3F* | <0.0001 | <0.0001 | <0.0001 | <0.0001 |
| 10 | *TP53,GATA3,PIK3CA,PIK3R1,MDC1,B2M,LATS1,CCND1,HIST1H3F,GREM1* | <0.0001 | <0.0001 | <0.0001 | <0.0001 |

*There is no significant common among driver gene set between Breast, Colorectal, HeadNeck, Melanoma_Lung when k=2 $\sim$ 10.

Table S 3.2.2. Significant common driver gene set between Colorectal, HeadNeck and Melanoma_Lung

| k | common gene set | p1 | p2 | p3 | q |
|---|---|---|---|---|---|
| 7 | *TP53,AURKB,B2M,NRAS,PIK3CA,PIK3R1,CCND3* | <0.0001 | 0.025 | 0.04 | <0.0001 |
| 8 | *TP53,AURKB,B2M,NRAS,PIK3CA,PIK3R1,BMPR1A,WT1* | <0.0001 | 0.017 | 0.026 | <0.0001 |
| 9 | *TP53,AURKB,B2M,NRAS,PIK3CA,PIK3R1,BMPR1A,RAB35,HIST1H3F* | <0.0001 | 0.008 | 0.007 | <0.0001 |
| 10 | *TP53,AURKB,B2M,NRAS,PIK3CA,PIK3R1,BMPR1A,PIM1,SMAD2,FH* | <0.0001 | 0.016 | 0.002 | <0.0001 |

Table S 3.2.3. Significant common driver gene set between Breast_Lung and Melanoma_Lung

| k | common gene set | p1 | p2 | q |
|---|---|---|---|---|
| **5** | *TP53,ASXL2,AURKB,GATA3,NRAS* | <0.0001 | 0.010 | <0.0001 |
| **6** | *TP53,ASXL2,AURKB,CCND1,GATA3,NRAS* | <0.0001 | 0.011 | <0.0001 |
| **7** | *TP53,ASXL2,AURKB,CCND1,GATA3,NRAS,MAP2K4* | <0.0001 | 0.01 | <0.0001 |
| **8** | *TP53,ASXL2,AURKB,CCND1,GATA3,NRAS,MAP2K4,FH* | <0.0001 | 0.006 | <0.0001 |
| **9** | *TP53,ASXL2,AURKB,CCND1,GATA3,NRAS,MAP2K4,GNA11,SMAD2* | <0.0001 | 0.004 | <0.0001 |
| **10** | *TP53,ASXL2,AURKB,CCND1,GATA3,NRAS,MAP2K4,GNA11,FH,INPP4B* | <0.0001 | <0.0001 | <0.0001 |

#### 3.3 Brain cancer

Table S 3.3.1. Significant common driver gene set between Breast_Brain and Lung_Brain

| k | common gene set | p1 | p2 | q |
|---|---|---|---|---|
| 4 | *TP53,FGFR1,HIST1H3E,RAF1* | 0.022 | <0.0001 | <0.0001 |
| 5 | *TP53,FGFR1,MAP2K4,RAF1,KIF5B* | <0.0001 | <0.0001 | <0.0001 |
| 6 | *TP53,FGFR1,HIST1H3E,RAF1,KIF5B,MAP2K4* | <0.0001 | <0.0001 | <0.0001 |
| 7 | *TP53,FGFR1,HIST1H3E,RAF1,KIF5B,MAP2K4,EIF1AX* | <0.0001 | <0.0001 | <0.0001 |
| 8 | *TP53,FGFR1,HIST1H3E,RAF1,KIF5B,MAP2K4,EIF1AX,PRDM1* | <0.0001 | <0.0001 | <0.0001 |
| 9 | *TP53,FGFR1,HIST1H3E,RAF1,KIF5B,MAP2K4,EML4,PRDM1,GATA1* | <0.0001 | <0.0001 | <0.0001 |
| 10 | *TP53,FGFR1,HIST1H3E,RAF1,KIF5B,MAP2K4,EML4,PRDM1,GATA1,SDHD* | <0.0001 | <0.0001 | <0.0001 |

Table S 3.3.2. Significant common driver gene set between Breast_Brain and Melanoma_Brain

| k | common gene set | p1 | p2 | q |
|---|---|---|---|---|
| 6 | *TP53,BRAF,FLCN,GATA3,HRAS,NRAS* | 0.033 | <0.0001 | <0.0001 |
| 7 | *TP53,BRAF,FLCN,GSK3B,HRAS,NRAS,CDK4* | <0.0001 | <0.0001 | <0.0001 |
| 8 | *TP53,BRAF,FLCN,GATA3,HRAS,NRAS,CDK4,AKT3* | <0.0001 | <0.0001 | <0.0001 |
| 9 | *TP53,BRAF,FLCN,GATA3,HRAS,NRAS,CDK4,AKT3,EIF4A2* | <0.0001 | <0.0001 | <0.0001 |
| 10 | *TP53,BRAF,FLCN,GATA3,HRAS,NRAS,KIT,AKT3,EIF4A2,ABRAXAS1* | <0.0001 | <0.0001 | <0.0001 |

*There is no significant common driver gene set between Lung_Brain and Melanoma_Brain when k=2 $\sim$ 10.
*There is no significant common driver gene set between Breast, Lung and Melanoma_Brain when k=2 $\sim$ 10.

### 4 Supplementary Tables: Typical patterns with high metastatic tropisms

#### 4.1 colorectal liver metastases

Table S 4.1.1. Significant common driver gene set between Colorectal_Liver and Colorectal_P

| k | common gene set | p1 | p2 | q |
|---|---|---|---|---|
| 9 | *APC,BMPR1A,EIF4A2,GATA2,H3F3A,PDPK1,RAB35,RHOA,STK40* | <0.0001 | 0.028 | <0.0001 |

Table S 4.1.2. Colorectal_P specific mutated driver gene sets relative to Colorectal_Liver

| k | Specific gene set | p1 | p2 | q |
|---|---|---|---|---|
| 2 | *BRAF,APC* | <0.0001 | 0.263 | <0.0001 |
| 9 | *BRAF,CSF1R,ERBB2,FLT3,FLT4,KRAS,PTEN,RBM10,CDH1* | 0.022 | 0.959 | 0.009 |
| 10 | *BRAF,CSF1R,ERBB2,FLT3,FLT4,KRAS,NSD1,RBM10,PIK3R1,PRKN* | 0.006 | 0.851 | 0.003 |

*Colorectal_Liver has no specific mutated driver gene sets relative to Colorectal_P when k=2 $\sim$ 10.

Table S 4.1.3. Significant common driver gene set between Colorectal_liver and Hepatocellular_P

| k | common gene set | p1 | p2 | q |
|---|---|---|---|---|
| 10 | *APC,AXIN1,BCOR,CDH1,CDK4,CTNNB1,EZH2,INPP4B,MAP2K1,NPM1* | <0.0001 | 0.042 | <0.0001 |

Table S 4.1.4. Colorectal_Liver and Hepatocellular_P specific mutated driver gene sets relative to each other

| Type | k | Specific gene set | p1 | p2 | q |
|---|---|---|---|---|---|
| **Colorectal_Liver / Hepatocellular_P** | 3 | *APC,EGFR,PHOX2B* | 0.007 | 1 | <0.0001 |
| | 4 | *APC,EGFR,PHOX2B,NOTCH3* | <0.0001 | 1 | <0.0001 |
| | 5 | *APC,EGFR,PHOX2B,JUN,NOTCH3* | 0.002 | 1 | <0.0001 |
| | 6 | *APC,EGFR,PHOX2B,JUN,NOTCH3,ESR1* | <0.0001 | 1 | <0.0001 |
| | 7 | *APC,EGFR,PHOX2B,AXL,KRAS,MET,BCOR* | 0.003 | 0.943 | 0.001 |
| | 8 | *APC,EGFR,PHOX2B,AXL,KRAS,MET,LMO1,NPM1* | <0.0001 | 0.941 | 0.001 |
| | 9 | *APC,EGFR,PHOX2B,AXL,KRAS,MET,LMO1,NPM1,BCOR* | <0.0001 | 0.934 | <0.0001 |
| | 10 | *TP53,ATRX,BRAF,ERBB2,FGFR3,KRAS,MST1R,NRAS,RAD51B,STK11* | <0.0001 | 1 | <0.0001 |
| **Hepatocellular_P / Colorectal_Liver** | 2 | *JAK1,TERT* | 0.034 | 1 | <0.0001 |
| | 3 | *JAK1,TERT,BAP1* | 0.007 | 1 | <0.0001 |
| | 4 | *JAK1,TERT,BAP1,KLF4* | <0.0001 | 1 | <0.0001 |
| | 5 | *JAK1,TERT,BAP1,KLF4,XPO1* | <0.0001 | 1 | <0.0001 |
| | 6 | *JAK1,TERT,BAP1,KLF4,XPO1,TGFBR1* | <0.0001 | 1 | <0.0001 |
| | 7 | *JAK1,TERT,BAP1,KLF4,XPO1,TGFBR1,PMAIP1* | <0.0001 | 1 | <0.0001 |
| | 8 | *JAK1,TERT,BAP1,KLF4,XPO1,TGFBR1,EZH2,NEGR1* | <0.0001 | 1 | <0.0001 |
| | 9 | *JAK1,TERT,BAP1,KLF4,XPO1,TGFBR1,PMAIP1,NEGR1,PRKACA* | <0.0001 | 1 | <0.0001 |
| | 10 | *JAK1,TERT,BAP1,KLF4,XPO1,TGFBR1,PMAIP1,NEGR1,PRKACA,SDHB* | <0.0001 | 1 | <0.0001 |

#### 4.2 NSCLC cancer with brain metastasis

Table S 4.2.1. Significant common driver gene set between Lung_Brain and Lung_P

| k | common gene set | p1 | p2 | q |
|---|---|---|---|---|
| 8 | *BRAF,EGFR,EML4,ERBB2,KIF5B,KRAS,MET,ROS1* | <0.0001 | <0.0001 | <0.0001 |
| 10 | *BRAF,EGFR,EML4,ERBB2,KIF5B,KRAS,MET,ROS1,RASA1,GATA1* | <0.0001 | <0.0001 | <0.0001 |

Lung_P and Lung_Brain specific mutated driver gene sets relative to each other can be seen in S2.2.2.

Table S 4.2.2. Lung_Brain and Brain_P specific mutated driver gene sets relative to each other

| **Type** | **k** | **Specific gene set** | **p1** | **p2** | **q** |
|---|---|---|---|---|---|
| **Lung_Brain / Brain_P** | 4 | *AXIN1,IL7R,KEAP1,NFE2L2* | 0.43 | 1 | <0.0001 |
| | 5 | *AXIN1,IL7R,KEAP1,KIF5B,NFE2L2* | 0.012 | 1 | <0.0001 |
| | 7 | *AXIN1,IL7R,KEAP1,KIF5B,NFE2L2,STK11,EPHA7* | 0.028 | 1 | <0.0001 |
| | 8 | *AXIN1,IL7R,KEAP1,KIF5B,NFE2L2,STK11,EPHA7,AKT3* | 0.024 | 1 | <0.0001 |
| | 9 | *AXIN1,IL7R,KEAP1,KIF5B,NFE2L2,STK11,EPHA7,AKT3,B2M* | 0.039 | 1 | <0.0001 |
| | 10 | *AXIN1,IL7R,KEAP1,KIF5B,NFE2L2,STK11,EPHA7,AKT3,B2M,CTNNB1* | 0.006 | 1 | <0.0001 |
| **Brain_P / Lung_Brain** | 2 | *ATRX,TERT* | <0.0001 | 0.972 | <0.0001 |
| | 3 | *ATRX,TERT,BRAF* | <0.0001 | 1 | <0.0001 |
| | 4 | *ATRX,TERT,BRAF,H3F3A* | <0.0001 | 0.999 | <0.0001 |
| | 5 | *ATRX,TERT,BRAF,H3F3A,MAX* | <0.0001 | 0.999 | <0.0001 |
| | 6 | *ATRX,TERT,BRAF,CDKN2B,H3F3A,MAX* | <0.0001 | 1 | <0.0001 |
| | 7 | *ATRX,TERT,BRAF,FANCA,H3F3A,MAX,H3F3C* | <0.0001 | 1 | <0.0001 |
| | 8 | *ATRX,TERT,BRAF,FANCA,H3F3A,MAX,CDKN2B,ZNF710* | <0.0001 | 0.999 | <0.0001 |
| | 9 | *ATRX,TERT,BRAF,FANCA,H3F3A,MAX,GOPC,H3F3C,RAC1* | <0.0001 | 0.999 | <0.0001 |
| | 10 | *ATRX,TERT,BRAF,AHCYL2,H3F3A,CDKN2B,GOPC,HDAC5,IDH1,TRAP1* | <0.0001 | 0.998 | <0.0001 |

#### 4.3 breast and prostate cancer with bone metastasis

Table S 4.3.1. Significant common driver gene set between Breast_P and Breast_Bone

| k | common gene set | p1 | p2 | q |
|---|---|---|---|---|
| 2 | *TP53,GATA3* | <0.0001 | 0.018 | <0.0001 |
| 3 | *TP53,GATA3,CDH1* | <0.0001 | <0.0001 | <0.0001 |
| 4 | *TP53,GATA3,CDH1,SMARCA4* | <0.0001 | <0.0001 | <0.0001 |
| 5 | *TP53,GATA3,CDH1,SMARCA4,AMER1* | <0.0001 | <0.0001 | <0.0001 |
| 6 | *TP53,CDH1,DNMT3B,GATA3,PAK5,SMARCA4* | <0.0001 | <0.0001 | <0.0001 |
| 7 | *TP53,AMER1,CDH1,GATA3,RHEB,SMARCA4,TNFAIP3* | <0.0001 | <0.0001 | <0.0001 |
| 8 | *TP53,CDH1,DNMT3B,GATA3,PAK5,RHEB,SMARCA4,TNFAIP3* | <0.0001 | <0.0001 | <0.0001 |
| 9 | *TP53,CDH1,DNMT3B,GATA3,PAK5,RHEB,SMARCA4,TNFAIP3,SPOP* | <0.0001 | <0.0001 | <0.0001 |
| 10 | *TP53,CDH1,DNMT3B,GATA3,PAK5,RHEB,SMARCA4,TNFAIP3,SPOP,FH* | <0.0001 | <0.0001 | <0.0001 |

Breast_P and Breast_Bone specific mutated driver gene sets relative to each other can be seen in S2.1.1.

Table S 4.3.2. Significant common driver gene set between Breast_Bone and Prostate_Bone

| k | common gene set | p1 | p2 | q |
|---|---|---|---|---|
| 6 | *TP53,APC,ARID1A,GATA3,KMT2D,SMARCA4* | <0.0001 | 0.034 | <0.0001 |
| 7 | *TP53,APC,ARID1A,GATA3,KMT2D,SMARCA4,MITF* | <0.0001 | 0.011 | <0.0001 |
| 8 | *TP53,APC,ARID1A,GATA3,KMT2D,SMARCA4,MITF,TBX3* | <0.0001 | 0.016 | <0.0001 |
| 9 | *TP53,APC,ARID1A,GATA3,KMT2D,SMARCA4,MITF,TBX3,NTRK3* | <0.0001 | 0.007 | <0.0001 |
| 10 | *TP53,APC,ARID1A,GATA3,KMT2D,SMARCA4,MITF,TBX3,NTRK3,VHL* | <0.0001 | 0.003 | <0.0001 |

*Prostate_P has no specific mutated driver gene sets relative to Prostate_Bone when k=2 $\sim$ 10.
*Prostate_Bone has no specific mutated driver gene sets relative to Prostate_P when k=2 $\sim$ 10.

Table S 4.3.3. Breast_Bone and Prostate_Bone specific mutated driver gene sets relative to each other

| Type | k | Specific gene set | p1 | p2 | q |
|---|---|---|---|---|---|
| **Breast_Bone / Prostate_Bone** | 7 | *AKT1,BRIP1,CDH1,IKBKE,PIK3CA,SOX17,TBX3* | 0.033 | 1 | 0.047 |
| **Prostate_Bone / Breast_Bone** | 5 | *CDK12,EGFR,KDM6A,TMPRSS2,AR* | 0.035 | 1 | 0.035 |
| | 6 | *CDK12,EGFR,KDM6A,TMPRSS2,AR,VHL* | 0.011 | 1 | 0.018 |
| | 7 | *CDK12,EGFR,KDM6A,TMPRSS2,AR,NOTCH4,RNF43* | 0.003 | 0.999 | <0.0001 |
| | 8 | *CDK12,EGFR,KDM6A,TMPRSS2,AR,NOTCH4,RNF43,CTNNB1* | <0.0001 | 0.998 | <0.0001 |
| | 9 | *CDK12,EGFR,KDM6A,TMPRSS2,AR,NOTCH4,RNF43,CTNNB1,JAK1* | <0.0001 | 0.999 | <0.0001 |
| | 10 | *CDK12,EGFR,KDM6A,TMPRSS2,NOTCH4,RNF43,CTNNB1,JAK1,PBRM1,SPOP* | <0.0001 | 1 | <0.0001 |

Table S 4.3.4. Significant common driver gene set between Prostate_P and Prostate_Bone

| k | common gene set | p1 | p2 | q |
|---|---|---|---|---|
| 3 | *FOXA1,SPOP,TMPRSS2* | 0.001 | <0.0001 | <0.0001 |
| 4 | *FOXA1,SPOP,TMPRSS2,KDM6A* | 0.002 | <0.0001 | <0.0001 |
| 5 | *FOXA1,SPOP,TMPRSS2,KMT2A,PTPRD* | <0.0001 | <0.0001 | <0.0001 |
| 6 | *FOXA1,SPOP,TMPRSS2,KMT2A,PTPRD,ASXL2* | <0.0001 | <0.0001 | <0.0001 |
| 7 | *FOXA1,SPOP,TMPRSS2,KDM6A,PTPRD,MTOR,POLE* | 0.001 | <0.0001 | <0.0001 |
| 8 | *FOXA1,SPOP,TMPRSS2,KDM6A,PTPRD,MTOR,NOTCH4,ASXL2* | 0.001 | <0.0001 | <0.0001 |
| 9 | *FOXA1,SPOP,TMPRSS2,KDM6A,PTPRD,MTOR,NOTCH4,POLE,HRAS* | 0.024 | <0.0001 | <0.0001 |
| 10 | *FOXA1,SPOP,TMPRSS2,KMT2A,PTPRD,ARID1B,NOTCH4,ASXL2,FLT1,IKBKE* | 0.002 | <0.0001 | <0.0001 |

### 5 Supplementary Tables: Metastasis-promoting mechanisms for comparable cancers

#### 5.1 Female-Breast, Endometrium and Ovary

*Primary cancers*

Table S 5.1.1. Significant common driver gene set between Breast_P and Endometrial_P

| k | common gene set | p1 | p2 | q |
|---|---|---|---|---|
| 4 | *TP53,CDH1,GATA3,ARID1A* | <0.0001 | <0.0001 | <0.0001 |
| 5 | *TP53,CDH1,GATA3,CD276,PTEN* | <0.0001 | <0.0001 | <0.0001 |
| 6 | *TP53,CDH1,GATA3,CD276,ARID1A,SMARCD1* | <0.0001 | <0.0001 | <0.0001 |
| 7 | *TP53,CDH1,GATA3,CTNNB1,MALT1,NFE2L2,SOX17* | <0.0001 | <0.0001 | <0.0001 |
| 8 | *TP53,CDH1,GATA3,CTNNB1,NFE2L2,SOX17,CD276,YES1* | <0.0001 | <0.0001 | <0.0001 |
| 9 | *TP53,CDH1,GATA3,CTNNB1,NFE2L2,SOX17,CD276,YES1,MALT1* | <0.0001 | <0.0001 | <0.0001 |
| 10 | *TP53,CDH1,GATA3,CTNNB1,NFE2L2,SOX17,CD276,YES1,MALT1,KRAS* | <0.0001 | <0.0001 | <0.0001 |

Table S 5.1.2. Significant common driver gene set between Breast_P and Ovarian_P

| k | common gene set | p1 | p2 | q |
|---|---|---|---|---|
| 4 | *TP53,ARID1A,CDH1,GATA3* | <0.0001 | <0.0001 | <0.0001 |
| 5 | *TP53,ARID1A,CDH1,GATA3,CDC73* | <0.0001 | <0.0001 | <0.0001 |
| 6 | *TP53,ARID1A,CDH1,GATA3,CDC73,DAXX* | <0.0001 | <0.0001 | <0.0001 |
| 7 | *TP53,ARID1A,CDH1,GATA3,KRAS,NUP93,TNFAIP3* | <0.0001 | <0.0001 | <0.0001 |
| 8 | *TP53,ARID1A,CDH1,GATA3,KRAS,DAXX,TNFAIP3,CDC73* | <0.0001 | <0.0001 | <0.0001 |
| 9 | *TP53,ARID1A,CDH1,GATA3,KRAS,DAXX,TNFAIP3,CDC73,EED* | <0.0001 | <0.0001 | <0.0001 |
| 10 | *TP53,ARID1A,CDH1,GATA3,KRAS,DAXX,TNFAIP3,CDC73,EED,SMARCD1* | <0.0001 | <0.0001 | <0.0001 |

Table S 5.1.3. Significant common driver gene set between Endometrial_P and Ovarian_P

| k | common gene set | p1 | p2 | q |
|---|---|---|---|---|
| 2 | *TP53,ARID1A* | <0.0001 | <0.0001 | <0.0001 |
| 3 | *TP53,ARID1A,CD276* | <0.0001 | <0.0001 | <0.0001 |
| 4 | *TP53,ARID1A,CD276,CTLA4* | <0.0001 | <0.0001 | <0.0001 |
| 5 | *TP53,ARID1A,CD276,CTLA4,NFKBIA* | <0.0001 | <0.0001 | <0.0001 |
| 6 | *TP53,ARID1A,CD276,CTLA4,NFKBIA,GOLGB1* | <0.0001 | <0.0001 | <0.0001 |
| 7 | *TP53,ARID1A,CD276,CTLA4,NFKBIA,GOLGB1,TCEANC2* | <0.0001 | <0.0001 | <0.0001 |
| 8 | *TP53,ARID1A,CD276,CTLA4,NFKBIA,GOLGB1,TCEANC2,EIF1AX* | <0.0001 | <0.0001 | <0.0001 |
| 9 | *TP53,ARID1A,CD276,CTLA4,NFKBIA,GOLGB1,TCEANC2,EIF1AX,NUP93* | <0.0001 | <0.0001 | <0.0001 |
| 10 | *TP53,ARID1A,CD276,CTLA4,NFKBIA,GOLGB1,TCEANC2,EIF1AX,NUP93,CD79B* | <0.0001 | <0.0001 | <0.0001 |

Table S 5.1.4. Breast_P and Endometrial_P specific mutated driver gene sets relative to each other
N

| Type | k | Specific gene set | p1 | p2 | q |
|---|---|---|---|---|---|
| **Breast_P / Endometrial_P** | 2 | *TP53,PIK3CA* | 0.033 | 0.962 | 0.012 |
| | 3 | *TP53,PIK3CA,GATA3* | <0.0001 | 0.976 | <0.0001 |
| | 4 | *AKT1,PIK3CA,BRCA2,NF1* | <0.0001 | 1 | <0.0001 |
| | 7 | *AKT1,PIK3CA,ARID1A,BRCA2,MAP3K1,NF1,PTEN* | 0.026 | 1 | <0.0001 |
| | 8 | *AKT1,PIK3CA,ARID1A,BRCA2,MAP3K1,NF1,PTEN,ESR1* | 0.009 | 1 | <0.0001 |
| | 10 | *AKT1,PIK3CA,ARID1A,BRCA2,MAP3K1,NF1,PTEN,ESR1,BCOR,KMT2C* | 0.025 | 1 | <0.0001 |
| **Endometrial_P / Breast_P** | 7 | *ACVR1,FBXW7,NTRK3,PIK3R1,PPP2R1A,NFKBIA,KRAS* | 0.038 | 0.981 | 0.025 |
| | 8 | *ACVR1,FBXW7,NTRK3,PIK3R1,PPP2R1A,NFKBIA,KRAS,EIF3G* | 0.03 | 0.978 | 0.016 |
| | 9 | *ACVR1,FBXW7,NTRK3,PIK3R1,PPP2R1A,NFKBIA,KRAS,HIST1H3G,TACC3* | 0.015 | 0.98 | 0.01 |
| | 10 | *FANCA,FBXW7,NTRK3,PIK3R1,PPP2R1A,NFKBIA,SPOP,HIST1H3G,TMEM64,MITF* | 0.006 | 0.984 | 0.011 |

Table S 5.1.5. Breast_P and Ovarian_P specific mutated driver gene sets relative to each other

| Type | k | Specific gene set | p1 | p2 | q |
|---|---|---|---|---|---|
| **Breast_P / Ovarian_P** | 2 | *TP53,GATA3* | <0.0001 | 1 | <0.0001 |
| | 3 | *TP53,GATA3,CDH1* | <0.0001 | 1 | <0.0001 |
| | 4 | *AKT1,BRCA2,PIK3CA,PTEN* | <0.0001 | 0.989 | <0.0001 |
| | 5 | *AKT1,BRCA2,PIK3CA,PTEN,ERBB2* | <0.0001 | 0.997 | <0.0001 |
| | 6 | *AKT1,BRCA2,PIK3CA,PTEN,MAP3K1,KDM6A* | 0.003 | 0.997 | <0.0001 |
| | 9 | *AKT1,BRCA2,PIK3CA,PTEN,MAP3K1,KMT2C,ESR1,NF1,NOTCH1* | 0.034 | 0.968 | 0.018 |
| | 10 | *AKT1,BRCA2,PIK3CA,PTEN,MAP3K1,KMT2C,ESR1,NF1,NOTCH1,GRIN2A* | 0.024 | 1 | <0.0001 |
| **Ovarian_P / Breast_P** | 10 | *ASXL1,BRAF,CASC3,CTNNB1,FANCA,LATS1,MST1R,NF2,RAD50,SOCS1* | 0.046 | 1 | <0.0001 |

Table S 5.1.6. Endometrial_P and Ovarian_P specific mutated driver gene sets relative to each other

| Type | k | Specific gene set | p1 | p2 | q |
|---|---|---|---|---|---|
| **Endometrial_P / Ovarian_P** | 3 | *PIK3CA,PIK3R1,SPOP* | 0.007 | 0.889 | 0.01 |
| | 4 | *PIK3CA,PIK3R1,SPOP,EP300* | 0.002 | 0.943 | 0.003 |
| | 5 | *PIK3CA,PIK3R1,SPOP,EP300,CBFB* | 0.005 | 0.952 | 0.002 |
| | 10 | *PIK3CA,PPP2R1A,DPP9,SPOP,FBXW7,HIST1H3G,MED12,PTEN,TGFBR1,ATM* | 0.021 | 0.998 | 0.002 |
| **Ovarian_P / Endometrial_P** | 10 | *ANKRD11,ATRX,HGF,KIT,KMT2A,MST1R,NF2,PIK3C2G,ROS1,TERT* | 0.014 | 1 | <0.0001 |

Table S 5.1.7. Significant common driver gene set among Breast_P, Endometrial_P and Ovarian_P

| **k** | **common gene set** | **p1** | **p2** | **p3** | **q** |
|---|---|---|---|---|---|
| 4 | *TP53,ARID1A,CDH1,GATA3* | <0.0001 | <0.0001 | <0.0001 | <0.0001 |
| 5 | *TP53,ARID1A,CDH1,GATA3,CD276* | <0.0001 | <0.0001 | <0.0001 | <0.0001 |
| 6 | *TP53,ARID1A,CDH1,GATA3,CD276,SMARCD1* | <0.0001 | <0.0001 | <0.0001 | <0.0001 |
| 7 | *TP53,ARID1A,CDH1,GATA3,CD276,SMARCD1,SRSF2* | <0.0001 | <0.0001 | <0.0001 | <0.0001 |
| 8 | *TP53,ARID1A,CDH1,GATA3,CD276,SMARCD1,SRSF2,CTLA4* | <0.0001 | <0.0001 | <0.0001 | <0.0001 |
| 9 | *TP53,ARID1A,CDH1,GATA3,CD276,SMARCD1,SRSF2,CTLA4,GOLGB1* | <0.0001 | <0.0001 | <0.0001 | <0.0001 |
| 10 | *TP53,RAID1A,CDH1,GATA3,CD276,SMARCD1,SRSF2,CTLA4,GOLGB1,TCEANC2* | <0.0001 | <0.0001 | <0.0001 | <0.0001 |

*Metastatic cancers*

Table S 5.1.8. Significant common driver gene set between Breast_M and Endometrial_M

| k | common gene set | p1 | p2 | q |
|---|---|---|---|---|
| 2 | *TP53,ARID1A* | <0.0001 | <0.0001 | <0.0001 |
| 3 | *TP53,ARID1A,CD276* | <0.0001 | <0.0001 | <0.0001 |
| 4 | *TP53,ARID1A,CD276,CTLA4* | <0.0001 | <0.0001 | <0.0001 |
| 5 | *TP53,ARID1A,CD276,CTLA4,NFKBIA* | <0.0001 | <0.0001 | <0.0001 |
| 6 | *TP53,ARID1A,CD276,CTLA4,NFKBIA,GOLGB1* | <0.0001 | <0.0001 | <0.0001 |
| 7 | *TP53,ARID1A,CD276,CTLA4,NFKBIA,GOLGB1,TCEANC2* | <0.0001 | <0.0001 | <0.0001 |
| 8 | *TP53,ARID1A,CD276,CTLA4,NFKBIA,GOLGB1,TCEANC2,EIF1AX* | <0.0001 | <0.0001 | <0.0001 |
| 9 | *TP53,ARID1A,CD276,CTLA4,NFKBIA,GOLGB1,TCEANC2,EIF1AX,NUP93* | <0.0001 | <0.0001 | <0.0001 |
| 10 | *TP53,ARID1A,CD276,CTLA4,NFKBIA,GOLGB1,TCEANC2,EIF1AX,NUP93,CD79B* | <0.0001 | <0.0001 | <0.0001 |

Table S 5.1.9. Significant common driver gene set between Breast_M and Ovarian_M

| k | common gene set | p1 | p2 | q |
|---|---|---|---|---|
| 7 | *TP53,CDH1,CDKN1B,GATA3,HGF,KRAS,SMARCA4* | <0.0001 | <0.0001 | <0.0001 |
| 8 | *TP53,CDH1,CDKN1B,GATA3,HGF,KRAS,SMARCA4,CDC73* | <0.0001 | <0.0001 | <0.0001 |
| 9 | *TP53,CDH1,CDKN1B,GATA3,HGF,KRAS,SMARCA4,TSHR,MITF* | <0.0001 | <0.0001 | <0.0001 |
| 10 | *TP53,CDH1,CDKN1B,GATA3,HGF,KRAS,SMARCA4,TSHR,CDC73,POLD1* | <0.0001 | <0.0001 | <0.0001 |

*Ovarian_M has no specific mutated driver gene sets relative to Breast_M when k=2 $\sim$ 10.

Table S 5.1.10. Significant common driver gene set between Endometrial_M and Ovarian_M

| k | common gene set | p1 | p2 | q |
|---|---|---|---|---|
| 2 | *TP53,ARID1A* | <0.0001 | 0.003 | <0.0001 |
| 3 | *TP53,ARID1A,KRAS* | <0.0001 | <0.0001 | <0.0001 |
| 4 | *TP53,CTNNB1,KRAS,HIST1H3F* | <0.0001 | <0.0001 | <0.0001 |
| 5 | *TP53,CTNNB1,KRAS,HIST1H3F,RPS6KB2* | <0.0001 | <0.0001 | <0.0001 |
| 6 | *TP53,CTNNB1,KRAS,HIST1H3F,RPS6KB2,RYBP* | <0.0001 | <0.0001 | <0.0001 |
| 7 | *TP53,CTNNB1,KRAS,HIST1H3F,RPS6KB2,RYBP,EYA2* | <0.0001 | <0.0001 | <0.0001 |
| 8 | *TP53,CTNNB1,KRAS,HIST1H3F,RPS6KB2,RYBP,EYA2,HNRNPUL1* | <0.0001 | <0.0001 | <0.0001 |
| 9 | *TP53,CTNNB1,KRAS,HIST1H3F,RPS6KB2,RYBP,EYA2,HNRNPUL1,TMPRSS2* | <0.0001 | <0.0001 | <0.0001 |
| 10 | *TP53,CTNNB1,KRAS,HIST1H3F,RPS6KB2,RYBP,EYA2,HNRNPUL1,TMPRSS2,FLCN* | <0.0001 | <0.0001 | <0.0001 |

Table S 5.1.11. Breast_M and Endometrial_M specific mutated driver gene sets relative to each othe

| Type | k | Specific gene set | p1 | p2 | q |
|---|---|---|---|---|---|
| **Breast_M / Endometrial_M** | 2 | *TP53,GATA3* | <0.0001 | 0.338 | <0.0001 |
| | 3 | *TP53,GATA3,PIK3CA* | <0.0001 | 0.257 | <0.0001 |
| | 4 | *ERBB2,ESR1,PIK3CA,PTEN* | 0.043 | 0.995 | 0.01 |
| | 5 | *ERBB2,ESR1,PIK3CA,PTEN,AKT1* | 0.001 | 0.982 | <0.0001 |
| | 6 | *ERBB2,ESR1,PIK3CA,PTEN,AKT1,ARID1A* | 0.018 | 1 | <0.0001 |
| | 7 | *ERBB2,ESR1,PIK3CA,PTEN,AKT1,ARID1A,KMT2A* | 0.013 | 1 | <0.0001 |
| **Endometrial_M / Breast_M** | 2 | *KRAS,CTNNB1* | 0.021 | 0.968 | 0.03 |
| | 3 | *KRAS,CTNNB1,NPM1* | 0.012 | 0.965 | 0.006 |
| | 4 | *KRAS,PIK3R1,ACVR1,CD79A* | 0.002 | 0.968 | 0.003 |

Table S 5.1.12. Breast_M specific mutated driver gene sets relative to Ovarian_M

| k | Specific gene set | p1 | p2 | q |
|---|---|---|---|---|
| 2 | *TP53,GATA3* | <0.0001 | 1 | <0.0001 |
| 3 | *TP53,GATA3,PIK3CA* | <0.0001 | 0.087 | <0.0001 |
| 4 | *TP53,GATA3,PIK3CA,ESR1* | <0.0001 | 0.06 | <0.0001 |
| 7 | *AKT1,ARID1A,PIK3CA,ESR1,KMT2A,PTEN,ERBB2* | 0.013 | 1 | 0.001 |
| 8 | *AKT1,ARID1A,PIK3CA,ESR1,KMT2A,PTEN,ERBB2,PIK3R1* | 0.006 | 1 | 0.001 |

Table S 5.1.13. Endometrial_M and Ovarian_M specific mutated driver gene sets relative to each other

| Type | k | Specific gene set | p1 | p2 | q |
|---|---|---|---|---|---|
| **Endometrial_M / Ovarian_M** | 2 | *PIK3CA,PIK3R1* | <0.0001 | 0.892 | <0.0001 |
| | 3 | *PIK3CA,PIK3R1,AKT1* | <0.0001 | 0.895 | <0.0001 |
| | 4 | *PIK3CA,PIK3R1,AKT1,BCL2L11* | <0.0001 | 0.88 | <0.0001 |
| | 5 | *PIK3CA,PIK3R1,AKT1,BCL2L11,ERBB3* | <0.0001 | 0.873 | <0.0001 |
| | 6 | *PIK3CA,PIK3R1,AKT1,BCL2L11,ERBB3,ARID1B* | <0.0001 | 0.996 | <0.0001 |
| | 7 | *PIK3CA,PIK3R1,AKT1,BCL2L11,ERBB3,ARID1B,FOXO1* | <0.0001 | 0.989 | <0.0001 |
| | 8 | *PIK3CA,PIK3R1,AKT1,BCL2L11,ERBB3,ARID1B,FOXO1,CD79A* | <0.0001 | 0.991 | <0.0001 |
| | 9 | *PIK3CA,PIK3R1,AKT1,BCL2L11,ERBB3,ARID1B,FOXO1,GRB7,GRM4* | <0.0001 | 0.987 | <0.0001 |
| | 10 | *PIK3CA,PIK3R1,AKT1,BCL2L11,ERBB3,ARID1B,CD79A,GRB7,GRM4,SUFU* | <0.0001 | 0.993 | <0.0001 |
| **Ovarian_M / Endometrial_M** | 10 | *AXL,BAP1,CDK12,GRIN2A,JAK3,KDM5A,MSH2,NF1,PTPRS,TET2* | 0.025 | 1 | <0.0001 |

Table S 5.1.14. Significant common driver gene set between Breast_M, Endometrial_M and Ovarian_M

| **k** | **common gene set** | **p1** | **p2** | **p3** | **q** |
|---|---|---|---|---|---|
| 5 | *TP53,CDH1,CTNNB1,GATA3,KRAS* | <0.0001 | <0.0001 | <0.0001 | <0.0001 |
| 6 | *TP53,CDH1,CTNNB1,GATA3,KRAS,CDKN1B* | <0.0001 | <0.0001 | <0.0001 | <0.0001 |
| 7 | *TP53,CDH1,CTNNB1,GATA3,KRAS,CDKN1B,CDC73* | <0.0001 | <0.0001 | <0.0001 | <0.0001 |
| 8 | *TP53,CDH1,CTNNB1,GATA3,KRAS,CDKN1B,CDC73,RAD51C* | <0.0001 | <0.0001 | <0.0001 | <0.0001 |
| 9 | *TP53,CDH1,CTNNB1,GATA3,KRAS,MYOD1,CDC73,RAD51C,PRKAR1A* | <0.0001 | <0.0001 | <0.0001 | <0.0001 |
| 10 | *TP53,CDH1,CTNNB1,GATA3,KRAS,MYOD1,CDC73,RAD51C,PRKAR1A,TMPRSS2* | <0.0001 | <0.0001 | <0.0001 | <0.0001 |

#### 5.2 Male-Bladder, Prostate cancer and Renal cell carcinoma

*Primary cancers*

*There is no significant common driver gene set between Bladder_P and Prostate_P when k=2 ∼ 10.

Table S 5.2.1. Significant common driver gene set between Bladder_P and Renal_P

| k | common gene set | p1 | p2 | q |
|---|---|---|---|---|
| 5 | *AKT3,TERT,TFE3,VHL,PTCH1* | 0.034 | 0.015 | <0.0001 |
| 6 | *AKT3,TERT,TFE3,VHL,ATRX,NF2* | 0.024 | <0.0001 | <0.0001 |
| 7 | *AKT3,TERT,TFE3,VHL,ATRX,NF2,TAFA2* | 0.007 | 0.003 | <0.0001 |
| 8 | *AKT3,TERT,TFE3,VHL,ATRX,NF2,TNIP2,HCN1* | 0.005 | <0.0001 | <0.0001 |
| 9 | *AKT3,TERT,TFE3,VHL,ATRX,NF2,HIST1H3E,RAD52,VPS35* | 0.002 | 0.001 | <0.0001 |
| 10 | *AKT3,TERT,TFE3,VHL,ATRX,NF2,HIST1H3E,TNFRSF14,VCL,HCN1* | 0.002 | <0.0001 | <0.0001 |

Table S 5.2.2. Significant common driver gene set between Prostate_P and Renal_P

| k | common gene set | p1 | p2 | q |
|---|---|---|---|---|
| 5 | *FOXA1,NF2,SPOP,TMPRSS2,VHL* | <0.0001 | 0.016 | <0.0001 |
| 6 | *FOXA1,NF2,SPOP,TMPRSS2,VHL,PTEN* | <0.0001 | 0.009 | <0.0001 |
| 7 | *FOXA1,NF2,SPOP,TMPRSS2,VHL,PTEN,MET* | <0.0001 | <0.0001 | <0.0001 |
| 8 | *FOXA1,NF2,SPOP,TMPRSS2,VHL,PTEN,MET,KMT2A* | <0.0001 | <0.0001 | <0.0001 |
| 9 | *FOXA1,NF2,SPOP,TMPRSS2,VHL,PTEN,MET,KMT2A,BRAF* | <0.0001 | <0.0001 | <0.0001 |
| 10 | *FOXA1,NF2,SPOP,TMPRSS2,VHL,PTEN,MET,NOTCH4,BRAF,TFE3* | <0.0001 | <0.0001 | <0.0001 |

*There is no significant common driver gene set among Bladder_P, Prostate_P and Renal_P when k=2 ∼ 10.

Table S 5.2.3. Bladder_P and Prostate_P specific mutated driver gene sets relative to each other

| Type | k | Specific gene set | p1 | p2 | q |
|---|---|---|---|---|---|
| **Bladder_P / Prostate_P** | 3 | *ATRX,TERT,CTLA4* | <0.0001 | 0.983 | 0.002 |
| | 4 | *ATRX,TERT,CTLA4,FOXP1* | <0.0001 | 0.96 | 0.001 |
| | 5 | *ATRX,TERT,BAP1,STAG2,TGFBR1* | 0.01 | 0.995 | 0.002 |
| | 6 | *ATRX,TERT,BAP1,STAG2,TGFBR1,COP1* | 0.008 | 0.997 | 0.004 |
| | 7 | *ATRX,TERT,BAP1,STAG2,TGFBR1,COP1,SMAD3* | 0.002 | 0.996 | 0.001 |
| | 9 | *ATRX,TERT,COP1,ERCC5,FBXW7,KDM6A,NFE2L2,TGFBR1,AKT3* | 0.041 | 1 | 0.003 |
| | 10 | *ATRX,TERT,COP1,ERCC5,FBXW7,KDM6A,NFE2L2,TGFBR1,AKT3,BAP1* | 0.039 | 1 | 0.003 |
| **Prostate_P / Bladder_P** | 2 | *SPOP,TMPRSS2* | <0.0001 | 0.999 | <0.0001 |
| | 3 | *SPOP,TMPRSS2,FOXA1* | <0.0001 | 0.995 | <0.0001 |
| | 4 | *SPOP,TMPRSS2,FOXA1,MAP2K4* | <0.0001 | 1 | <0.0001 |
| | 5 | *SPOP,TMPRSS2,FOXA1,PTEN,PGR* | <0.0001 | 1 | <0.0001 |
| | 6 | *SPOP,TMPRSS2,FOXA1,PTEN,GSK3B,NKX2-1* | <0.0001 | 1 | <0.0001 |
| | 7 | *SPOP,TMPRSS2,FOXA1,PTEN,GSK3B,PGR,EIF1AX* | <0.0001 | 1 | <0.0001 |
| | 8 | *SPOP,TMPRSS2,FOXA1,PTEN,GSK3B,PGR,EIF1AX,NKX2-1* | <0.0001 | 1 | <0.0001 |
| | 9 | *SPOP,TMPRSS2,FOXA1,PTEN,GSK3B,PGR,EIF1AX,NKX2-1,SYF2* | <0.0001 | 1 | <0.0001 |
| | 10 | *SPOP,TMPRSS2,FOXA1,PTEN,GSK3B,PGR,EIF1AX,NKX2-1,SPPL3,ANTXR2* | <0.0001 | 1 | <0.0001 |

Table S 5.2.4. Bladder_P and Renal_P specific mutated driver gene sets relative to each other

| **Type** | **k** | **Specific gene set** | **p1** | **p2** | **q** |
|---|---|---|---|---|---|
| **Bladder_P / Renal_P** | 2 | *TERT,PTCH1* | 0.011 | 0.755 | 0.026 |
| | 3 | *TERT,ATRX,CTLA4* | 0.002 | 0.734 | 0.001 |
| | 4 | *TERT,CDKN2A,FOXP1,KRAS* | 0.003 | 1 | 0.001 |
| | 5 | *TP53,ERBB2,FGFR3,HRAS,KRAS* | <0.0001 | 0.951 | <0.0001 |
| | 6 | *TP53,ERBB2,FGFR3,HRAS,KRAS,ATR* | <0.0001 | 0.907 | <0.0001 |
| | 7 | *TP53,ERBB2,FGFR3,HRAS,KRAS,CDKN1A,CDKN2A* | <0.0001 | 0.952 | <0.0001 |
| | 8 | *TP53,ERBB2,FGFR3,HRAS,KRAS,CDKN1A,CDKN2A,BLM* | <0.0001 | 0.939 | <0.0001 |
| | 9 | *TP53,MAPK1,FGFR3,HRAS,KRAS,CDKN1A,IFNGR1,KDM6A,NFE2L2* | <0.0001 | 0.869 | <0.0001 |
| | 10 | *TP53,MAPK1,FGFR3,HRAS,KRAS,CDKN1A,IFNGR1,KDM6A,NFE2L2,YES1* | <0.0001 | 0.857 | <0.0001 |
| **Renal_P / Bladder_P** | 3 | *NF2,TFE3,VHL* | 0.001 | 0.92 | 0.002 |
| | 4 | *NF2,TFE3,VHL,ELOC* | <0.0001 | 0.923 | <0.0001 |
| | 5 | *KEAP1,NF2,PBRM1,RPS6KA4,SFPQ* | 0.011 | 1 | <0.0001 |
| | 6 | *KEAP1,PAK5,PBRM1,RPS6KA4,SFPQ,CDKN2B* | 0.046 | 1 | <0.0001 |
| | 8 | *KEAP1,PAK5,PBRM1,RPS6KA4,SFPQ,CDKN2B,PIK3CG,TBX3* | 0.049 | 1 | <0.0001 |
| | 9 | *KEAP1,PAK5,PBRM1,RPS6KA4,SFPQ,CDKN2B,PIK3CG,TBX3,TRAF2* | 0.032 | 1 | <0.0001 |
| | 10 | *KEAP1,PAK5,PBRM1,RPS6KA4,SFPQ,CDKN2B,PIK3CG,TBX3,PIM1,FH* | 0.015 | 1 | <0.0001 |

Table S 5.2.5. Prostate_P and Renal_P specific mutated driver gene sets relative to each other

| Type | k | Specific gene set | p1 | p2 | q |
|---|---|---|---|---|---|
| Prostate_P / Renal_P | 2 | *SPOP,TMPRSS2* | <0.0001 | 1 | <0.0001 |
| | 3 | *SPOP,TMPRSS2,FOXA1* | <0.0001 | 1 | <0.0001 |
| | 4 | *SPOP,TMPRSS2,FOXA1,CDK12* | <0.0001 | 1 | <0.0001 |
| | 5 | *APC,BRCA2,CDK12,CTNNB1,ERG* | 0.004 | 0.97 | <0.0001 |
| | 8 | *TP53,APC,BRCA2,CDK12,CTNNB1,ERG,PTPRD,ZFHX3* | 0.048 | 0.861 | 0.045 |
| | 9 | *TP53,APC,BRCA2,CDK12,CTNNB1,ERG,PTPRD,ZFHX3,CUL3* | 0.021 | 0.988 | 0.012 |
| | 10 | *TP53,APC,BRCA2,CDK12,CTNNB1,ERG,PTPRD,ZFHX3,CUL3,IDH1* | 0.01 | 0.989 | 0.003 |
| Renal_P / Prostate_P | 2 | *NF2,VHL* | 0.008 | 1 | 0.007 |
| | 3 | *NF2,VHL,TFE3* | 0.001 | 1 | <0.0001 |
| | 4 | *NF2,VHL,TERT,TSC2* | 0.004 | 0.998 | 0.002 |

#### *Metastatic cancers*

Table S 5.2.6. Significant common driver gene set between Bladder_M and Prostate_M

| k | common gene set | p1 | p2 | q |
|---|---|---|---|---|
| 4 | *TP53,CDK12,FGFR3,BRCA2* | 0.026 | <0.0001 | <0.0001 |
| 5 | *TP53,CDK12,FGFR3,BRCA2,PIK3R1* | 0.039 | <0.0001 | <0.0001 |
| 6 | *TP53,CDK12,FGFR3,BRD4,APC,FLT4* | 0.018 | <0.0001 | <0.0001 |
| 7 | *TP53,CDK12,FGFR3,BRD4,APC,DICER1,FLT4* | 0.025 | <0.0001 | <0.0001 |
| 8 | *TP53,CDK12,FGFR3,BRD4,APC,DICER1,FLT4,FGFR1* | 0.01 | <0.0001 | <0.0001 |
| 9 | *TP53,CDK12,FGFR3,BRD4,APC,DICER1,FLT4,FGFR1,CSF1R* | 0.01 | <0.0001 | <0.0001 |
| 10 | *TP53,CDK12,FGFR3,BRD4,APC,DICER1,FLT3,CHEK2,LINC00114,SMO* | 0.002 | <0.0001 | <0.0001 |

Table S 5.2.7. Significant common driver gene set between Bladder_M and Renal_M

| k | common gene set | p1 | p2 | q |
|---|---|---|---|---|
| 5 | *KRAS,NF2,TERT,TSC2,VHL* | 0.004 | <0.0001 | <0.0001 |
| 6 | *KRAS,NF2,TERT,TSC2,VHL,TFE3* | 0.009 | <0.0001 | <0.0001 |
| 7 | *KRAS,NF2,TERT,TSC2,VHL,TFE3,SMAD4* | <0.0001 | <0.0001 | <0.0001 |
| 8 | *KRAS,NF2,TERT,TSC2,VHL,TFE3,SMAD4,SMAD3* | <0.0001 | <0.0001 | <0.0001 |
| 9 | *KRAS,NF2,TERT,TSC2,VHL,TFE3,SMAD4,GPS2,ALOX12B* | <0.0001 | <0.0001 | <0.0001 |
| 10 | *KRAS,NF2,TERT,TSC2,VHL,TFE3,SMAD4,GPS2,SMAD3,NRAS* | <0.0001 | <0.0001 | <0.0001 |

Table S 5.2.8. Significant common driver gene set between Prostate_M and Renal_M

| k | common gene set | p1 | p2 | q |
|---|---|---|---|---|
| 3 | *TP53,CDK12,VHL* | <0.0001 | 0.004 | <0.0001 |
| 4 | *TP53,CDK12,VHL,APC* | <0.0001 | 0.002 | <0.0001 |
| 5 | *TP53,CDK12,VHL,APC,NF2* | <0.0001 | <0.0001 | <0.0001 |
| 6 | *TP53,CDK12,VHL,APC,NF2,KDM6A* | <0.0001 | <0.0001 | 0.002 |
| 7 | *TP53,CDK12,VHL,APC,NF2,KDM6A,FGFR3* | <0.0001 | <0.0001 | <0.0001 |
| 8 | *TP53,CDK12,VHL,APC,NF2,KDM6A,FGFR3,MEN1* | <0.0001 | 0.001 | <0.0001 |
| 9 | *TP53,CDK12,VHL,APC,NF2,KDM6A,FGFR3,MEN1,TFE3* | <0.0001 | <0.0001 | <0.0001 |
| 10 | *TP53,CDK12,VHL,TERT,NF2,KDM6A,ERBB3,SPOP,TFE3,BRCA2* | <0.0001 | <0.0001 | <0.0001 |

Table S 5.2.9. Bladder_M and Prostate_M specific mutated driver gene sets relative to each other

| Type | k | Specific gene set | p1 | p2 | q |
|---|---|---|---|---|---|
| Bladder_M / Prostate_M | 3 | *TERT,NRAS,AKT2* | 0.005 | 0.987 | 0.001 |
| | 5 | *TERT,NRAS,CDKN1A,KRAS,RBM10* | 0.011 | 0.997 | 0.004 |
| | 6 | *TERT,NRAS,CDKN1A,KRAS,RBM10,H3F3A* | 0.003 | 0.993 | <0.0001 |
| | 7 | *TERT,NRAS,CDKN1A,KRAS,RBM10,H3F3A,VTCN1* | 0.003 | 0.996 | 0.002 |
| | 8 | *TERT,NRAS,CDKN1A,KRAS,RBM10,H3F3A,VTCN1,ALOX12B* | 0.001 | 0.993 | <0.0001 |
| | 9 | *TERT,NRAS,CDKN1A,KRAS,RBM10,H3F3A,VTCN1,ALOX12B,SRC* | <0.0001 | 0.998 | <0.0001 |
| | 10 | *TERT,NRAS,CDKN1A,KRAS,RBM10,H3F3A,VTCN1,ALOX12B,SRC,SRSF2* | <0.0001 | 0.988 | <0.0001 |
| Prostate_M / Bladder_M | 2 | *FOXA1,TMPRSS2* | 0.01 | 1 | 0.014 |
| | 3 | *FOXA1,TMPRSS2,AR* | <0.0001 | 0.91 | <0.0001 |
| | 4 | *FOXA1,TMPRSS2,PTEN,SPOP* | 0.002 | 0.873 | 0.001 |
| | 5 | *FOXA1,TMPRSS2,PTEN,SPOP,AR* | 0.011 | 0.719 | 0.008 |
| | 6 | *FOXA1,TMPRSS2,PTEN,SPOP,AR,PIK3R1* | 0.018 | 0.705 | 0.015 |
| | 7 | *FOXA1,TMPRSS2,PTEN,SPOP,AR,PIK3R1,CDKN1B* | 0.009 | 0.721 | 0.012 |
| | 8 | *FOXA1,TMPRSS2,PTEN,SPOP,AR,PIK3R1,CDKN1B,MALT1* | 0.005 | 0.698 | 0.003 |
| | 9 | *FOXA1,TMPRSS2,PTEN,SPOP,AR,PIK3R1,CDKN1B,MALT1,INPP4B* | 0.003 | 0.721 | 0.002 |
| | 10 | *FOXA1,TMPRSS2,PTEN,SPOP,AR,PIK3R1,CDKN1B,MALT1,INPP4B,BRAF* | 0.007 | 0.913 | 0.01 |

Table S 5.2.10. Bladder_M and Renal_M specific mutated driver gene sets relative to each other

| Type | k | Specific gene set | p1 | p2 | q |
|---|---|---|---|---|---|
| Bladder_M / Renal_M | 2 | *KRAS,TERT* | 0.006 | 0.992 | 0.005 |
| | 3 | *KRAS,TERT,SMAD3* | 0.003 | 0.992 | <0.0001 |
| | 4 | *KRAS,TERT,KDM6A,NRAS* | 0.014 | 0.908 | 0.006 |
| | 6 | *CDKN1A,ERBB3,KDM6A,RB1,PTPRD,KMT2D* | 0.013 | 0.988 | 0.007 |
| | 7 | *CDKN1A,ERBB3,KDM6A,RB1,RPTOR,CREBBP,INSR* | 0.004 | 1 | 0.001 |
| | 8 | *CDKN1A,ERBB3,KDM6A,RB1,RPTOR,CREBBP,INSR,KDM5A* | 0.001 | 1 | <0.0001 |
| | 9 | *CDKN1A,ERBB3,KDM6A,RB1,RPTOR,CREBBP,INSR,KDM5A,ERCC2* | 0.002 | 1 | <0.0001 |
| | 10 | *CDKN1A,ERBB3,KDM6A,RB1,RPTOR,CREBBP,INSR,KDM5A,ERCC2,EWSR1* | 0.002 | 1 | <0.0001 |
| Renal_M / Bladder_M | 2 | *NF2,VHL* | 0.001 | 0.928 | 0.004 |
| | 3 | *NF2,VHL,MET* | <0.0001 | 0.947 | <0.0001 |
| | 4 | *NF2,VHL,MET,TFE3* | <0.0001 | 0.946 | <0.0001 |

Table S 5.2.11. Prostate_M and Renal_M specific mutated driver gene sets relative to each other

| Type | k | Specific gene set | p1 | p2 | q |
|---|---|---|---|---|---|
| **Prostate_M / Renal_M** | 2 | *FOXA1,TMPRSS2* | 0.008 | 1 | 0.018 |
| | 3 | *FOXA1,TMPRSS2,AR* | 0.001 | 1 | 0.001 |
| | 4 | *FOXA1,TMPRSS2,AR,SPOP* | <0.0001 | 1 | <0.0001 |
| | 5 | *FOXA1,TMPRSS2,AR,SPOP,KDM6A* | <0.0001 | 0.983 | 0.001 |
| | 6 | *FOXA1,TMPRSS2,AR,SPOP,KMT2C,EPHA3* | <0.0001 | 0.992 | <0.0001 |
| | 7 | *FOXA1,TMPRSS2,AR,SPOP,KMT2C,EPHA3,EPHA5* | <0.0001 | 0.997 | <0.0001 |
| | 8 | *FOXA1,TMPRSS2,AR,SPOP,KMT2C,EPHA3,EPHA5,NCOR1* | <0.0001 | 1 | <0.0001 |
| | 9 | *FOXA1,TMPRSS2,AR,SPOP,KMT2C,EPHA3,EPHA5,NCOR1,BRCA2* | <0.0001 | 0.999 | <0.0001 |
| | 10 | *FOXA1,TMPRSS2,AR,KMT2D,KMT2C,EPHA3,EPHA5,NCOR1,EPHB1,RPTOR* | <0.0001 | 1 | <0.0001 |
| **Renal_M / Prostate_M** | 2 | *NF2,VHL* | 0.007 | 1 | 0.002 |
| | 3 | *BAP1,PBRM1,TERT* | 0.007 | 0.987 | 0.001 |
| | 4 | *BAP1,PBRM1,TERT,NF2* | 0.001 | 0.992 | <0.0001 |
| | 5 | *BAP1,PBRM1,TERT,SMARCB1,KDM5C* | 0.003 | 0.996 | <0.0001 |
| | 6 | *BAP1,PBRM1,TERT,SMARCB1,TSC1,SETD2* | 0.042 | 0.998 | 0.016 |
| | 7 | *BAP1,PBRM1,TERT,SMARCB1,TSC1,SETD2,IL10* | 0.011 | 0.987 | 0.007 |
| | 8 | *BAP1,PBRM1,TERT,SMARCB1,TSC1,SETD2,IL10,TFE3* | 0.01 | 0.99 | 0.003 |
| | 9 | *BAP1,PBRM1,TERT,SMARCB1,TSC1,SETD2,IL10,TSC2,DNMT1* | 0.002 | 1 | <0.0001 |
| | 10 | *BAP1,PBRM1,MET,SMARCB1,TSC1,SETD2,IL10,TSC2,DNMT1,INPP4A* | 0.042 | 1 | 0.001 |

Table S 5.2.12. Significant common driver gene set among Bladder_M, Prostate_M and Renal_M

| **k** | **common gene set** | **p1** | **p2** | **p3** | **q** |
|---|---|---|---|---|---|
| 4 | *TP53,CDK12,FGFR3,VHL* | 0.005 | <0.0001 | 0.005 | <0.0001 |
| 5 | *TP53,CDK12,FGFR3,VHL,BRCA2* | 0.005 | <0.0001 | 0.008 | <0.0001 |
| 6 | *TP53,CDK12,FGFR3,VHL,BRCA2,NF2* | 0.031 | <0.0001 | <0.0001 | <0.0001 |
| 7 | *TP53,CDK12,FGFR3,VHL,BRCA2,MET,TFE3* | 0.006 | <0.0001 | <0.0001 | <0.0001 |
| 8 | *TP53,CDK12,FGFR3,VHL,BRCA2,MET,TFE3,NF2* | 0.029 | <0.0001 | <0.0001 | <0.0001 |
| 9 | *TP53,CDK12,FGFR3,VHL,BRCA2,MET,TFE3,DIS3,PPM1D* | 0.004 | <0.0001 | <0.0001 | <0.0001 |
| 10 | *TP53,CDK12,FGFR3,VHL,BRCA2,MET,TFE3,DIS3,PPM1D,LINC00114* | 0.002 | <0.0001 | <0.0001 | <0.0001 |

#### 5.3 Esophagogastric & Gastrointestinal Stromal tumor

*Primary cancers*

Table S 5.3.1. Significant common driver gene set between Esophagogastric_P and GastrointestinalStromal_P

| k | common gene set | p1 | p2 | p3 | q |
|---|---|---|---|---|---|
| 4 | *TP53,CDK12,FGFR3,VHL* | 0.005 | <0.0001 | 0.005 | <0.0001 |
| 5 | *TP53,CDK12,FGFR3,VHL,BRCA2* | 0.005 | <0.0001 | 0.008 | <0.0001 |
| 6 | *TP53,CDK12,FGFR3,VHL,BRCA2,NF2* | 0.031 | <0.0001 | <0.0001 | <0.0001 |
| 7 | *TP53,CDK12,FGFR3,VHL,BRCA2,MET,TFE3* | 0.006 | <0.0001 | <0.0001 | <0.0001 |
| 8 | *TP53,CDK12,FGFR3,VHL,BRCA2,MET,TFE3,NF2* | 0.029 | <0.0001 | <0.0001 | <0.0001 |
| 9 | *TP53,CDK12,FGFR3,VHL,BRCA2,MET,TFE3,DIS3,PPM1D* | 0.004 | <0.0001 | <0.0001 | <0.0001 |
| 10 | *TP53,CDK12,FGFR3,VHL,BRCA2,MET,TFE3,DIS3,PPM1D,LINC00114* | 0.002 | <0.0001 | <0.0001 | <0.0001 |

*Esophagogastric_P has no specific mutated driver gene sets relative to GastrointestinalStromal_P when k=2 $\sim$ 10.
*GastrointestinalStromal_P has no specific mutated driver gene sets relative to Esophagogastric_P when k=2 $\sim$ 10.

#### *Metastatic cancers*

Table S 5.3.2. Significant common driver gene set between Esophagogastric_M and GastrointestinalStromal_M

| k | common gene set | p1 | p2 | q |
|---|---|---|---|---|
| 2 | *TP53,BCOR* | 0.018 | 0.016 | 0.001 |
| 3 | *TP53,BCOR,TGFBR1* | <0.0001 | 0.001 | <0.0001 |
| 4 | *TP53,BCOR,TGFBR1,STK11* | <0.0001 | <0.0001 | <0.0001 |
| 5 | *TP53,BCOR,TGFBR1,FGFR2,TGFBR2* | <0.0001 | <0.0001 | <0.0001 |
| 6 | *TP53,BCOR,TGFBR1,FGFR2,TGFBR2,BRCA1* | <0.0001 | <0.0001 | <0.0001 |
| 7 | *TP53,BCOR,TGFBR1,FGFR2,TGFBR2,NTRK3,PRKAR1A* | <0.0001 | <0.0001 | <0.0001 |
| 8 | *TP53,BCOR,TGFBR1,FGFR2,CCDC6,STK11,PRKAR1A,CMIP* | <0.0001 | <0.0001 | <0.0001 |
| 9 | *TP53,BCOR,TGFBR1,FGFR2,NTRK3,BRCA1,PRKAR1A,CMIP,REL* | <0.0001 | <0.0001 | <0.0001 |
| 10 | *TP53,BCOR,TGFBR1,FGFR2,NTRK3,BRCA1,CCDC6,CMIP,HGF,STK11* | <0.0001 | <0.0001 | <0.0001 |

*Esophagogastric_M has no specific mutated driver gene sets relative to GastrointestinalStromal_M when k=2 $\sim$ 10.
*GastrointestinalStromal_M has no specific mutated driver gene sets relative to Esophagogastric_M when k=2 $\sim$ 10.

#### 5.4 melamona & skin cancer

*Primary cancers* *There is no significant common driver gene set between Melamona_P and Skin_P when k=2 ∼ 10.

Table S 5.4.1. Melamona_P specific mutated driver gene sets relative to Skin_P

| k | Specific gene set | p1 | p2 | q |
|---|---|---|---|---|
| 4 | *BRAF,GNA11,GNAQ,NRAS* | 0.001 | 0.927 | 0.006 |
| 5 | *BRAF,GNA11,GNAQ,NRAS,RAC1* | 0.003 | 1 | 0.001 |
| 6 | *BRAF,GNA11,GNAQ,NRAS,KIT,GATA1* | <0.0001 | 0.998 | <0.0001 |
| 7 | *BRAF,GNA11,GNAQ,NRAS,KIT,GATA1,IRS1* | <0.0001 | 1 | <0.0001 |
| 8 | *BRAF,GNA11,GNAQ,NRAS,KIT,GATA1,IRS1,B2M* | <0.0001 | 1 | <0.0001 |
| 9 | *BRAF,GNA11,GNAQ,NRAS,KIT,GATA1,ETV6,B2M,PDGFRA* | <0.0001 | 1 | <0.0001 |
| 10 | *BRAF,GNA11,GNAQ,NRAS,KIT,GATA1,IRS1,B2M,PDGFRA,PAK1* | <0.0001 | 1 | <0.0001 |

*Skin_P has no specific mutated driver gene sets relative to Melamona_P when k=2 ∼ 10.

*Metastatic cancers* *There is no significant common driver gene set between Melamona_M and Skin_M when k=2 ∼ 10.

Table S 5.4.2. Melamona_M and Skin_M specific mutated driver gene sets relative to each other

| **Type** | **k** | **Specific gene set** | **p1** | **p2** | **q** |
|---|---|---|---|---|---|
| **Melamona_M / Skin_M** | 2 | *BRAF,NRAS* | <0.0001 | 0.885 | <0.0001 |
| | 3 | *BRAF,NRAS,GNAQ* | <0.0001 | 0.88 | <0.0001 |
| | 4 | *BRAF,NRAS,GNAQ,NF1* | <0.0001 | 0.97 | <0.0001 |
| | 5 | *BRAF,NRAS,GNAQ,NF1,GNA11* | <0.0001 | 0.967 | <0.0001 |
| | 6 | *BRAF,NRAS,GNAQ,NF1,GNA11,KIT* | <0.0001 | 0.997 | <0.0001 |
| | 7 | *BRAF,NRAS,GNAQ,NF1,GNA11,KIT,MAP2K1* | <0.0001 | 1 | <0.0001 |
| | 8 | *BRAF,NRAS,GNAQ,NF1,GNA11,KIT,MAP2K1,SDHA* | <0.0001 | 1 | <0.0001 |
| | 9 | *BRAF,NRAS,GNAQ,NF1,GNA11,KIT,MAP2K1,SDHA,PIM1* | <0.0001 | 1 | <0.0001 |
| | 10 | *BRAF,NRAS,GNAQ,NF1,GNA11,KIT,MAP2K1,SDHA,PIM1,BCL2L14* | <0.0001 | 1 | <0.0001 |
| **Skin_M / Melamona_M** | 7 | *HOXB13,IRS1,MAX,MYOD1,SMARCB1,STK11,TRAF7* | 0.008 | 1 | <0.0001 |
| | 8 | *HOXB13,IRS1,MAX,MYOD1,SMARCB1,STK11,TRAF7,MYD88* | 0.005 | 1 | <0.0001 |
| | 9 | *HOXB13,IRS1,MAX,MYOD1,SMARCB1,STK11,TRAF7,MYD88,BBC3* | 0.004 | 1 | <0.0001 |
| | 10 | *HOXB13,IRS1,MAX,MYOD1,SMARCB1,STK11,TRAF7,MYD88,BBC3,NFKBIA* | 0.002 | 1 | <0.0001 |
